# A 4-lineage statistical suite to evaluate the support of large-scale retrotransposon insertion data to reconstruct evolutionary trees

**DOI:** 10.1101/2020.12.10.419374

**Authors:** G Churakov, A Kuritzin, K Chukharev, F Zhang, F Wünnemann, V Ulyantsev, J Schmitz

## Abstract

Retrophylogenomics makes use of genome-wide retrotransposon presence/absence insertion patterns to resolve questions in phylogeny and population genetics. In the genomics era, evaluating high-throughput data requires the associated development of appropriately powerful statistical tools. The currently used KKSC 3-lineage statistical test for estimating the significance of retrophylogenomic data is limited by the number of possible tree topologies it can assess in one step. To improve on this, we have extended the analysis to simultaneously compare 4-lineages, enabling us to evaluate ten distinct presence/absence insertion patterns for 26 possible tree topologies plus 129 trees with different incidences of hybridization or introgression. The new tool provides statistics for cases involving multiple ancestral hybridizations/introgressions, ancestral incomplete lineage sorting, bifurcation, and polytomy. The test is embedded in a user-friendly web R-application (http://retrogenomics.uni-muenster.de:3838/hammlet/) and is available for use by the scientific community.

## Introduction

Retrotransposon phylogenetic presence/absence markers are known to be virtually homoplasy-free (Doronina et al. 2019). Like other sequence alignments, they require careful manual inspections to be certain of the true orthology of their loci and evaluations of the potential tree topology. Statistical tests evaluate whether the number of shared insertions inherited from a common ancestor compared with the absence of such elements in other species is significant to support their relatedness. However, occasionally, presence/absence markers appear to support contradictory phylogenetic tree topologies. Such potential conflicts may be evoked by (1) extremely rare cases of parallel retrotransposon insertions in unrelated species (about 0.01% of diagnostic markers), (2) even rarer exact deletions in one or more related species (about 0.001% of diagnostic markers), or (3) most influentially, ancestral hybridization/introgression and incomplete lineage sorting (ILS), the extents of which can differ for diverse taxonomic groups (Doronina et al. 2019). ILS is mainly associated with the evolution of species on short internodal phylogenetic branches. These short branches are indicative of short periods between speciation, too short in fact for all inserted retrotransposons to have been fixed in all related representatives of a particular group before the next speciation occurred. Thus, some members of the group have the insertion and others do not, even though they still belong to the same phylogenetic group (Kuritzin et al. 2016).

Waddell et al. (2001) developed the first statistical test to evaluate the probability of a particular presence/absence marker insertion pattern supporting a prior hypothesis of relatedness against polytomy. However, the Waddell test returns p-values for only up to 5 marker combinations and is therefore no longer suitable for present-day genome analysis with hundreds or even thousands of markers. Kuritzin et al. (2016) developed the 3-lineage KKSC statistics, which introduced a multi-directional test that can evaluate the presence/absence patterns of phylogenetic markers represented in all potential tree topologies of three species without a prior hypothesis. KKSC also includes a less powerful one-directional test, in which the phylogenetic markers are screened from one reference lineage only and compared to the presence/absence patterns in other species (http://retrogenomics.uni-muenster.de:3838/KKSC_significance_test/). KKSC also includes a simple consideration of ancestral hybridization/introgression based on the symmetric or asymmetric distribution of phylogenetic markers for 3 potential concurring tree topologies involving 3 species/lineages. The KKSC statistics processes three empirical values to directly calculate p-values under binomial distribution criteria for selecting a tree against hybridization, a tree against polytomy or hybridization against polytomy (Kuritzin et al. 2016). However, the calculation is limited to evaluating the evidence for the interrelatedness of only 3 lineages and their seven corresponding potential tree topologies. Furthermore, they insist on a combination of sets of 3 taxa to evaluate larger trees. By changing to a likelihood maximization and likelihood ratio calculation and expanding the evaluation to handle 4 lineages, thereby testing 155 different tree topologies, 129 of which include varying extents of multiple hybridization/introgression scenarios, most phylogenetic questions can be accessed directly or in combining of 4 species sets. The 4-lineage (4-LIN) test requires multi-directional screening for phylogenetic markers starting from four different reference species and considers 10 distinct informative presence/absence patterns for each phylogenetic marker.

## Methods

Following our previous investigation (Kuritzin et al. 2016), we extended the diffusion approximation of Kimura (1955a; 1955b) to four lineages (Doronina et al. 2017a). Fisher (1922) and Wright (1933) first introduced the diffusion approximation of random genetic drift to population genetics. The diffusion approximation transfers a complex stochastic distribution to an appropriate and mathematically more tractable diffusion Markov process (Glynn 1990). The approximation enables us to evaluate a large number of branching processes. Based on presence/absence patterns of inserted retrotransposons, we developed new statistical criteria depending on likelihood ratio calculations to identify the most likely supported phylogenetic tree from all possible 155 four-lineage topologies that include multiple hybridization/introgression scenarios, binary trees, and partial or fully unsolved trees. To apply the 4-LIN statistical test, retrotransposon presence/absence patterns must first be derived from species representing four lineages. To enable an almost unbiased screening, we recommend screening qualitatively equal genome assemblies (covering equal percentages of the genomes). There is no limit to the number of phylogenetic markers that can be evaluated.

We derived a mathematical model describing the behavior of retrotransposon insertions in populations over time of speciation for four lineages. We then selected five variable parameters (n_0_, T_1_, T_3_, γ_1_, γ_3_), representing the effective population size coefficient (n_0_), the length of the first branch (T_1_), the length of the second branch (T_3_), the gamma value for a potential first hybridization (γ_1_), and the gamma value of a possible second hybridization (γ_3_). By applying permutations to the parameter set, fixing numbers of parameters, and excluding all symmetric tree variants, we defined 155 individual trees (see Appendix 1, Table 2, and below) composed of five groups (groups Ѳ4, Ѳ3, Ѳ2, Ѳ1, Ѳ0, see below). We have compiled the frequencies within the analyzed data ordered into 10 possible presence/absence patterns (-+++, --++, -+-+, -++-, +-++, +--+, +-+-, ++-+, ++--, +++-with the absence of a retrotransposon (-) or its presence (+) at an orthologous genomic locus in the four species/lineages). A simplified pattern (0, 0, 0, 100, 0, 0, 0, 0, 0, 0) inserted into the 4-LIN statistical suite (http://retrogenomics.uni-muenster.de:3838/hammlet/) reveals a highly significant affiliation of two of the species/lineages. The probability of the most likely topology among the predefined 155 trees is derived after using likelihood ratio tests based on up to five variable parameters and five groups (Ѳ4, Ѳ3, Ѳ2, Ѳ1, Ѳ0, see below) with different amounts of variable parameters between them.

### Model assumptions

Let’s consider four lineages *A*_*1*_, *A*_*2*_, *A*_*3*_, *A*_*4*_ sharing common ancestry. We can describe each relevant branch *B*_*1*_, *B*_*2*_, *B*_*3*_, *B*_*4*_ (see Fig. 1) as an isolated population of individuals over an evolutionary time interval *t* ∈ {Δ_*0*_, Δ_*1*_} with an effective population size *N*_*k*_(*t*) (where 1≤k≤4, indicating an index of respective branch B, Δ_*0*_ indicates the interval of the ancestral branch, and Δ_*k*_ indicates the interval of the respected branch *B*_*k*_). We use the fusion model to record hybridization/introgression, whereby two separated ancestral populations reproduce to form a new branch (Kuritzin et al. 2016). It should be noted that both hybridization and introgression can be described in terms of the fusion model but cannot be distinguished by experimental presence/absence pattern data. We can consider two possible basic models, 2H1 and 2H2, of speciation for four lineages, including each of two potential hybridization/introgression events (Fig. 1a,b).

**Figure 1.**
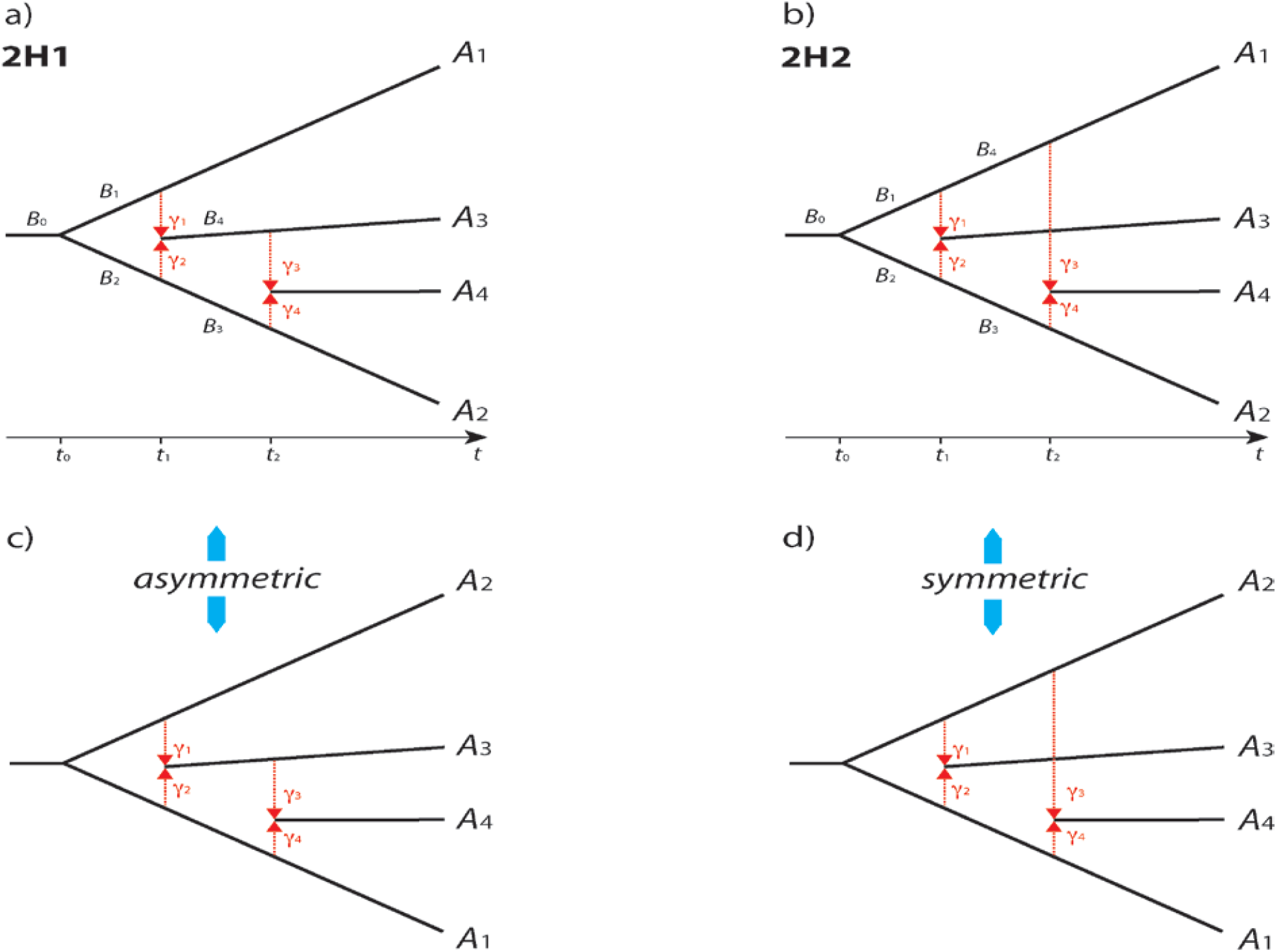
Two basic evolutionary models involving the four lineages *A*_*1*_, *A*_*2*_, *A*_*3*_, *A*_*4*_. Asymmetric and symmetric permutations. Black lines indicate different lineages, and red vertical dotted arrows indicate hybridization/introgression events. A) Basic model 2H1: sequential hybridization/introgression, B) Basic model 2H2: parallel hybridization/introgression. The proportions of two fused subpopulations forming a new population at time *t* = *t*_*1*_ are denoted *γ*_*1*_ and *γ*_*2*,_ where *γ*_*1*_ + *γ*_*2*_ = 1. At the second fusion point (*t* = *t*_*2*_), the sub-population proportions are denoted *γ*_*3*_ and *γ*_*4*_ where *γ*_*3*_ + *γ*_*4*_ = 1. C) Asymmetric permutation of model 2H1. Permutations of A_1_, A_2_, A_3_, A_4,_ and A_2_, A_1_, A_3_, A_4_ lead to different trees. Lineage A_3_ results from hybridization/introgression between lineages A_1_ and A_2_ for both the upper and lower permutations, respectively. However, lineage A_4_ is the result of the hybridization/introgression of A_2_ and A_3_ lineages in the upper tree and lineages A_1_ and A_3_ in the lower tree. B) The symmetric permutation of model 2H2 leads to identical phylogenetic trees (A_1_, A_2_, A_3_, A_4_ and A_2_, A_1_, A_3_, A_4_), taking into account the exchange of the value of *γ*_1_ by *γ*_2_ and *γ*_3_ by *γ*_4_.

For each of these two hybridization/introgression scenarios, we can derive 24 permutations of the four lineages *A*_*1*_, *A*_*2*_, *A*_*3*_, *A*_*4*_. However, due to symmetry (see Fig. 1d) the number of rearrangements for 2H2 can be reduced to 12 permutations (see below).

For a neutral insertion of retroelements, in generation *t* of the branch *B*_*k*_, we consider ten different events ω_*i,j*_ (1 ≤ *i* ≤ *j* ≤ 4) for four lineages where:

> in ω_*i,i*_ a retroelement is absent in the orthologous locus of lineage *A*_*i*_ but present in the other three lineages;
>
> in ω_*i,j*_ a retroelement is absent in the orthologous loci of lineages *A*_*i*_ and *A*_*j*_ (*i* ≠ *j*) but present in the other two lineages.

Assuming that the probability of new insertions for each individual of a population is small and the effective population size is large, we can conclude that the total number of retroelement insertions with the property ω_*i,j*_ are independent, assumedly Poisson-distributed, random variables with parameters *a*_*i,j*_.

We analyzed these models to derive formulas for *a*_*i,j*_ (see Appendix 1, S1.3.1.-S1.3.30). The calculations were carried out similarly to those in Kuritzin et al. (2016) under the assumption that the effective population size functions *N*_*k*_(*t*) at the corresponding intervals are constant (then *n*_*k*_ indicates the efficient population size on the branch *k*). Hence, denoting 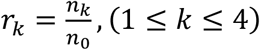 and fixing *r*_*k*_ = 1, we can write *a*_*i,j*_ = *n*_0_*ā*_*i,j*_, with parametric function *ā*_*i,j*_ depending on the four values *T*_1,_ *T*_3_, *γ*_1,_ and *γ*_3_ (noting that *γ*_2_ = *1* − *γ*_*1*_; and *γ*_4_ = *1* − *γ*_3_), where, 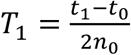, *and* 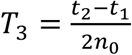, reflecting the periods from the first split of two branches from an ancestral population to the first hybridization/introgression event, and from the first hybridization/introgression event to the second hybridization/introgression event, where *n*_0_ is the parameter directly proportional to the effective population size (see Appendix 1, S1.4.1-S1.4.10).

### Model investigations

For modeling different tree variants, we need to fix the model parameters *T*_1_, *T*_3_, *γ*_1_, *γ*_3_ to their boundaries. It should be mentioned that fixing *T*_*1*_ and *T*_*3*_ to the boundary may lead to branch lengths of zero (*T*_*1*_=0 and *T*_3_=0) and that *γ*_1_ and *γ*_3_ can be fixed to the two extremes *γ*=0 or *γ*=1, leading to different models. When a branch is of zero length (*T*=0), in most cases, *γ* is undefined (changing *γ* does not change the model).

For further discussion, we introduce the “TTgg” nomenclature of models. For example, the 2H1 model we annotate as H1:TTgg, where each T or g reflects the status of a parameter: the first “T” reflects T_1_, the second “T” corresponds to T_3_. The first “g” shows the status of *γ*_1_, and the second “g” represents *γ*_3_; the prefix (H1 or H2) refers to the basic models (2H1 or 2H2).

To switch between different models, we can vary specific parameters, e.g., adjusting *γ*_3_ to 0 will switch H1:TTgg (2H1 model with double hybridization/introgression; see Appendix 1, Table 1) to H1:TTg0 (1H1 model). Or, adjusting *γ*_3_ to 1 will switch the same model to H1:TTg1 (1H4 model).

From the model 2H1, which has 24 different permutations of its four lineages, we can create 20 different models (each with 24 permutations). From model 2H2, which has 12 permutations, we can derive 22 different models, each with 12 permutations (see Appendix 1, Table 1). Thus, we can derive a total of 744 variants (24 × 20 + 12 × 22) of trees (models with fixed orders of species; e.g., 2H1:1234 (H1:TTgg:1234)). Excluding symmetric options (see Fig. 1c,d) and duplicated models with identical biological meaning reduce this to a set of 15 models and 155 unique trees (see Appendix 1, Table 2 for a non-redundant list of model variants and permutations).

Assuming models 2H1 and 2H2 (Fig. 1a,b), we can sort diagnostic cases into five groups from complex to simple, depending on the number of fixed parameters (Figs. 2, 3, 4, 5).

**Figure 2.**
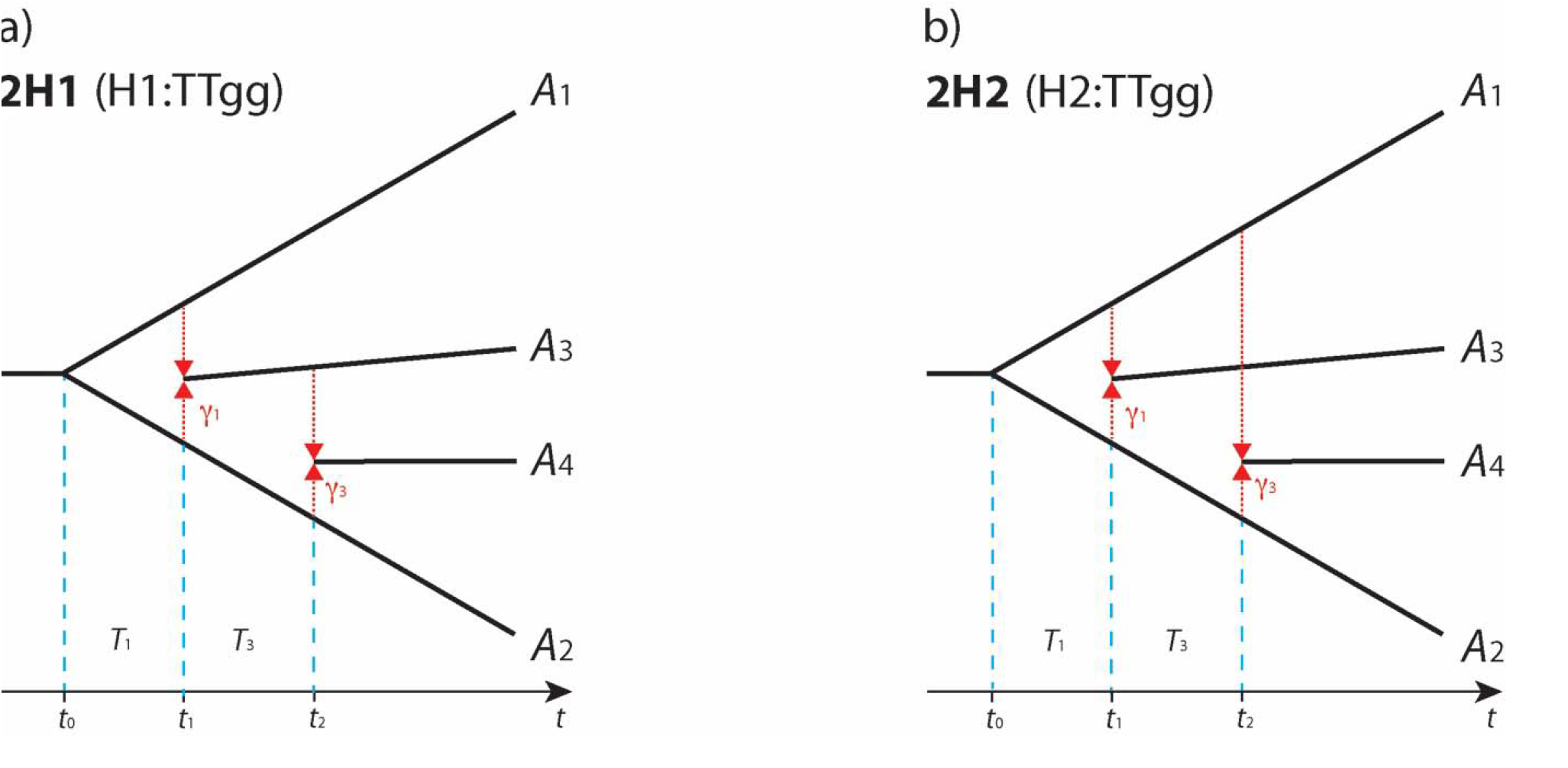
Group θ4. Black arrows below the trees indicate the times (t) of splits with events *t*_*0*_, *t*_*1*_, *t*_*2*_. Here, *t*_*0*_ denotes the initial split, *t*_*1*_ marks the first hybridization/introgression event (red arrows), and *t*_*2*_ the second (red arrows). Model names (2H1, 2H2) and aliases (H1:TTgg, H2:TTgg) are above the trees. A_1_, A_2_, A_3_, A_4_ denote the order of lineages in the first permutation (1234), and *T*_*1*_ and *T*_*3*_ indicate different periods, while *γ*_1_, and *γ*_3_ are the hybridization/introgression coefficients. a) two consecutive hybridizations/introgressions, b) two parallel hybridizations/introgressions.

**Figure 3.**
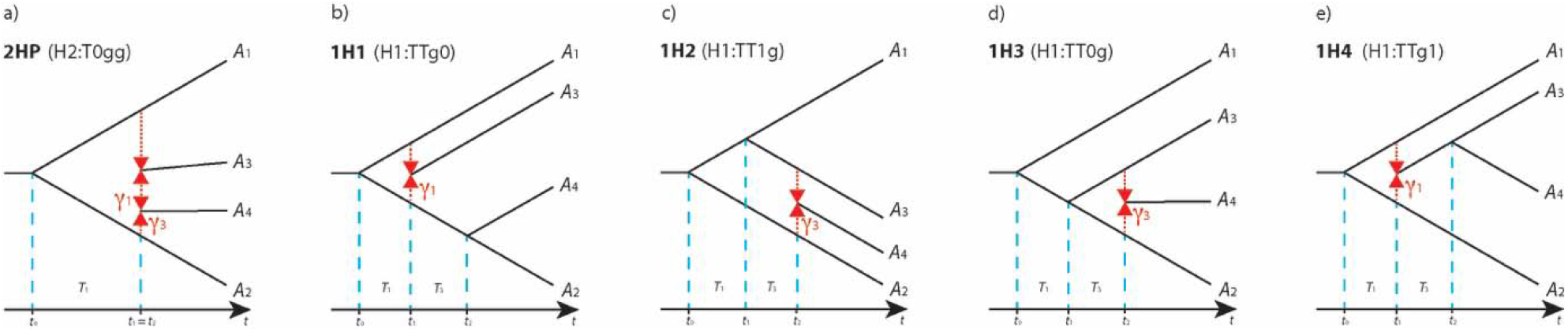
Group θ3. Black arrows below the trees indicate the times of splits (t), with *t*_0_, *t*_*1*_, *t*_*2*_. *t*_*0*_ denoting the initial splitting point, *t*_*1*_ the first hybridization/introgression (red arrows) or split event, and *t*_*2*_ the second hybridization/introgression (red arrows) or split event. *t*_*1*_=*t*_*2*_ indicates simultaneous events. Names (and aliases) of models are presented above the trees. A_1_, A_2_, A_3_, A_4_ indicate the order of lineages in the first permutation (1234), and *T*_1_ and *T*_3_ indicate different periods, *γ*_1_, and *γ*_3_ are the hybridization/introgression coefficients. Parameters in the last two trees are only shown if they are not fixed. a) two simultaneous hybridizations/introgressions, b) hybridization/introgression at *t*_*1*_. c) hybridization/introgression after the first split (between the youngest and most distant branches). d) hybridization/introgression after the first split (between the two most budding sister branches), e) branch derived from hybridization/introgression after initial split represents the last diversification.

**Figure 4.**
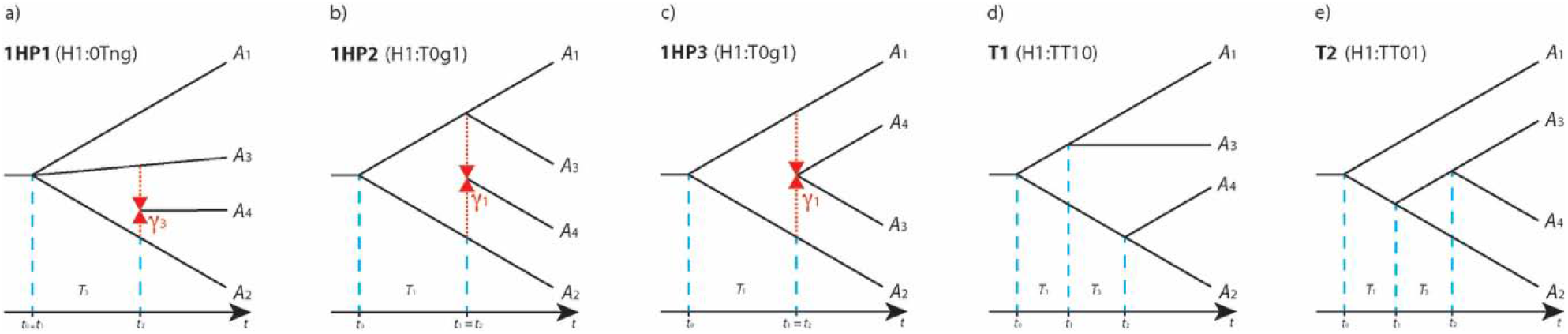
Group θ2. Black arrows below the trees indicate the times of splits (*t*), with *t*_0,1,2_ representing specific time events. *t*_*0*_ indicates the initial splitting point, *t*_*1*_ the first hybridization/introgression (red arrows) or split event, and *t*_*2*_ indicates a second split/hybridization/introgression event. *t*_0_=*t*_1,_ etc., indicate simultaneous events. Models and aliases are labeled above the tree graphics. A_1_, A_2_, A_3_, A_4_ indicate the order of lineages in the first permutation (1234). *T*_*1*_ and *T*_*3*_ indicate different periods and *γ*_1_ and *γ*_3_ are the hybridization/introgression coefficients. The last two sets of parameters are shown only if they are not fixed. a) three branches (A_1_, A_3_, A_2_) were the result of an initial diversification and an incidence of hybridization/introgression (at *t*_2_) between two of these branches. b) hybridization/introgression took place simultaneously with the second diversification (at *t*_1_ = *t*_2_). c) The resulting hybridization/introgression (at *t*_*1*_ = *t*_*2*_) immediately diversified into two branches (A_4_, A_3_). d) Tree variant with two independent, final diversifications (at *t*_1_, *t*_2_). e) Option with two subsequent final diversifications.

**Figure 5.**
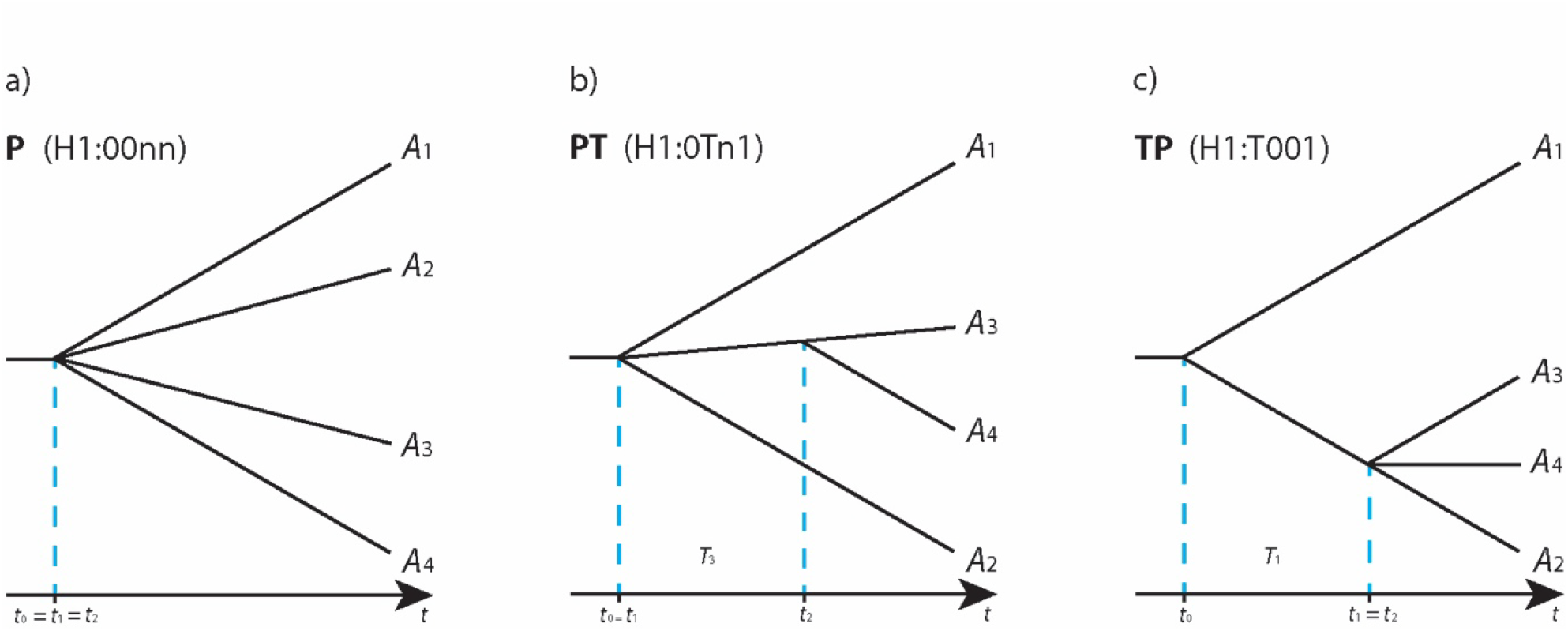
Groups θ1 and θ0. The arrows below the trees indicate the times (*t*) of splits, with *t*_*0*_,*t*_*1*_,*t*_*2*_ being specific events. *t*_*0*_ shows the initial split, *t*_1_ the split, and *t*_2_ the second split. *t*_0_=*t*_1_, etc., signifies that two events appeared simultaneously. The names of models (and aliases) are shown above the graph. A_1_,A_2_,A_3_,A_4_ indicate the order of lineages in the first permutation (1234). *T*_*3*_ and *T*_*1*_ indicate different periods and are shown only if they are not fixed in the respective model. A) The first diversification (at *t*_0_=*t*_1_) is polytomy followed by a dichotomic split. B) The first diversification (at *t*_*0*_) is dichotomy followed by polytomy (at *t*_*1*_=*t*_*2*_). C) Polytomy.

The most complex group Ѳ4 includes 24 permutations of 2H1 (H1:TTgg, Fig. 2a, root model) and 12 permutations from 2H2 (H2:TTgg, Fig. 2b, root model), in which all parameters are free to change.

Group Ѳ3 comprises five specific models, in which three parameters are free and one is fixed. One with two hybridizations/introgressions at the same time after the first split of 2HP (H2:T0gg, six permutations, see Fig. 3a), and four models with single hybridizations/introgressions. Model 1H1 represents a situation where hybridization/introgression follows the initial split before a second one (H1:TTg0, 12 permutations, see Fig. 3b). Model 1H2 represents hybridization/introgression after the initial and first splits and between distant branches (H1:TT1g, 24 permutations, see Fig. 3c). Model 1H3 contains a hybridization/introgression between sister branches following the initial and first splits (H1:TT0g, 12 permutations, Fig. 3d). Model 1H4 shows hybridization/introgression after the initial split and a subsequent second split after hybridization/introgression (H1:TTg1, six permutations, Fig. 3e).

Group Ѳ2 comprises three hybridization/introgression-polytomy trees and two binary tree models, wherein two parameters are free. Model 1HP1 represents hybridization/introgression-polytomy where a second hybridization/introgression coefficient *γ*_3_ is introduced, and the first branch T_1_ is collapsed (H1:0Tng, 12 permutations, Fig. 4a). In model 1HP2, the second branch T_2_ is collapsed and the first hybridization/introgression coefficient *γ*_1_ is set, which indicates that hybridization/introgression and branch divergence happen simultaneously (H1:T0g1, 12 permutations, Fig. 4b, see also Doronina et al. (2017a). Model 1HP3 shows hybridization/introgression combined with diversification of hybridization/introgression branches (H1:T0g1, six permutations, Fig. 4c). The first binary model T1 represents two independent diversifications following the first split (H1:TT10, six permutations, Fig. 4d), while the second model T2 represents three consequent diversifications of an ancestral branch (H1:TT01, 12 permutations, Fig. 4e).

Group Ѳ1 comprises two models with one free parameter. The first model in group Ѳ1 is PT, where the first diversification point forms a polytomy and the second diversification point yields two new branches (H1:0Tn1, six permutations, Fig. 5B). The model TP reflects the situation where a polytomy appears after a second diversification point (H1:T001, four permutations, Fig. 5C). The last group, designated Ѳ0, includes one tree with all four parameters fixed, reflecting polytomy P (00nn, one permutation, Fig. 5A).

For the five groups (Ѳ4 - 36 trees, Ѳ3 - 60 trees, Ѳ2 - 48 trees, Ѳ1 - 10 trees, Ѳ0 - 1 tree), we calculated the optimized likelihood values using only free and defined parameters for each considered tree. From the specific groups ѲN where N={0,1,2,3,4} in each case N+1 parameters (from n_0_, T_1_, T_3_, γ_1_, γ_3_) are used for optimization, and parameter n_0_ is used in all optimizations.

### 3. Statistical testing

The various parameters n_0_, T_1_, T_3_, γ_1_, γ_3_ contribute specifically to the likelihood estimation. However, they don’t have equal influence on all the models. The model parameters are estimated using the maximum likelihood method. Based on the empirical data x, the likelihood function *l*(*x; θ*) is calculated for each of the 155 trees as a function with unknown parameters *θ*:

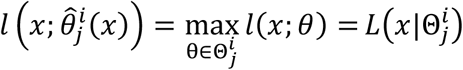

where *i* and *j* are indexes (*i* < *j*, 0 ≤ *j* ≤ 3) and 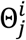 is a chain of parametric sets for a specific tree (see Appendix 1, 1.5.1). These values are then used to estimate the parameters. The numbers of parameters taken in this function vary between groups from five parameters n_0_, T_1_, T_3_, γ_1_, γ_3_ in group Ѳ4 to one parameter n_0_ in group Ѳ0. In the next step, the tree with the highest likelihood function 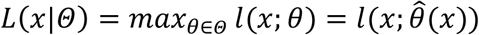 is selected, and likelihood ratios are calculated:

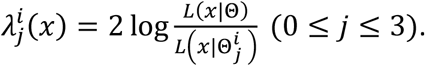

(see Appendix 1, Table S2, and S1.5.1-S1.5.2).

The corresponding p-values are formally defined by the expression S1.5.3, but the exact significance is not necessarily achieved in this case. In practice, the chi-square or chi-square mixtures (e.g., mixing 50% of the chi-square distribution with one degree of freedom with point mass 0 (distribution function = 0); or mixing several chi-square distributions with different degrees of freedom) pre-test approximation S1.5.4 is often used inappropriately in such circumstances. An alternative approach is to use computer simulations. We apply both of these approaches. Because each of the trees of group θ_*j*_ is a particular case of some model of group θ_(*j*+*1*)_(0 ≤ *j* ≤ 3), then *L*(*x*|θ_4_) ≥ *L*(*x*|θ_3_) ≥ *L*(*x*|θ_2_) ≥ *L*(*x*|θ_*1*_) ≥ *L*(*x*|θ_0_).

The general idea of the method is that the hypotheses 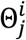, where the condition: corresponding p-value 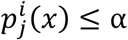 is correct, are rejected, and the simplest tree is selected from the remaining models (depending on the smallest number of free parameters). For selecting the optimal model, we consider the logarithmic likelihood ratio 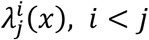 (see Appendix 1, S1.5.2). Then, we get the critical value *Zα*, where α=0.05, from the preliminary chi-square or chi-square mixture distribution or the most reliable eCDF empirical distribution (for eCDF, see below). If the P-value 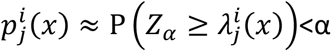, we reject the simplest model; alternatively, accepting 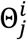. Dependent on the selected algorithm (see below under Statistical algorithms), we search for the first accepting of 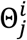, or for the first rejection.

#### Simulations

To select the best fit for the approximation distribution from chi-square or chi-square mixture, we perform simulations using 15 sets of four parameters, T_1_, T_3_, γ_1_, and γ_3_, with 1000 repetitions for each set of parameters. For the reverse algorithm, we use the chi-square and chi-square mixture proposed by Han and Chang (2010) with two geometrical mass points (1 and 0). We use chi-square with one degree of freedom and 50% chi-square with 50% geometrical mass point 0 (Self and Liang 1987) for the stepwise algorithm. For each set of parameters, we calculated initial values of *a*_*i,j*_. For individual repetitions from initial values *a*_*i,j*_, we generate 10 random Poisson-distributed values and use these values for the subsequent optimization and model selection. Then we measure model misrecognition levels for all criteria and algorithm combinations. As a result, we could demonstrate that the chi-square demonstrates a better recognition of complex models (Ѳ4, Ѳ3) and a comparable level of recognition for simple models (Ѳ2, Ѳ1, and Ѳ0) than any of the tested mixtures. Thus, we selected the chi-square calculation for our approximation criterion.

Next, we run simulations for 277 different sets of parameters T_1_, T_3_, γ_1_, and γ_3_ (parameter n_0_ was set to be equal to 100, and reflects the effective size of the initial population in the range between 10,000 to 100,000 what approximate a normal biological situation) with 1000 repetitions for each set of parameters. For each set of parameters, we used a set of Poisson-distributed randomized values and calculated the likelihood ratio similar to the previous simulation. We applied the results of simulations to test the frequency of misrecognition of the model under the chi-square and eCDF statistical analyses. We also generated 75 new parameter sets (5 sets for each model), with T1=0.5, T3=0.5, γ1=0.5, γ3=0.5, to derive parameters. Fixed parameters were applied according to the model used. In these sets, we used ranges of n0 values (n0 = 50, 100, 200, 400, 1000) and 1000 repetitions. For these sets, we also calculated the frequencies of the model misrecognition to see how many markers are sufficient for stable recognition of the often complex phylogenetic scenarios.

#### Empirical distributions

According to the *Law of Large Numbers* (e.g., see (Grimmett and Stirzaker 1992), we generated the *empirical cumulative distribution function* (eCDF) (van der Vaart 1998) of the likelihood ratio for each internal comparison to determine the critical boundary for the given probabilities (see also Appendix 1, chapter 5). In each comparison, we distinguished between two models that we defined as either more or less complex according to their group affiliation (e.g., when comparing tree 2H1:1234 (best from the group Ѳ4) with tree T2:2143 (best from the group Ѳ2), 2H1 is selected as the more complex tree and T2 as the simpler tree). From the optimization processes, we collected values of parameters (*n*_0_, *T*_1_, *T*_3_, *γ*_1_, *γ*_3_) for the simpler tree (e.g., T2) to recover ten initial values of *ξ*_0_ (equivalent to the defined values *y*_*i,j*_). Using a randomization procedure where 10 random Poisson-distributed values were added to *ξ*_0_, we get a 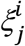, and performed optimization for simple models and all models from the more complex group (e.g., model T2 and all models from group Ѳ4: 2H1, 2H2) to obtain the best trees from both (e.g., best tree related to model T2 and best tree from group Ѳ4) and to determine the likelihood ratios. We then repeated the last steps of the randomization of the initial values and calculation of the likelihood ratio multiple times. By default, for the probability α = 0.05 we performed 500 repetitions. The number of repetitions can be easily extended by the users. We then sorted the likelihood ratios 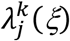 in ascending order. Finally, using the *Law of Large Numbers*, replacing probability by frequency, we can take the approximation: 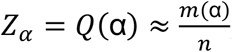, where *m* (α) – the number of values 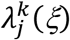 – is more than or equal to α, *n* is the number of repetitions, and *Z*_α_ is the calculated critical boundary for the probability α (e.g., for probability α = 0.05 and 500 repetitions we should take the 475^th^ value as a critical boundary).

#### Statistical algorithms

We propose two statistical algorithms based on likelihood ratio and chi-square or empirical distributions, (1) reverse and (2) stepwise, to find the essential true tree (Fig. 6). We call a resulting tree “essential” in cases when we receive trees that show not the highest likelihood among all trees.

**Figure 6.**
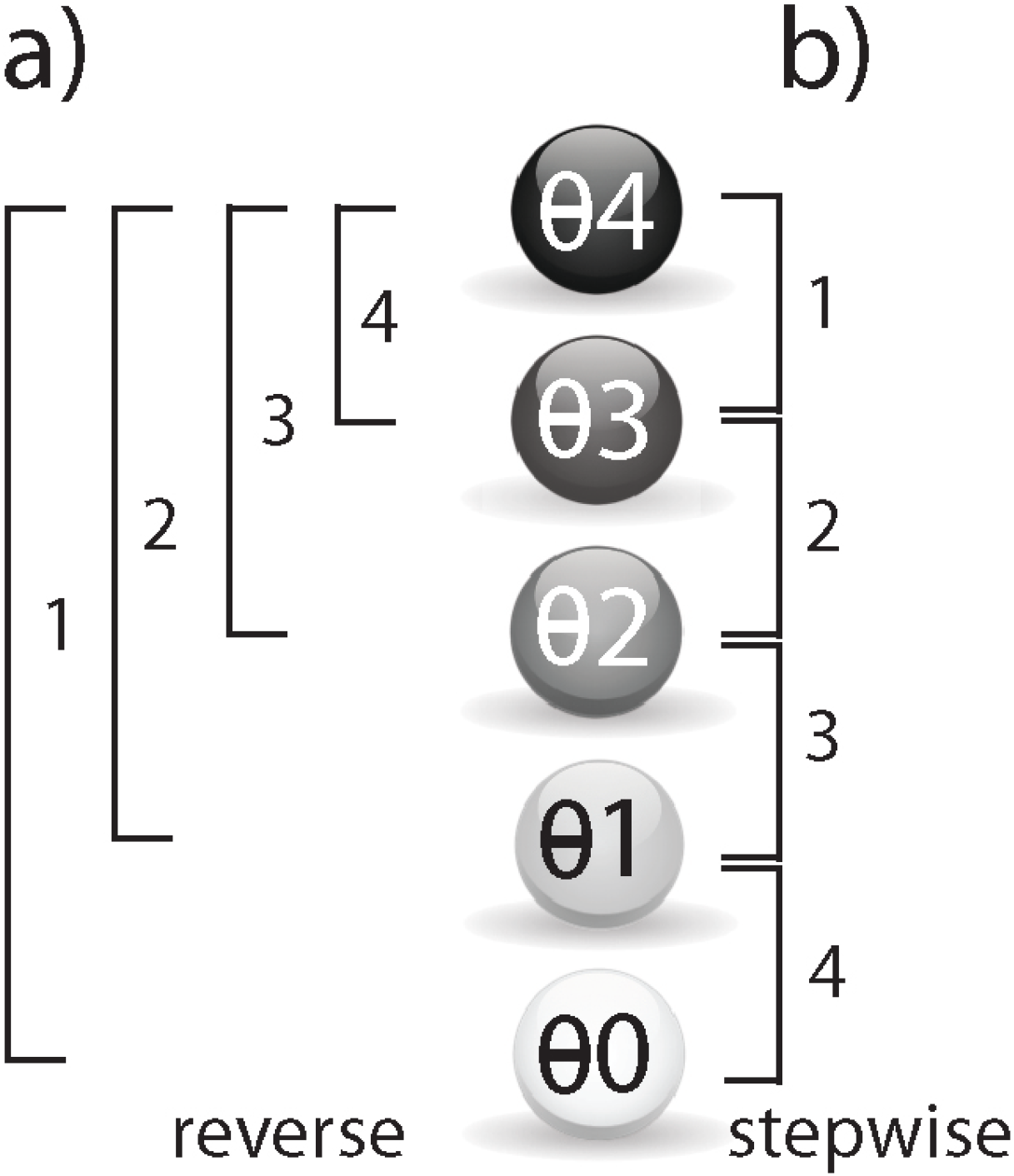
Statistical strategies. Gray circles represent the 5 groups of models from which the best tree estimation (corresponding to maximum likelihood) is derived. Brackets indicate the order of performed comparisons. a) In the reverse method, we compare the most complex tree (Ѳ4) with the simplest tree (Ѳ0), then consistently increase the complexity of the second tree, b) In the stepwise method, we compare the more complex tree with the next less complex one (e.g., Ѳ4 and Ѳ3, Ѳ3, and Ѳ2 etc.). To determine the complexity of the tree, we take the number of parameters in the likelihood function.

#### Reverse method

To find a model with the best likelihood, e.g., of the group Ѳ4, 4-LIN starts by comparing to group Ѳ0 (polytomy). If we cannot reject polytomy, we stop the comparisons and accept polytomy. Alternatively, we start comparing the best model from group Ѳ4 by screening for models from group *Ѳj* where 0 < *j* ≤ 3 until an alternative model cannot be rejected. If such an alternative model cannot be rejected, we accept the best tree from this model. In other cases we increase j by 1 and take the best model from the next alternative group until no more comparisons can be performed, in which case, the best tree from group Ѳ4 is accepted (see Fig. 6a).

#### Stepwise method

We start comparisons with the best model from group Ѳ4 and compare it with the best model from groupѲ3. If we reject the model from groupѲ3, then the best tree from group Ѳ4 is accepted. If the model from group Ѳ3 cannot be rejected, we continue comparisons by taking the best model from group *Ѳ*_*j*_ and comparing it with the best model from group *Ѳ*_*j+1*_, where 0 ≤ *j* <4 or until no more comparisons can be performed, in which case, polytomy is assumed (Fig. 6b).

Given the complexity of the calculations and the optimization formulas for maximizing the likelihood function *l*(*x; θ*), and to receive eCDF critical values, we initially used Wolfram Mathematica 10 (https://www.wolfram.com/mathematica/) to check the source formulas and to test various optimization options.

### 4. Testing phylogenetic diagnostic markers

To demonstrate the effectiveness of the statistics, we applied the 4-LIN test to determine the significance of retrotransposon data supporting known validated phylogenetic trees of primates, domestic dogs, and mouse strains. We used the 2-n-way suite (Churakov et al. 2020) to generate 2-way genome alignments and to extract orthologous presence/absence retrotransposon loci in fasta format. Manual inspection of orthology and correction of MUSCLE alignments ensured reliable information for the computationally extracted loci. Such verified loci were then used to derive the frequency of diagnostic retrotransposon insertions sorted by the introduced ten possible tree topologies for four species.

### Great ape phylogenetic project

To investigate the phylogenetic significance of presence/absence patterns of active *SINE-VNTR-Alu* (SVA) SINEs in great apes, we screened all available great ape genomes. Human (*Homo sapiens*, version hg38), chimpanzee (*Pan troglodytes*, version Clint_PTRv2), bonobo (*Pan paniscus*, version panPan1.1), gorilla (*Gorilla gorilla*, version gorGor4), and orangutan (*Pongo abelii*, version Susie_PABv2) genomes (see Appendix 4) were used to generate 2-way genome alignments (Churakov et al. 2020); http://retrogenomics.uni-muenster.de/tools/twoway) with human as the target genome (human-chimpanzee, human-bonobo, human-gorilla, human-orangutan). With n-way (Churakov et al. 2020); http://retrogenomics.uni-muenster.de/tools/nway), we then created two sets of four species: (1) human, chimpanzee, gorilla, orangutan, and (2) human, chimpanzee, bonobo, gorilla. To extract presence/absence patterns for the ten possible tree topologies for four species, we ran one direct (starting from the human target) and three reverse searches (starting from the three different query genomes) in n-way. We restricted our search to diagnostic SVA retrotransposons active during the diversification of great apes. A maximum of 10-nt truncated ends for the relatively old SVA SINEs were allowed, all duplications were removed, and only perfect matches were considered.

### Dog lineage diversification project

To sort the phylogenetic history of domestic dog breeds we used the gray wolf (version UniMelb_Wolf_Refassem_1, GCA_008641055.1), beagle (version Beagle, GCA_000331495.1), German shepherd (version ASM864105v1, GCA_008641055.1), and boxer (version CanFam3.1, GCA_000002285.2) (see Appendix 4). We built an n-way project (Churakov et al. 2020) for these four dog breeds, with boxers as the target species and the remaining species as queries. All genomes and RepeatMasker reports were downloaded from NCBI. The fastCOEX tool (http://retrogenomics.uni-muenster.de/tools/fastCOEX/index.hbi?; (Doronina et al. 2017b) was applied to extract nearly full-length (< 6-nt truncations for young tRNA-SINEs) genomic dog-specific SINE elements (SINEC_Cf, SINEC_Cf2, and SINECA1_Cf), flanked by repeat-free sequences. We performed one direct and three reverse n-way searches with coordinates of the selected SINEs. All duplicates were removed, and the resulting perfect presence/absence patterns were downloaded as aligned fasta files.

### Mouse strain project

To confirm the phylogenetic relationships among inbred laboratory mouse strains, we downloaded their genomes and RepeatMasker reports from NCBI (see Appendix 4). We utilized CBA/J (CBA_J_v1), C57BL6/J (ASM377452v2), BALB/cJ (BALB_cJ_v1), and DBA/2J (DBA_2J_v1) mouse strain genomes to generate six 2-way genome alignments with CBA/J or C57Bl6/j selected as targets (http://retrogenomics.uni-muenster.de/tools/twoway). We built an n-way project for these four mouse strains and performed two direct n-way searches for SINE/*Alu* and SINE/B2 elements from CBA/J and C57Bl6/J strains (http://retrogenomics.uni-muenster.de/tools/nway). Perfect presence/absence patterns (allowing < 10 nt truncations) with all duplications removed were downloaded as aligned fasta files.

## Results and Discussion

Four-lineage phylogenies require the evaluation of 10 predefined presence/absence patterns for each inserted retrotransposon. In developing 4-LIN, we introduced y-based short names to describe the individual patterns with the two indices *i* and *j. I*=*j* denotes one absence and three presences (e.g., y_11_ = -+++). *J* > *i* denotes two absences and two presences (e.g., y_23_= +--+). We then sorted all ten presence/absence patterns in ascending order as follows: (1) [y_11_] -+++; (2) [y_12_] --++; (3) [y_13_] -+-+; (4) [y_14_] -++-; (5) [y_22_] +-++; (6) [y_23_] +--+; (7) [y_24_] +-+-; (8) [y_33_] ++-+; (9) [y_34_] ++--; (10) [y_44_] +++-(note: neither ++++ nor patterns with 3 minuses are phylogenetically informative).

We built two statistical algorithms with varying sensitivities to potentially short branches. The more robust and routinely used *reverse* algorithm was less reliable in detecting critical short branches and, correspondingly, in handling small numbers of phylogenetic markers. The *stepwise* algorithm was adapted to detect partly resolved trees. For example, the distribution y_11_=22; y_12_=25; y_13_=7; y_14_=11; y_22_=14; y_23_=12; y_24_=18; y_33_=16; y_34_=17; y_44_=24 (selected as an example from simulations) was unresolved by the reverse algorithm, while the stepwise algorithm revealed a partially resolved solution (PT model; H1:0Tn1:1234, T_3_=0.13).

### Command-line likelihood estimator

We also developed a standalone python console script (Python 2.7/3.6 or higher) we call hammlet (Hybridization/Introgression Models Maximum Likelihood Estimator) that can be installed on various operating systems. Hammlet performs likelihood calculations for different tree topologies, utilizes two different methods of statistical comparisons (chi-square and eCDF), finds essential trees for the data to be evaluated, and draws these trees. The command-line procedure leads the user from their data to the final tree with statistical evaluation and requires an input of the frequencies of phylogenetic markers supporting each of the ten possible, carefully evaluated presence/absence patterns (see above) and some additional parameters. It should be noted that the order of values in the user data vector is predetermined (see above). A typical result provides the user with the following sorts of information: the level or group (*N4*: Ѳ4 four parameters variable; *N3*: Ѳ3 three parameters variable and one fixed; *N2*: Ѳ2 two parameters variable and two fixed; *N1*: Ѳ1 one parameter variable and three fixed; *N0*: Ѳ0, polytomy), the model (e.g., PT, Model) and its alias name (e.g., H1:0Tn1, Alias), the permutation of the species orders (e.g., 1423, Order), the likelihood (LL), the effective population size coefficient (n0), the length of the first branch (T1), the length of the second branch (T3), the gamma value for the first hybridization/introgression (g1), and the gamma value for the second hybridization/introgression (g3). The next step is the drawing of the maximum likelihood tree using the command line: *hammlet draw*, followed by the information provided by the above parameters, wherein the initial order of species names A,B,C,D is replaced by A,D,B,C for the permutation 1423. The derived tree outfile is in svg-format and can then be visualized in any internet browser.

In addition to the procedure just described, the command-line interface can also calculate optimized likelihood values for user-defined sets of models and permutations but without statistical calculations (output in a user-defined file with comma-delimited format). For developers, the command-line interface provides a reverse technique to derive specific marker distributions from a preset phylogenetic tree topology. Here, the number of ten *y*_*i,j*_ marker compositions are derived from the five optimized parameters (n0, T1, T3, *γ*_1_, *γ*_3_) with subsequent repetitive Poisson-based randomization, followed by an optimization step. This can be used for the simulation of marker-tree variations. The command-line python script is available at https://github.com/ctlab/hammlet.

### R-based web interface

We used the R-programming language to develop a user-friendly interface. The latest version of hammlet is integrated into a Shiny App. After the user has input the observed frequencies of phylogenetic markers for all of the ten predefined presence/absence patterns for four lineages, they then select the statistical method to be used (Reverse/stringent or Stepwise/relaxed), significance level (0.05 or 0.01, default value is 0.05) and used statistical criterion (chi-square (raw and fast) or empirical distribution (exact but slow)). For empirical distribution user can select sample size (number of simulations, range between 500 and 2500). The program then calls hammlet to optimize parameters and calculate the likelihood of the 155 hypothetical trees, including unique and multiple hybridization/introgression scenarios and polytomies. For each of the five predefined groups (Ѳ4, Ѳ3, Ѳ2, Ѳ1, Ѳ0, see Methods), the tree with the best likelihood value is selected and statistically compared to the simple likely tree, in dependency to selected method. It then provides a visualization of the tree from the best-fitted model and a table of informative parameters of the selected tree. The statistical difference between the best model and polytomy is then shown under the tree figure. For the tree determined to be the most likely, an implemented KKSC module calculates the significance for all individual nodes (splits/hybridizations/introgressions) of the tree presented in the middle section of the final screen. A table of the tree parameters is shown at the bottom section of the screen (see instructions on the application page http://retrogenomics.uni-muenster.de:3838/hammlet/instructions.html). The figure of the likeliest tree and the table of detailed parameters can be downloaded. It is also possible to upload a file containing frequencies of phylogenetic markers for all ten predefined presence/absence patterns for four lineages.

### Simulation

We applied 75,000 sample sets of Poisson-distributed randomized *y*_*j,j*_ for testing different chi-square-based approximation criteria. Interestingly, the classical chi-square application in combination with the reverse algorithm revealed ∼94% of correct models and ∼88% of correct model recognitions for the stepwise algorithm. The best chi-square-based mixture received ∼85% of correct model recognitions for the reverse algorithm (Appendix 2, Table S1). For the stepwise algorithm, ∼50% of chi-square mixture analyses received ∼85% of correct model recognitions (Appendix 2, Table S2). It is to mention that although the chi-square test provides a quick and reliable fit for most of our data, because of certain violations of data independency, some estimations, e.g., the comparison of trees vs. hybridization, did not always receive a correct chi-square approximation. Therefore, we present alternatively the eCDF-based calculation that is entirely independent of the shortcomings of the chi-square analysis but requires more extended processing. Applying 277,000 sample sets of Poisson-distributed randomized *y*_*j,j*_ values, we used hammlet to test the frequencies of incorrectly recovered models from *y*_*j,j*_ values. We submitted these sets to both statistical methods (stepwise and reverse) and used chi-square- and eCDF-based critical values to determine whether the calculated maximum likelihood tree was detected correctly by statistical methods. Interestingly, the combination of chi-square-based critical values and the reverse method yielded approximately 90% of correctly recognized simple models (group Ѳ0, Ѳ1, and Ѳ2). Correct recognition of more complex models (group Ѳ3 and Ѳ4) on the same level (90%) depends on the initial values of parameters T1 and T3. Small values of branch length (T<0.4) could mislead recognition for simple models (group Ѳ1 and Ѳ2) or polytomy (group Ѳ0) (Appendix 2, Table S3). By the stepwise method and chi-square-based critical values, we correctly recovered 70% to 85% of tested simulations with similar dependences from the branch lengths. (Appendix 2 Table S4). Using eCDF and the reverse method, we correctly recognized up to 95% of the simulated models (Appendix 2, Table S4); the stepwise method yielded 89% correct model recognitions (Appendix 2, Table S5) but with similar dependences of the branch lengths. Based on these data, we can conclude that the eCDF-based statistics retrieved more stable results with randomized datasets than the chi-square-based statistics. However, because eCDF performs additional optimizations and therefore is time-consuming (from 10^th^ of minutes up to more than 1 hour of hammlet runs), a chi-square calculation can be used for raw and fast hammlet runs, providing a preliminary check of tree topologies.

We used another set of 75 samples to find the optimal number of markers needed for correct recognition of simple and complex models. From this set, we derived the correct recognition of 90% of models from Ѳ3 and Ѳ4 groups for all tested methods started from an initial value of n0=400 (equivalent to around 400/0.6≅667 markers). Complete, correct recognition at 100% of the most complex models 2H1 and 2H2 could be reached at the level of >1500 markers (Appendix 2, Table S6).

### Connection between trees and presence/absence data patterns

The 4-LIN test can easily identify significantly supported phylogenetic trees based on conflict-free inserted phylogenetic presence/absence data without maximum likelihood optimization. However, in cases when conflicting markers are present that interfere with the reconstruction of simple trees, one of two variants of partial polytomy emerges; for example, a non-zero value for y_12_ can be interpreted as a PT tree (H1:0Tn1:1234), and a non-zero value for y_44_ can be construed as a TP tree (H1:T00n:4123). Two non-zero values in some cases can also be interpreted as an utterly non-conflicting tree; for example, non-zero values for y_13_ and y_24_ can be construed as a T1 tree (H1:TT10:1234), and finally, non-zero values for y_33_ and y_34_ can be interpreted as a T2 tree (H1:TT01:3412). It should be mentioned that in the case of two non-zero values and the T1 model, the order of lineages depends on the relative values; for example, in the case of y_13_ and y_24_, if y_13_>y_24_ the order of species will be 1234, while in the opposite case, it will be 2143. However, the T2 model is free of such dependencies. In Appendix 3, Table S1, all possible tree variants defined by one or two non-zero values are presented and can be used as a reference list for non-conflicting datasets.

To derive a simple phylogenetic tree from the often diverse and sometimes conflicting presence/absence patterns of phylogenetic markers, taking into account stochastic variation and different possible permutations, it is necessary to use statistical applications. However, KKSC or other statistical applications, which consider only qualitative criteria, are less efficient at finding the actual tree topology for four lineages. In contrast to these strict qualitative approaches, 4-LIN (based on maximum likelihood ratios) evaluates each possible branch independently and summarizes this information only at the last stage of likelihood calculation to efficiently reconstruct the complete most probable 4-LIN tree.

However, the significance values calculated by the hammlet script refer to support for the entire tree topology vs. polytomy and do not consider individual branches. Individual significance values for specific branches are more complex to derive because the fixation of an ancient marker at one branch is not dependent on fixation of this marker at neighboring branches (Kuritzin et al. 2016). To obtain statistical support values for individual branches, we embedded elements of the KKSC-statistical calculation (Kuritzin et al. 2016) to compute the significance of each split. Using the KKSC-method in combination with the 4-LIN statistics to calculate specific support for individual branches requires six values (e.g., for tree T1 and order of species 1234 in a first comparison Y_1_=y_24_+y_44_; Y_2_=y_23_+y_33_; Y_3_=y_12_+y_11_ and in a second comparison Y_1_=y_13_+y_33_; Y_2_=y_12_+y_22_; Y_3_=y_14_+y_44_). In Appendix 3, Table S2, we present formulas to connect the results of the 4-LIN test with the KKSC-statistics for all possible resolved trees applied at http://retrogenomics.uni-muenster.de:3838/hammlet/.

### Examples for well-known phylogenetic relations

#### Great apes

SINE elements were previously used as phylogenetic markers to resolve the great ape sister relationship between humans and chimpanzees (Salem et al. 2003). To test the 4-LIN statistics in great apes, we first searched the genomes of humans, chimpanzees, gorillas, and orangutans for SVA SINE elements (active retrotransposons in great apes). This analysis detected 56 diagnostic markers whose presence/absence patterns were distributed as follows: y_11_=0; y_12_=0; y_13_=0; y_14_=0; y_22_=0; y_23_=0; y_24_=1; y_33_=0; y_34_=37; y_44_=18. Running the 4-LIN statistical test for this dataset (with both the stringent reverse and relaxed stepwise algorithms) and a cut-off value of p = 0.05, we found the maximum likelihood for tree T2 (H1:TT01:4312, T_1_=1.93, T_3_=3.36, p < 2.06.10^−09^), which confirms the current view on hominid phylogeny (((human, chimpanzee), gorilla), orangutan); (Salem et al. 2003) (Fig. 7a). Here both individual splits were also shown to be significantly supported (p = 2.58e-09, p = 5.7e-17, KKSC-test).

**Figure 7.**
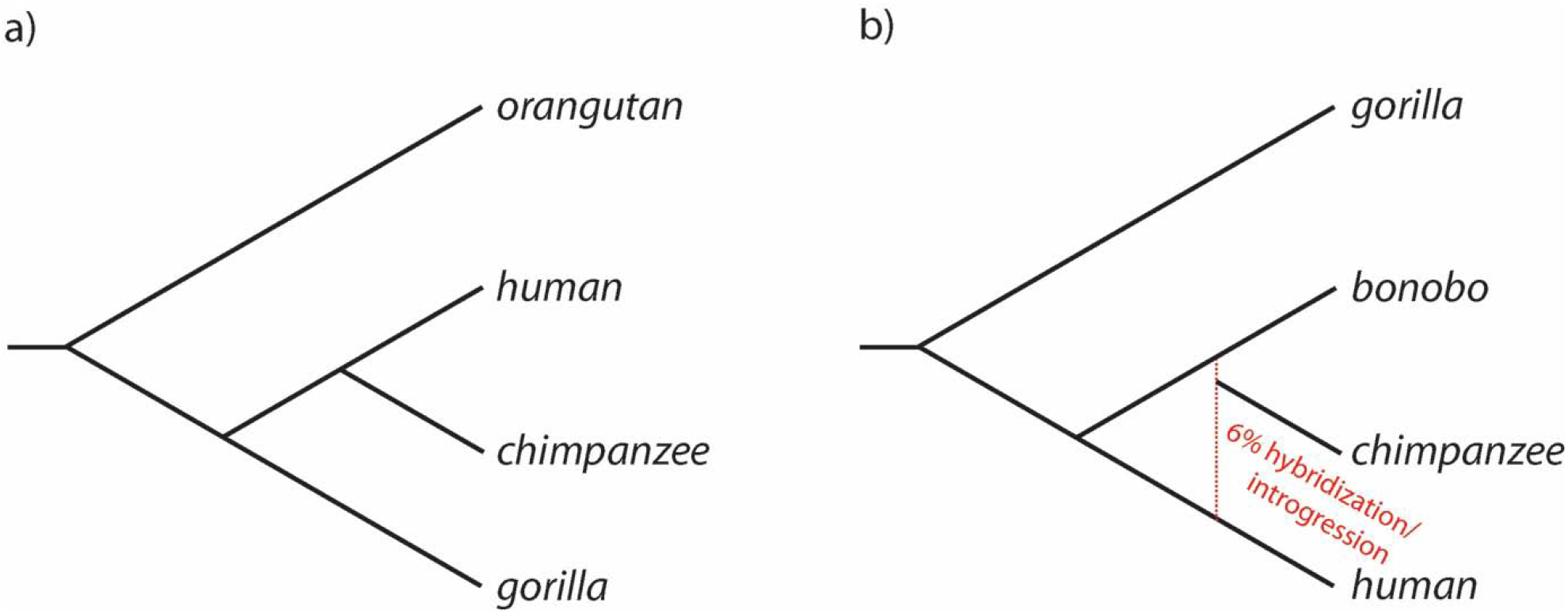
Phylogenetic trees of hominids. Black lines indicate tree branches; the red vertical dotted line shows hybridization/introgression. 7a) most likelihood tree for orangutans, humans, chimpanzees, and gorillas, 7b) hybridization/introgression scenario for gorillas, bonobos, chimpanzees, and humans.

We also screened a second set of species comprising humans, chimpanzees, bonobos, and gorillas. The 52 detected presence/absence patterns revealed the following distribution: y_11_=0; y_12_=0; y_13_=0; y_14_=43; y_22_=0; y_23_=1; y_24_=0; y_33_=0; y_34_=3; y_44_=5. The 4-LIN test for this data set favored the hybridization/introgression model 1H3 (H1:TT0g:4132, T_1_=0.81, T_3_=8.01, γ_3_=0.94). The most likely supported tree shows a close relationship between chimpanzee and bonobo, with humans as the sister group and gorilla as the outermost diversification (however, the KKSC-test did not show significant support for this split (p = 0.33)). This also agrees with the current view of their relationships (Salem et al. 2003). However, we also detected a distinct signal of ancestral hybridization/introgression (6%) between human and chimpanzee (this hybridization/introgression signal was not detected by the KKSC-test; p = 0.25; Fig. 7b).

#### Dogs

Domestic dogs (*Canis lupus familiaris*) evolved from the wolf lineage about 20,000-40,000 years ago and have since diversified into approximately 400 different breeds (Galibert et al. 2011). The large number of dog-specific SINE elements that were active over a long period (Peng et al. 2018) and the rapid diversification of lineages boosted by domestication makes domesticated dogs a good example for testing the 4-LIN statistical approach. We first used 2-n-way (Churakov et al. 2020) to find and extract diagnostic presence/absence markers from 2-way alignments generated from the genomes of the gray wolf, beagle, German shepherd, and boxer. Screening yielded 1221 SINE markers distributed over all 10 predefined presence/absence patterns as follows: y_11_=0; y_12_=2; y_13_=2; y_14_=506; y_22_=0; y_23_=0; y_24_=177; y_33_=1; y_34_=6; y_44_=527. The initial order of breeds was a boxer, beagle, German shepherd, and gray wolf. After running the 4-LIN test using both the reverse and stepwise algorithms and a cut-off value of p = 0.05, we found significant support for the model 1H3 ((H1:TT0g:4132) T_1_=3.57; T_3_=2.15; γ_3_=0.74, p<1.10^−64^), where beagle and German shepherd are closest, the boxer is the sister group to them (p = 1e-64, KKSC-test), and gray wolf is the clear outgroup. However, the boxer ancestral branch showed 36% of hybridization/introgression with the beagle ancestral branch resulting in the German shepherd ancestral branch (p = 1.07e-43, KKSC-test; Fig. 8).

**Figure 8.**
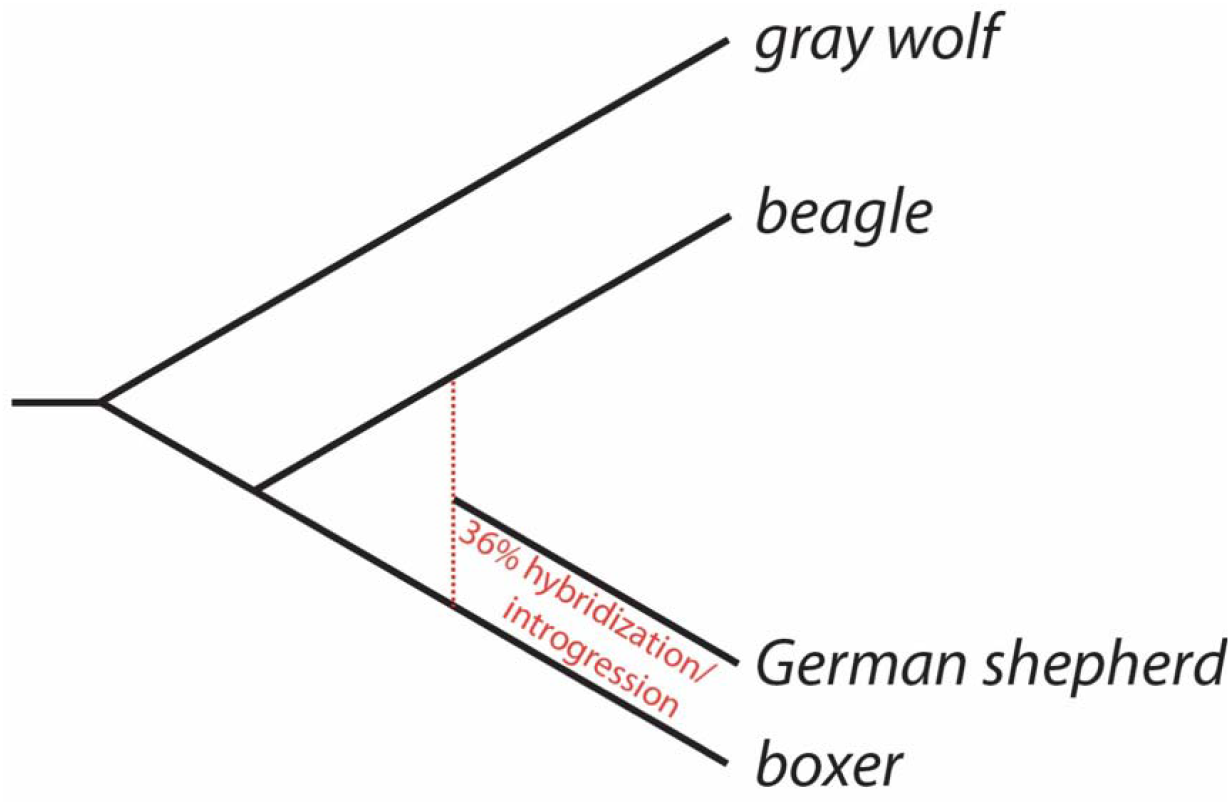
Phylogenetic tree of domestic dog breeds. Black lines indicate tree branches; red vertical dotted line shows hybridization/introgression events.

The boxer, beagle, and German shepherd fusion and their relationship to the gray wolf progenitor, confirm the findings of other phylogenetic studies (Parker 2012; Parker et al. 2017). The relative positions of the three investigated dog breeds fit the previously published tree (vonHoldt et al. 2010), where beagle and German shepherd are closest, and boxer is the sister group. A strong signal of hybridization/introgression between ancient branches leading to boxer and beagle indicates involvement of artificial selection (domestication) and interbreed crossings in the dog’s history. Our dog breed results show that the 4-LIN test can be helpful not only in reconstructing deep phylogenetic events but also in evaluating the history of domesticated breeds.

#### Mouse inbred strains

Mouse inbred strains started their human-laboratory diversification around 175 years ago (Atchley and Fitch 1991). Since then, more than 110 stable strains have been distributed in scientific laboratories worldwide. Due to their inbred nature, these strains carry many metabolic changes and are used in a wide range of biological experiments (Ghazalpour et al. 2012). For our investigations, we selected four mouse inbred strains: CBA/J, C57BL6/J, BALB/cJ, and DBA/2J, located most distantly on the phylogenetic mouse tree. A screen for SINE/*Alu* and SINE/B2 elements yielded 2575 markers distributed over all 10 predefined presence/absence patterns as follows: y_11_=341; y_12_=185; y_13_=215; y_14_=157; y_22_=356; y_23_=197; y_24_=258; y_33_=197; y_34_=147; y_44_=522. The 4-LIN test run with the reverse algorithm and a cut-off value of p = 0.05 showed significant support for the 1H4 model (H1:TTg1:2413, T_1_=0.22; T_3_=0.17; γ_1_=0.7, p<1.10^−64^, Fig. 9). The tree confirms a consolidation of CBA/J and BALV/cJ strains against C57Bl/6J and DBA/2J strains (p = 2.18e^-12^, KKSC-test). However, a hybridization/introgression scenario in the ancestry of the C57Bl/6J and DBA/2J strains is also significantly supported (p = 0.00003, KKSC-test). Such an ancestral hybridization/introgression event might indicate the documented crosses between strains during the early stages of modern mouse laboratory diversification (Atchley and Fitch 1993; Atchley and Fitch 1991).

**Figure 9.**
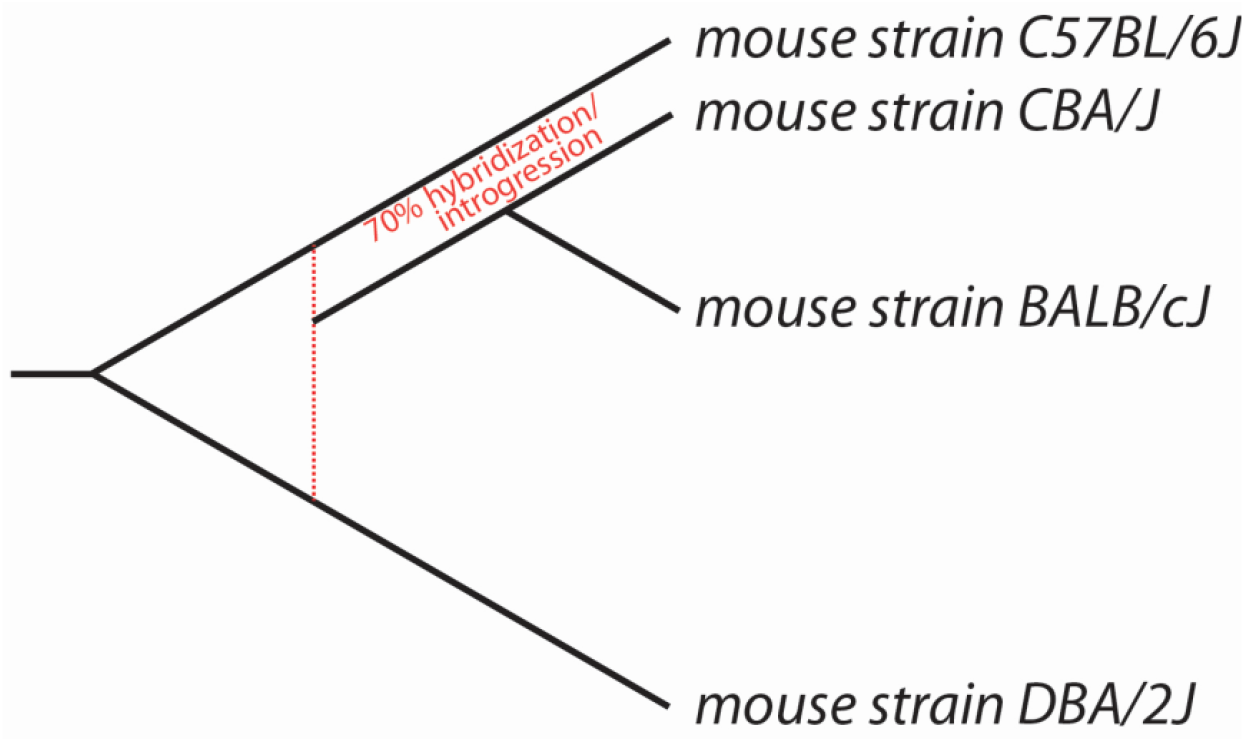
Phylogenetic tree of four inbred mouse strains. Black lines indicate the tree branches; the red vertical dotted line shows the ancestral hybridization/introgression event.

## Conclusions

Based on the ten possible presence/absence distribution patterns of phylogenetically informative retrotransposon markers for four lineages, the 4-LIN test calculates the best fitting phylogenomic tree within a maximum likelihood framework. It determines the statistical significance of a dataset’s support for a given tree compared to that of the next less-supported candidate tree. The need to provide a 4-LIN test and to develop the complex underlying mathematical framework arose from the growing interest in more extensive phylogenomic comparisons of whole-genome retrotransposon presence/absence patterns, where multiple comparisons of groups of only 3 species are inefficient. Compared to the currently available KKSC-statistics for evaluating phylogenomic data that measures three lineages and seven tree variants, the 4-LIN test is finetuned to compare 4 lineages and 155 different trees, including scenarios for multiple ancestral hybridizations/introgressions and incomplete lineage sorting. The contribution of ancestral hybridization/introgression to the phylogeny of species can now be quantified and applied to the reconstruction of complete tree topologies.

Moreover, the 4-LIN statistical approach is combined with the KKSC statistics to thoroughly evaluate complete tree topologies including their individual branches. However, the 4-LIN test is generally more sensitive than the KKSC and can resolve weak ancestral hybridization/introgression signals or very short branches (see, e.g., second hominid example). While the 4-LIN test is sufficient for resolving most phylogenomic conundrums, the ultimate goal of developing the 4-LIN test was to derive a strategy for an n-lineage analysis operating on complete presence/absence matrices of any number of lineages. The maximum likelihood approach and all the necessary mathematical background calculations we have described here provide the necessary technical underpinnings to target such a goal. The current advantage of 4-LIN comparisons is that the results of extended screenings involving four instead of three species/lineages are directly applicable to resolve more complex phylogenetic questions. This enables evaluation of data collected for ten different potential elementary tree topologies instead of just three with KKSC. For example, we used the 4-LIN test to evaluate the significance of phylogenetic marker distributions in great apes, dog breeds, and mouse strains.

## Supporting information

Matematical model

Supplementary tables (symulations)

Supplementary tables (non-conflicting patterns)

Sources of genomes used and RepeatMasker reports.

## Software availability

The new Web tool, including the methods presented here, can be found at: http://retrogenomics.uni-muenster.de:3838/hammlet/

https://github.com/ctlab/hammlet

## Acknowledgments

We thank Marsha Bundman for editorial help. We thank Norbert Grundmann for his help in integrating the shiny framework into a web server. Thanks to the two anonymous reviewers for their essential suggestions to significantly improve the manuscript. This work was supported by the Deutsche Forschungsgemeinschaft (DFG) grant number (SCHM 1469/10-1 to JS) and DFG grant number (281125614/GRK2220 to the Research Training Group Evolutionary processes in Adaptation and Disease [EvoPAD]).

## Author contributions

GC, AK, and JS conceived the 4-lineage project. GC and AK developed and optimized the statistical strategy. CK and VU designed python scripts for automatic likelihood maximization and statistical evaluation. FZ and FW designed R-scripts for the web interface and statistical evaluation. GC and JS wrote the paper with input from all other authors.

## Competing interests

The authors declare no competing interests.

## Notes

### Competing Interest Statement

The authors have declared no competing interest.

### Summary of Updates

Manuscript was significantly rewritten, statistic algorithms were updated and extended, addition necessary simulations were performed.

## References

Atchley, W. R., and W. Fitch. 1993. Genetic affinities of inbred mouse strains of uncertain origin. Mol Biol Evol 10:1150–69.

Atchley, W. R., and W. M. Fitch. 1991. Gene trees and the origins of inbred strains of mice. Science 254:554–8.

Churakov, G., F. Zhang, N. Grundmann, W. Makalowski, A. Noll, L. Doronina, and J. Schmitz. 2020. The multicomparative 2-n-way genome suite. Genome Res 30:1508–1516.

Doronina, L., G. Churakov, A. Kuritzin, J. Shi, R. Baertsch, H. Clawson, and J. Schmitz. 2017a. Speciation network in Laurasiatheria: retrophylogenomic signals. Genome Res 27:997–1003.

Doronina, L., A. Matzke, G. Churakov, M. Stoll, A. Huge, and J. Schmitz. 2017b. The Beaver’s Phylogenetic Lineage Illuminated by Retroposon Reads. Sci Rep 7:43562.

Doronina, L., O. Reising, H. Clawson, D. A. Ray, and J. Schmitz. 2019. True Homoplasy of Retrotransposon Insertions in Primates. Syst Biol 68:482–493.

Fisher, R. A. 1922. On the mathematical foundations of theoretical statistics. Philos. Trans. Roy. Soc. London Ser.A 222:309–368.

Galibert, F., P. Quignon, C. Hitte, and C. Andre. 2011. Toward understanding dog evolutionary and domestication history. C R Biol 334:190–6.

Ghazalpour, A., C. D. Rau, C. R. Farber, B. J. Bennett, L. D. Orozco, A. van Nas, C. Pan, H. Allayee, S. W. Beaven, M. Civelek, R. C. Davis, T. A. Drake, R. A. Friedman, N. Furlotte, S. T. Hui, J. D. Jentsch, E. Kostem, H. M. Kang, E. Y. Kang, J. W. Joo, V. A. Korshunov, R. E. Laughlin, L. J. Martin, J. D. Ohmen, B. W. Parks, M. Pellegrini, K. Reue, D. J. Smith, S. Tetradis, J. Wang, Y. Wang, J. N. Weiss, T. Kirchgessner, P. S. Gargalovic, E. Eskin, A. J. Lusis, and R. C. LeBoeuf. 2012. Hybrid mouse diversity panel: a panel of inbred mouse strains suitable for analysis of complex genetic traits. Mamm Genome 23:680–92.

Glynn, P. W. 1990. Diffusion Approximations. Elsevier Science

Publishers B. V. Grimmett, G. R., and D. R. Stirzaker. 1992. Probability and Random Processes: Problems and Solutions, 3rd edition. Oxford University Press.

Han, S. S., and J. T. Chang. 2010. Reconsidering the asymptotic null distribution of likelihood ratio tests for genetic linkage in multivariate variance components models under complete pleiotropy. Biostatistics 11:226–41.

Kimura, M. 1955a. Solution of a Process of Random Genetic Drift with a Continuous Model. Proc Natl Acad Sci U S A 41:144–50.

Kimura, M. 1955b. Stochastic processes and distribution of gene frequencies under natural selection. Cold Spring Harb Symp Quant Biol 20:33–53.

Kuritzin, A., T. Kischka, J. Schmitz, and G. Churakov. 2016. Incomplete Lineage Sorting and Hybridization Statistics for Large-Scale Retroposon Insertion Data. PLoS Comput Biol 12:e1004812.

Parker, H. G. 2012. Genomic analyses of modern dog breeds. Mamm Genome 23:19–27.

Parker, H. G., D. L. Dreger, M. Rimbault, B. W. Davis, A. B. Mullen, G. Carpintero-Ramirez, and E. A. Ostrander. 2017. Genomic Analyses Reveal the Influence of Geographic Origin, Migration, and Hybridization on Modern Dog Breed Development. Cell Rep 19:697–708.

Peng, C., L. Niu, J. Deng, J. Yu, X. Zhang, C. Zhou, J. Xing, and J. Li. 2018. Can-SINE dynamics in the giant panda and three other Caniformia genomes. Mob DNA 9:32.

Salem, A. H., D. A. Ray, J. Xing, P. A. Callinan, J. S. Myers, D. J. Hedges, R. K. Garber, D. J. Witherspoon, L. B. Jorde, and M. A. Batzer. 2003. Alu elements and hominid phylogenetics. Proc Natl Acad Sci U S A 100:12787–91.

Self, S. G., and K. Y. Liang. 1987. Asymptotic properties of maximum likelihood estimators and likelihood ratio tests under nonstandard conditions. Journal of the American Statistical Association 82:605–610.

van der Vaart, A. W. 1998. Asymptotic statistics.

vonHoldt, B. M., J. P. Pollinger, K. E. Lohmueller, E. Han, H. G. Parker, P. Quignon, J. D. Degenhardt, A. R. Boyko, D. A. Earl, A. Auton, A. Reynolds, K. Bryc, A. Brisbin, J. C. Knowles, D. S. Mosher, T. C. Spady, A. Elkahloun, E. Geffen, M. Pilot, W. Jedrzejewski, C. Greco, E. Randi, D. Bannasch, A. Wilton, J. Shearman, M. Musiani, M. Cargill, P. G. Jones, Z. Qian, W. Huang, Z. L. Ding, Y. P. Zhang, C. D. Bustamante, E. A. Ostrander, J. Novembre, and R. K. Wayne. 2010. Genome-wide SNP and haplotype analyses reveal a rich history underlying dog domestication. Nature 464:898–902.

Waddell, P. J., H. Kishino, and R. Ota. 2001. A phylogenetic foundation for comparative mammalian genomics. Genome Inform 12:141–54.

Wright, S. 1933. Evolution in Mendelian populations. Genetics 16:97–159.

