## Supplementary material for "A 4-lineage statistical suite to evaluate the support of large-scale retrotransposon insertion data to reconstruct evolutionary trees": Matematical model

### Detailed description of mathematical models.

#### 1. Model assumptions

First, let $A_1$, $A_2$, $A_3$, $A_4$ be the names of lineages, and branch $B_k$ the evolution of some isolated population over a time interval $t \in \Delta_k$ with the effective size $N_k(t)$. For any neutral insertion of a potential phylogenetically informative retroelement in some generation $t$ of branch $B_k$, where $k$ ($0 \leq k \leq 4$) is the index of specific branches, we consider ten different events $\omega_{i,j}$ ($1 \leq i \leq j \leq 4$) for four lineages $A_1$, $A_2$, $A_3$, $A_4$, where:

$\omega_{i,i}$ indicates that a retroelement in the orthologous locus is absent in lineage $A_i$ and present in the other three lineages;

$\omega_{i,j}$ indicates that a retroelement in the orthologous locus is absent in lineages $A_i$ and $A_j$ ($i \neq j$) and present in the other two lineages.

We denote:

$\nu_k(t)$ - the quantity of all insertions of retroelements in generation $t$ of the branch $B_k$;

$p_{i,j}^{k}(t)$ - the probability of $\omega_{i,j}$ for every insertion

$\mu_{i,j}^{k}(t)$ - the quantity of $\omega_{i,j}$ for generation $t$ of the branch $B_k$

Considering that the probability of new insertions for each individual of a population ($\alpha_k$ ($t$)) is small and the effective size of population for generation $t$: $N_k(t)$ is sufficiently large, we assume that $\nu_k(t)$ is a Poisson-distributed random variable with a mean:

$$n_k = n_k(t) = 2\, N_k(t)\, \alpha_k\, (t). \qquad \text{(S1.1.1)}$$

Then assuming that all insertions are independent events, we can conclude that $\mu_{i,j}^{k}(t)$ is a Poisson-distributed random variable with parameters $b_{i,j}^{k}(t) = n_k(t) p_{i,j}^{k}(t)$.

Now we consider all possible generations of different branches $B_k$ with potential retrotransposon insertions that are phylogenetically informative. Then the total number of retrotransposon insertions with properties $\omega_{i,j}$ equal to the sum:

$$\xi_{i,j} = \sum_{k} \sum_{t \in \Delta_k} \mu_{i,j}^{k}(t) \qquad \text{(S1.1.2)}$$

- are random, independent, Poisson-distributed variables with parameters:

$$a_{i,j} = \sum_k \sum_{t \in \Delta_k} b_{i,j}^{k}(t) = \sum_k \sum_{t \in \Delta_k} n_k(t) p_{i,j}^{k}(t),$$

or

$$a_{i,j} = \sum_{k} a_{i,j}^{k}\,, \qquad \text{(S1.1.3)}$$

where

$$a_{i,j}^{k} = \sum_{t \in \Delta_k} n_k(t) p_{i,j}^{k}(t).$$

Assuming that, as $t$ increases by 1 (the transition to the next generation), the corresponding functions are slowly changing; we can replace the summation by integration over the appropriate intervals:

$$a_{i,j}^{k} = \int_{\Delta_k} n_k(t) p_{i,j}^{k}(t) dt \qquad \text{(S1.1.4)}$$

#### 2. Elements of Kimura’s neutral theory of evolution

In this section, we repeat considerations presented previously in (Kuritzin et al. 2016) for the 4-

lineage situation.

We denote $X(t)$ as the frequency of one particular marker (retroelement) locus in the population at a time $t$. We consider $X(t)$ as a Markov process with the transition function $u(s, p, t, x)$, reflecting the conditional probability density of $X(t)$ for the condition $X(s) = p$ ($s < t$). This follows the standard Wright-Fisher coalescent model (Fisher 1922, Wright 1931), and then, using the diffusion approximation, this transition function follows the forward Kolmogorov's equation:

$$\frac{\partial_u}{\partial_t} = \frac{1}{4N(t)} \frac{\partial^2}{\partial x^2} [x(1-x)u] \tag{S1.2.1}$$

with the initial condition $u(s, p, s, x) = \delta ( x - p)$, where $\delta (x - p)$ is a Dirac delta function, and $N(t)$ denotes the effective population size at time $t$ (Kimura 1955a). The solution of this equation is represented in the form of a series, including Gegenbauer polynomials (Tran et al. 2013). For our purposes, it is sufficient to know some moments of the distribution of $X(t)$.

We denote

$$m_k(s,p,t) = \int_0^1 x^k u\,(s,p,t,x)\,dx \tag{S1.2.2}$$

which is the $k$-th (conditional) moment of distribution of $X(t)$ about 0.

Following Kimura (1955b), we can denote

$$m_1(s,p,t) = p, \tag{S1.2.3}$$

and $m_k(s,p,t)$ is a solution of the next differential equation:

$$\frac{d}{dt} m_k(s,p,t) = -\frac{k(k-1)}{4N(t)} (m_k(s,p,t) - m_{k-1}(s,p,t)) \tag{S1.2.4}$$

Instead of $t$, we introduce the new independent variable

$$\tau = \tau(s,t) = \int_s^t \frac{dt}{2N(t)} \tag{S1.2.5}$$

which is the “drift time” according to Waxman (2011).

We can then write equation (S1.2.4) in the form:

$$\frac{dm_k}{d\tau} = -\frac{k(k-1)}{2} (m_k - m_{k-1}) \tag{S1.2.6}$$

with the initial condition: $m_k|_{\tau=0} = p^k$.

We need the 2nd, 3rd, and 4th moments. Solving the equation (S1.2.6) for $k$ = 2, 3, 4, considering (S1.2.3), we obtain:

$$m_2(s,p,t) = p^2e^{-\tau} + p(1-e^{-\tau)}) \tag{S1.1.7}$$

$$m_3(s,p,t) = p^3e^{-3\tau} + \frac{3}{2}p^2(1-e^{-2\tau})e^{-\tau} + \frac{1}{2}p(1-e^{-\tau})^2(e^{-\tau}+2) \tag{S1.1.8}$$

$$\begin{aligned} m_4(s,p,t) &= \\ &= p^4e^{-6\tau} + 2p^3(1-e^{-\tau})(e^{-2\tau}+e^{-\tau}+1)e^{-3\tau} + \\ &+ \frac{3}{5}p^2(1-e^{-\tau})^2(2e^{-3\tau}+4e^{-2\tau}+6e^{-\tau}+3)e^{-\tau} + \\ &+ \frac{1}{5}p(1-e^{-\tau})^3(e^{-3\tau}+3e^{-2\tau}+6e^{-\tau}+5) \end{aligned} \tag{S1.1.9}$$

where $\tau = \tau(s,t)$, according (S1.2.5).

#### 3. Mathematical models

We distinguish two main speciation variants (tree topologies), assuming two hybridizations each.

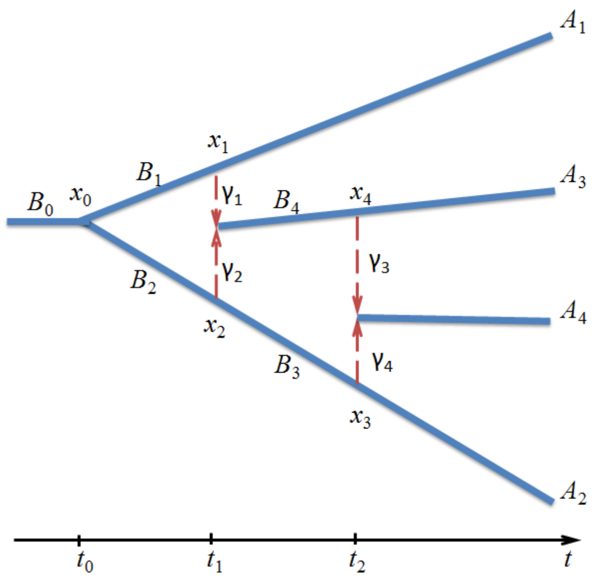

Figure 1. Tree 2H1

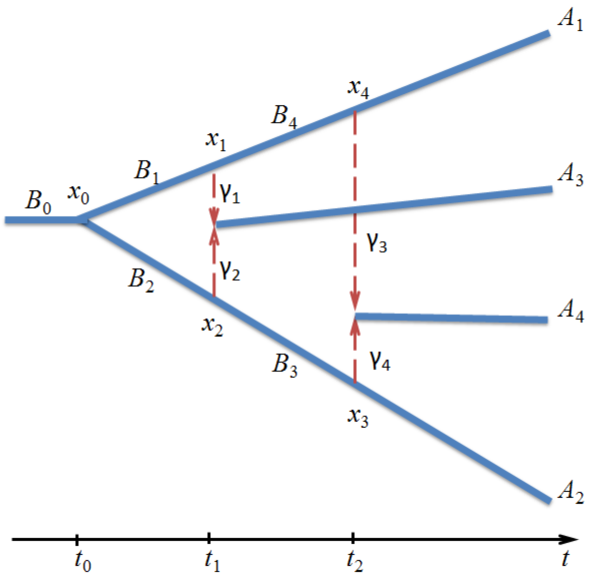

Figure 2. Tree 2H2

In these two trees, we use the fusion model of hybridization (Kuritzin et al. 2016), where the third and fourth branches arise from two ancestral populations at the points $x_1$-$x_2$ and $x_3$-$x_4$.

The proportions of the two fused sub-populations ($B_1$ and $B_2$) forming a new population are denoted by $\gamma_1$ and $\gamma_2$, respectively ($\gamma_1 + \gamma_2 = 1$). For the second point of fusion, the proportions of the sub-populations $B_3$ and $B_4$ are $\gamma_3$ and $\gamma_4$, respectively, ($\gamma_3 + \gamma_4 = 1$), and the corresponding intervals of $t$ for the branches are: $\Delta_1 = \Delta_2 = (t_0, t_1)$, $\Delta_3 = \Delta_4 = (t_1, t_2)$, and $\Delta_0 = (-\infty, t_0)$ .

It should be noted that for each hybridization-affected tree topology, we can have 24 permutations of the four lineages $A_1$, $A_2$, $A_3$, $A_4$. However, for the tree topology in Figure 2, due to symmetry, the number of different rearrangements may be reduced to 12.

#### 3.1. Hybridization topology 2H1

Retrotransposon insertions later shared between lineages might occur at any branch; here, we consider the five branches $B_0$, $B_1$, $B_2$, $B_3$, $B_4$.

##### 3.1.1. Branch $B_0$

For a retrotransposon insertion on the branch $B_0$ at $t < t_0$, frequencies at the right ends of the respective branches form the random vector $\mathcal{X} = (X_0, X_1, X_2, X_3, X_4)$.

Then, by introducing the (non-random) vector $x = (x_0, x_1, x_2, x_3, x_4)$, we note that the probability density function of vector $\mathcal{X}$

$$\begin{gathered} f^0(x) = u_0(t, p, t_0, x_0) u_1(t_0, x_0, t_1, x_1) u_2(t_0, x_0, t_1, x_2) \\ u_3(t_1, x_2, t_2, x_3) u_4(t_1, \gamma_1 x_1 + \gamma_2 x_2, t_2, x_4) \end{gathered}, \qquad \text{(S1.3.1)}$$

where $p = (2\,N_0(t))^{-1}$, and the index of the function $u$ indicates the tree branch number.

Let $\omega_j$ be when the insertion is fixed in the lineage $A_j$.

Then, as it follows from the neutral theory of molecular evolution (Kimura 1955a, 1955b), the conditional probability (under condition $X = x$ ) is:

$$\mathrm{P}(\omega_1|x) = x_1$$

$$\mathrm{P}(\omega_2|x) = x_3$$

$$\mathrm{P}(\omega_3|x) = x_4$$

$$\mathrm{P}(\omega_4|x) = \gamma_3 x_4 + \gamma_4 x_3$$

(whereby the probability of losing the mutation: $\mathrm{P}(\overline{\omega}_j|x) = 1 - \mathrm{P}(\omega_j|x)$).

Taking into account the independence of corresponding events, we can write:

$$
\left.\begin{aligned}
\mathsf{P}(\omega_{1,1}|x) &= (1-x_1)x_3x_4(\gamma_3x_4+\gamma_4x_3)\\
\mathsf{P}(\omega_{1,2}|x) &= (1-x_1)(1-x_3)x_4(\gamma_3x_4+\gamma_4x_3)\\
\mathsf{P}(\omega_{1,3}|x) &= (1-x_1)x_3(1-x_4)(\gamma_3x_4+\gamma_4x_3)\\
\mathsf{P}(\omega_{1,4}|x) &= (1-x_1)x_3x_4(\gamma_3(1-x_4)+\gamma_4(1-x_3))\\
\mathsf{P}(\omega_{2,2}|x) &= x_1(1-x_3)x_4(\gamma_3x_4+\gamma_4x_3)\\
\mathsf{P}(\omega_{2,3}|x) &= x_1(1-x_3)(1-x_4)(\gamma_3x_4+\gamma_4x_3)\\
\mathsf{P}(\omega_{2,4}|x) &= x_1(1-x_3)x_4(\gamma_3(1-x_4)+\gamma_4(1-x_3))\\
\mathsf{P}(\omega_{3.3}|x) &= x_1x_3(1-x_4)(\gamma_3x_4+\gamma_4x_3)\\
\mathsf{P}(\omega_{3,4}|x) &= x_1x_3(1-x_4)(\gamma_3(1-x_4)+\gamma_4(1-x_3))\\
\mathsf{P}(\omega_{4,4}|x) &= x_1x_3x_4(\gamma_3(1-x_4)+\gamma_4(1-x_3))
\end{aligned}\right| \qquad \text{(S1.3.2)}
$$

Using the full probability formula, we obtain:

$$p_{i,j}^{0}(t) = \mathsf{P}(\omega_{i,j}) = \int_0^1\int_0^1\int_0^1\int_0^1\int_0^1 \mathsf{P}(\omega_{i,j}|x) f^{0}(x)dx_0dx_1dx_2dx_3dx_4. \qquad \text{(S1.3.3)}$$

Substituting the corresponding expressions here and using the formulas for moments given in Section 2, we obtain the polynomial function of $p$ = (2 $N_0(t)$)$^{-1}$ and the exponent $e^{-\tau_k}$, where

$$\tau_k = \int_{\Delta_k} \frac{dt}{2N_k(t)} \quad (k = 1, 2, 3, 4), \qquad \text{(S1.3.4)}$$

and also $e^{-\tau}$, where

$$\tau = \int_t^{t_0} \frac{ds}{2N_0(s)} \qquad \text{(S1.3.5)}$$

Assuming (following Kimura) that $N_0(t)$ and $n_0(t)$ are constant, we get $\tau = (t_0 - t)p$, and calculating:

$$a_{i,j}^{0} = \int_{-\infty}^{t_0} n_0(t)p_{i,j}^{0}(t)dt,$$

and taking into account:

$$\int_{-\infty}^{t_0} e^{-m\tau}\, dt = \frac{1}{mp}.$$

Note also that with constant population size on the branches

$$\begin{aligned}
\tau_1 &= \frac{t_1 - t_0}{2N_1} \\
\tau_2 &= \frac{t_1 - t_0}{2N_2} = \frac{N_1}{N_2}\tau_1 \\
\tau_3 &= \frac{t_2 - t_1}{2N_3} \\
\tau_4 &= \frac{t_2 - t_1}{2N_4} = \frac{N_3}{N_4}\tau_3.
\end{aligned} \qquad \text{(S1.3.6)}$$

Neglecting the terms of order $p^2$ and higher (assuming that $N_0(t) >> 1$) we can write

$$\begin{aligned}
a_{1,1}^0 = n_0 \Big[&\frac{1}{6}\gamma_3\gamma_2^2 e^{-3\tau_2-\tau_4} - \frac{2}{3}\gamma_2 e^{-\tau_2} - \frac{1}{3}\gamma_1\gamma_3\gamma_2 e^{-\tau_1-\tau_2-\tau_4} + (\gamma_2 - \\
&- 1)\gamma_3\gamma_2 e^{-\tau_4} - \frac{1}{3}(3\gamma_2 - 2)\gamma_3\gamma_2 e^{-\tau_2-\tau_4} + \frac{1}{6}\gamma_4\gamma_2 e^{-3\tau_2-\tau_3} - \\
&- \frac{1}{3}\gamma_4\gamma_2 e^{-\tau_2-\tau_3} + \frac{1}{3}\gamma_1 e^{-\tau_1} + \frac{1}{6}\gamma_1(\gamma_1 + 4\gamma_2 - 2)\gamma_3 e^{-\tau_1-\tau_4} - \\
&- \frac{1}{6}\gamma_1\gamma_4 e^{-\tau_1-\tau_2-\tau_3} + \gamma_2\Big]
\end{aligned} \qquad \text{(S1.3.7.1)}$$

$$\begin{aligned}
a_{12}^0 = n_0 \Big[&-\frac{1}{6}\gamma_3\gamma_1^2 e^{-3\tau_1-\tau_4} + \frac{2}{3}\gamma_3\gamma_1 e^{-\tau_1} + \frac{1}{3}(\gamma_1 - 2)\gamma_3\gamma_1 e^{-\tau_1-\tau_4} + \\
&+ \frac{1}{3}\gamma_2\gamma_3\gamma_1 e^{-\tau_1-\tau_2-\tau_4} + \frac{1}{6}\gamma_4\gamma_1 e^{-\tau_1-\tau_2-\tau_3} - \frac{1}{6}\gamma_2^2\gamma_3 e^{-3\tau_2-\tau_4} + \\
&+ \frac{2}{3}\gamma_2\gamma_3 e^{-\tau_2} + \frac{1}{3}(\gamma_2 - 2)\gamma_2\gamma_3 e^{-\tau_2-\tau_4} - \frac{1}{6}\gamma_2\gamma_4 e^{-3\tau_2-\tau_3} + \\
&+ \frac{1}{3}\gamma_2\gamma_4 e^{-\tau_2-\tau_3}\Big]
\end{aligned} \qquad \text{(S1.3.7.2)}$$

$$a_{13}^0 = n_0\left[-\frac{1}{6}\gamma_3\gamma_2^2 e^{-3\tau_2-\tau_4} + \frac{1}{3}\gamma_1\gamma_3\gamma_2 e^{-\tau_1-\tau_2-\tau_4} - (\gamma_2-1)\gamma_3\gamma_2 e^{-\tau_4} + \right.$$
$$+\frac{1}{3}(3\gamma_2-2)\gamma_3\gamma_2 e^{-\tau_2-\tau_4} + \frac{2}{3}\gamma_4\gamma_2 e^{-\tau_2} - \frac{1}{6}\gamma_4\gamma_2 e^{-3\tau_2-\tau_3} - \frac{1}{6}\gamma_1(\gamma_1 +$$
$$+ 4\gamma_2 - 2)\gamma_3 e^{-\tau_1-\tau_4} - \frac{1}{3}\gamma_1\gamma_4 e^{-\tau_1} + \frac{1}{6}\gamma_1\gamma_4 e^{-\tau_1-\tau_2-\tau_3} + \frac{1}{3}(\gamma_2 - \quad \text{(S1.3.7.3)}$$
$$\left. - 2)\gamma_4 e^{-\tau_2-\tau_3} - (\gamma_2-1)\gamma_4\right]$$

$$a_{14}^0 = n_0\left[-\frac{1}{6}\gamma_3\gamma_2^2 e^{-3\tau_2-\tau_4} + \frac{1}{3}\gamma_1\gamma_3\gamma_2 e^{-\tau_1-\tau_2-\tau_4} - (\gamma_2-1)\gamma_3\gamma_2 e^{-\tau_4} + \right.$$
$$+\frac{1}{3}(3\gamma_2-2)\gamma_3\gamma_2 e^{-\tau_2-\tau_4} - \frac{1}{6}\gamma_4\gamma_2 e^{-3\tau_2-\tau_3} + \frac{1}{3}\gamma_4\gamma_2 e^{-\tau_2-\tau_3} - \quad \text{(S1.3.7.4)}$$
$$\left. - \frac{1}{6}\gamma_1(\gamma_1 + 4\gamma_2 - 2)\gamma_3 e^{-\tau_1-\tau_4} + \frac{1}{6}\gamma_1\gamma_4 e^{-\tau_1-\tau_2-\tau_3}\right]$$

$$a_{22}^0 = n_0\left[\frac{1}{6}\gamma_3\gamma_1^2 e^{-3\tau_1-\tau_4} - \frac{2}{3}\gamma_3\gamma_1 e^{-\tau_1} + (\gamma_1-1)\gamma_3\gamma_1 e^{-\tau_4} - \frac{1}{3}(3\gamma_1 - \right.$$
$$- 2)\gamma_3\gamma_1 e^{-\tau_1-\tau_4} - \frac{1}{3}\gamma_2\gamma_3\gamma_1 e^{-\tau_1-\tau_2-\tau_4} - \frac{1}{6}\gamma_4\gamma_1 e^{-\tau_1-\tau_2-\tau_3} +$$
$$+\frac{1}{3}\gamma_2\gamma_3 e^{-\tau_2} + \frac{1}{6}\gamma_2(4\gamma_1+\gamma_2-2)\gamma_3 e^{-\tau_2-\tau_4} + \frac{1}{6}(2\gamma_1 + \quad \text{(S1.3.7.5)}$$
$$\left. + \gamma_2)\gamma_4 e^{-\tau_2-\tau_3} + \gamma_3\gamma_1\right]$$

$$a_{23}^0 = n_0\left[-\frac{1}{6}\gamma_3\gamma_1^2 e^{-3\tau_1-\tau_4} - (\gamma_1-1)\gamma_3\gamma_1 e^{-\tau_4} + \frac{1}{3}(3\gamma_1-2)\gamma_3\gamma_1 e^{-\tau_1-\tau_4} + \right.$$
$$+\frac{1}{3}\gamma_2\gamma_3\gamma_1 e^{-\tau_1-\tau_2-\tau_4} + \frac{1}{6}\gamma_4\gamma_1 e^{-\tau_1-\tau_2-\tau_3} - \frac{1}{6}\gamma_2(4\gamma_1 + \gamma_2 - \quad \text{(S1.3.7.6)}$$
$$\left. - 2)\gamma_3 e^{-\tau_2-\tau_4} - \frac{1}{6}(2\gamma_1+\gamma_2-2)\gamma_4 e^{-\tau_2-\tau_3}\right]$$

$$a_{24}^0 = n_0\left[-\frac{1}{6}\gamma_3\gamma_1^2 e^{-3\tau_1-\tau_4} - (\gamma_1-1)\gamma_3\gamma_1 e^{-\tau_4} + \frac{1}{3}(3\gamma_1-2)\gamma_3\gamma_1 e^{-\tau_1-\tau_4} + \right.$$
$$+\frac{1}{3}\gamma_2\gamma_3\gamma_1 e^{-\tau_1-\tau_2-\tau_4} - \frac{2}{3}\gamma_4\gamma_1 e^{-\tau_1} + \frac{1}{6}\gamma_4\gamma_1 e^{-\tau_1-\tau_2-\tau_3} - \frac{1}{6}\gamma_2(4\gamma_1 + \quad \text{(S1.3.7.7)}$$
$$\left. + \gamma_2 - 2)\gamma_3 e^{-\tau_2-\tau_4} + \frac{1}{3}\gamma_2\gamma_4 e^{-\tau_2} - \frac{1}{6}(2\gamma_1+\gamma_2)\gamma_4 e^{-\tau_2-\tau_3} + \gamma_4\gamma_1\right]$$

$$a_{33}^0 = n_0 \left[ -\frac{1}{3}\gamma_1\gamma_2\gamma_3 e^{-\tau_1-\tau_2-\tau_4} + \frac{1}{6}\gamma_2(4\gamma_1 + 3\gamma_2 - 2)\gamma_3 e^{-\tau_2-\tau_4} + \frac{1}{6}\gamma_1(3\gamma_1 + \right.$$
$$+ 4\gamma_2 - 2)\gamma_3 e^{-\tau_1-\tau_4} + \frac{1}{3}\gamma_1\gamma_4 e^{-\tau_1} - \frac{1}{6}\gamma_1\gamma_4 e^{-\tau_1-\tau_2-\tau_3} + \frac{1}{3}\gamma_2\gamma_4 e^{-\tau_2} + \quad \text{(S1.3.7.8)}$$
$$\left. + \frac{1}{6}(2\gamma_1 + \gamma_2 - 2)\gamma_4 e^{-\tau_2-\tau_3} \right]$$

$$a_{34}^0 = n_0 \left[ \frac{1}{3}\gamma_1\gamma_3 e^{-\tau_1} + \frac{1}{3}\gamma_2\gamma_3 e^{-\tau_2} + \frac{1}{3}\gamma_1\gamma_2\gamma_3 e^{-\tau_1-\tau_2-\tau_4} - \frac{1}{6}\gamma_2(4\gamma_1 + 3\gamma_2 - \right.$$
$$- 2)\gamma_3 e^{-\tau_2-\tau_4} - \frac{1}{6}\gamma_1(3\gamma_1 + 4\gamma_2 - 2)\gamma_3 e^{-\tau_1-\tau_4} + \frac{1}{6}\gamma_1\gamma_4 e^{-\tau_1-\tau_2-\tau_3} - \quad \text{(S1.3.7.9)}$$
$$\left. - \frac{1}{6}(2\gamma_1 + \gamma_2 - 2)\gamma_4 e^{-\tau_2-\tau_3} \right]$$

$$a_{44}^0 = n_0 \left[ -\frac{1}{3}\gamma_1\gamma_2\gamma_3 e^{-\tau_1-\tau_2-\tau_4} + \frac{1}{6}\gamma_2(4\gamma_1 + 3\gamma_2 - 2)\gamma_3 e^{-\tau_2-\tau_4} + \right.$$
$$+ \frac{1}{6}\gamma_1(3\gamma_1 + 4\gamma_2 - 2)\gamma_3 e^{-\tau_1-\tau_4} - \frac{1}{6}\gamma_1\gamma_4 e^{-\tau_1-\tau_2-\tau_3} + \frac{1}{6}(2\gamma_1 + \quad \text{(S1.3.7.10)}$$
$$\left. + \gamma_2)\gamma_4 e^{-\tau_2-\tau_3} \right]$$

##### 3.1.2. Branch $B_1$

For a retrotransposon insertion on the branch $B_1$ at $t \epsilon (t_0, t_1)$, its frequencies at branches $B_0$, $B_2$, and $B_3$ are null. Then as in terms 3.1.1, we consider the random vector $\mathcal{X} = (0, X_1, 0, 0, X_4)$.

Then, introducing the (non-random) vector $x = (0, x_1, 0, 0, x_4)$ we see that the probability density function of the new vector $\mathcal{X}$

$$f^1(x) = u_1(t, p, t_1, x_1)u_4(t_1, \gamma_1 x_1, t_2, x_4), \quad \text{(S1.3.8)}$$

where $p = (2\, N_1(t))^{-1}$

Note, that if $i$ or $j$ does not equal 2, then all of such $\mathsf{P}(\omega_{i,j}) = 0$. Next, substituting in formulas (S1.3.2) $x_2 = x_3 = 0$:

$$\left. \begin{aligned} \mathsf{P}(\omega_{1,2}|x) &= (1 - x_1)x_4^2\gamma_3 \\ \mathsf{P}(\omega_{2,2}|x) &= x_1 x_4^2\, \gamma_3 \\ \mathsf{P}(\omega_{2,3}|x) &= x_1(1 - x_4)\gamma_3 x_4 \\ \mathsf{P}(\omega_{2,4}|x) &= x_1 x_4(\gamma_3(1 - x_4) + \gamma_4) \end{aligned} \right| \quad \text{(S1.3.9)}$$

Using the full probability formula, we obtain:

$$p_{i,j}^{1}(t)=\mathrm{P}(\omega_{i,j})=\int_0^1\int_0^1 \mathrm{P}(\omega_{i,j}|x)f^{1}(x)dx_1dx_4. \qquad \text{(S1.3.10)}$$

Proceeding similarly to 3.1.1 we write

$$\left.\begin{aligned}
a_{12}^1 &= n_1\left[\frac{1}{6}e^{-\tau_4}\gamma_3(-6\tau_1+e^{-3\tau_1}-9e^{-\tau_1}+8)\gamma_1^3+e^{-\tau_4}\gamma_3(\tau_1+e^{-\tau_1}-1)\gamma_1^2\right]\\
a_{22}^1 &= n_1\left[\frac{1}{6}e^{-\tau_4}\gamma_1^3\gamma_3(6\tau_1-e^{-3\tau_1}+9e^{-\tau_1}-8)-(-1+e^{-\tau_4})\gamma_1^2\gamma_3(\tau_1+\right.\\
&\qquad \left.+e^{-\tau_1}-1)\right]\\
a_{23}^1 &= n_1\left[\frac{1}{6}e^{-\tau_4}(2+e^{-3\tau_1}-3e^{-\tau_1})\gamma_3\gamma_1^3+((-1+e^{-\tau_1})(-1+e^{-\tau_4})\gamma_3+\right.\\
&\qquad \left.+e^{-\tau_4}\gamma_2(\tau_1+e^{-\tau_1}-1)\gamma_3)\gamma_1^2+\gamma_2\gamma_3(\tau_1-e^{-\tau_4}\tau_1)\gamma_1\right]\\
a_{24}^1 &= n_1\left[\frac{1}{6}e^{-\tau_4}\gamma_3(-6\tau_1+e^{-3\tau_1}-9e^{-\tau_1}+8)\gamma_1^3+(e^{-\tau_4}\gamma_3(\tau_1+e^{-\tau_1}-1)+\right.\\
&\qquad \left.+\gamma_4(\tau_1+e^{-\tau_1}-1))\gamma_1^2\right]
\end{aligned}\right| \qquad \text{(S1.3.11)}$$

##### 3.1.3. Branch $B_2$

For a retrotransposon insertion that takes place on the branch $B_2$ at $t\epsilon(t_0,t_1)$, its frequencies at branches $B_0$ and $B_1$ are null. Then, as in section 3.1.1, we consider the vector:

$$\mathcal{X} = (X_2, X_3, X_4)$$

Now, introducing vector $x = (x_2, x_3, x_4)$ we see that the probability density function of vector $\mathcal{X}$ is:

$$f^2(x) = u_2(t,p,t_1,x_2)u_3(t_1,x_2,t_2,x_3)u_4(t_1,\gamma_2x_2,t_2,x_4), \qquad \text{(S1.3.12)}$$

where $p = (2\,N_2(t))^{-1}$.

Note, that if $i$ or $j$ does not equal 2, then all of such $\mathrm{P}(\omega_{i,j}) = 0$. Next, substituting in formulas (S1.3.2) $x_1 = 0$ we have:

$$\begin{aligned} P(\omega_{1,1}|x) &= x_3 x_4(\gamma_3 x_4 + \gamma_4 x_3) \\ P(\omega_{1,2}|x) &= (1-x_3)x_4(\gamma_3 x_4 + \gamma_4 x_3) \\ P(\omega_{1,3}|x) &= x_3(1-x_4)(\gamma_3 x_4 + \gamma_4 x_3) \\ P(\omega_{1,4}|x) &= x_3 x_4(\gamma_3(1-x_4) + \gamma_4(1-x_3)) \end{aligned} \tag{S1.3.13}$$

Using the full probability formula, we obtain:

$$p_{i,j}^2(t) = P(\omega_{i,j}) = \int_0^1\int_0^1\int_0^1 P(\omega_{i,j}|x) f^2(x) dx_2 dx_3 dx_4. \tag{S1.3.14}$$

Proceeding similarly to S1.3.1.1 we can write

$$\begin{aligned} a_{12}^1 &= n_1\left[\frac{1}{6}e^{-\tau_4}\gamma_3(-6\tau_1 + e^{-3\tau_1} - 9e^{-\tau_1} + 8)\gamma_1^3 + e^{-\tau_4}\gamma_3(\tau_1 + e^{-\tau_1} - 1)\gamma_1^2\right] \\ a_{22}^1 &= n_1\left[\frac{1}{6}e^{-\tau_4}\gamma_1^3\gamma_3(6\tau_1 - e^{-3\tau_1} + 9e^{-\tau_1} - 8) - (-1 + e^{-\tau_4})\gamma_1^2\gamma_3(\tau_1 + \right. \\ &\left. + e^{-\tau_1} - 1)\right] \\ a_{23}^1 &= n_1\left[\frac{1}{6}e^{-\tau_4}(2 + e^{-3\tau_1} - 3e^{-\tau_1})\gamma_3\gamma_1^3 + ((-1 + e^{-\tau_1})(-1 + e^{-\tau_4})\gamma_3 + \right. \\ &\left. + e^{-\tau_4}\gamma_2(\tau_1 + e^{-\tau_1} - 1)\gamma_3)\gamma_1^2 + \gamma_2\gamma_3(\tau_1 - e^{-\tau_4}\tau_1)\gamma_1\right] \\ a_{24}^1 &= n_1\left[\frac{1}{6}e^{-\tau_4}\gamma_3(-6\tau_1 + e^{-3\tau_1} - 9e^{-\tau_1} + 8)\gamma_1^3 + (e^{-\tau_4}\gamma_3(\tau_1 + e^{-\tau_1} - 1) + \right. \\ &\left. + \gamma_4(\tau_1 + e^{-\tau_1} - 1))\gamma_1^2\right] \end{aligned} \tag{S1.3.15}$$

##### 3.1.4. Branch $B_3$

For a retrotransposon insertion on the branch $B_3$ at $t\epsilon(t_1, t_2)$, its frequencies at branches $B_0$, $B_1$, $B_2$, and $B_4$ are null. Then, as in section 3.1.1, we consider the one-dimensional vector $\mathcal{X} = X_3$.

It then follows that all $P(\omega_{i,j})$ (and $a_{i,j}^3$ correspondents) except $P(\omega_{1,3})$ are null. Then assuming in formulas (S1.3.1) $x_1 = x_2 = x_4 = 0$ we have

$$P(\omega_{1,3}|x) = \gamma_4 x_3^2 \tag{S1.3.16}$$

Using the full probability formula, we obtain:

$$p_{1,3}^{3}(t) = \int_0^1 \mathrm{P}(\omega_{1,3}|x) f^3(x) dx_3 = \gamma_4 m_2(t, p, t_2). = \gamma_4 \left(p^2 e^{-\tau} + p(1 - e^{-\tau})\right) \qquad (S1.3.17)$$

where

$$\tau = \int_t^{t_2} \frac{ds}{2N_3(s)} = \frac{t_2 - t}{2N_3}.$$

Hence according to (S1.1.5):

$$a_{1,3}^{3} = \int_{t_1}^{t_2} n_3 p_{1,3}^{3}(t) dt,$$

and neglecting the terms of order $p^2$, we can write:

$$a_{13}^{3} = n_3 \gamma_4 (\tau_3 + e^{-\tau_3} - 1) \qquad (S1.3.18)$$

##### 3.1.5. Branch $B_4$

For a retrotransposon insertion on the branch $B_4$ at $t \epsilon (t_1, t_2)$, its frequencies at branches $B_0$, $B_1$, $B_2$, and $B_3$ are null. Then, as in section 3.1.1, we consider the one-dimensional vector $\mathcal{X} = X_4$.

Then, for $x = x_4$ we see that the probability density function of vector $\mathcal{X}$

$$f^4(x) = u_4(t, p, t_2, x_4), \qquad (S1.3.19)$$

where $p = (2\,N_4(t))^{-1}$

It then follows that all $\mathrm{P}(\omega_{i,j})$ (and $a_{i,j}^{4}$ correspondents) except $\mathrm{P}(\omega_{1,2})$ are null. Then assuming in formulas (S1.3.2) $x_1 = x_2 = x_3 = 0$ we have

$$\mathrm{P}(\omega_{1,2}|x) = \gamma_3 x_4^2 \qquad (S1.3.20)$$

From here, as in section 3.1.1, we obtain:

$$a_{12}^{4} = n_4 \gamma_3 (\tau_4 + e^{-\tau_4} - 1) \qquad (S1.3.21)$$

#### 3.2. Hybridization topology 2H2

As in the section 1.3.1, we consider five branches: $B_0$, $B_1$, $B_2$, $B_3$, $B_4$.

##### 3.2.1. Branch $B_0$

For a retrotransposon insertion that takes place on the branch $B_0$ at $t < t_0$, its frequencies at the terminal ends of the respective branches form the random vector $\mathcal{X} = (X_0, X_1, X_2, X_3, X_4)$.

Then, introducing the (non-random) vector $x = (x_0, x_1, x_2, x_3, x_4)$, we see that the probability density function of vector $\mathcal{X}$

$$f^0(x) = u_0(t,p,t_0,x_0)u_1(t_0,x_0,t_1,x_1)u_2(t_0,x_0,t_1,x_2) \\ u_3(t_1,x_2,t_2,x_3)u_4(t_1,x_1,t_2,x_4), \qquad \text{(S1.3.22)}$$

where $p = (2\,N_0(t))^{-1}$, and the index of the function $u$ indicates the tree branch number.

Let $\omega_j$ = an event whereby this insertion is fixed in lineage $A_j$. Then, as in section 3.1.1 we have

$$\mathrm{P}(\omega_1|x) = x_4$$

$$\mathrm{P}(\omega_2|x) = x_3$$

$$\mathrm{P}(\omega_3|x) = \gamma_1 x_1 + \gamma_2 x_2$$

$$\mathrm{P}(\omega_4|x) = \gamma_3 x_4 + \gamma_4 x_3$$

(whereby the probability of losing the mutation: $\mathrm{P}(\overline{\omega}_j|x) = 1 - \mathrm{P}(\omega_j|x)$).

Taking into account the independence of similar events we can write:

$$\left.\begin{aligned}
\mathrm{P}(\omega_{1,1}|x) &= x_3(1-x_4)(\gamma_1 x_1 + \gamma_2 x_2)(\gamma_3 x_4 + \gamma_4 x_3)\\
\mathrm{P}(\omega_{1,2}|x) &= (1-x_3)(1-x_4)(\gamma_1 x_1 + \gamma_2 x_2)(\gamma_3 x_4 + \gamma_4 x_3)\\
\mathrm{P}(\omega_{1,3}|x) &= x_3(1-x_4)(\gamma_1(1-x_1) + \gamma_2(1-x_2))(\gamma_3 x_4 + \gamma_4 x_3)\\
\mathrm{P}(\omega_{1,4}|x) &= x_3(1-x_4)(\gamma_1 x_1 + \gamma_2 x_2)(\gamma_3(1-x_4) + \gamma_4(1-x_3))\\
\mathrm{P}(\omega_{2,2}|x) &= (1-x_3)x_4(\gamma_1 x_1 + \gamma_2 x_2)(\gamma_3 x_4 + \gamma_4 x_3)\\
\mathrm{P}(\omega_{2,3}|x) &= (1-x_3)x_4(\gamma_1(1-x_1) + \gamma_2(1-x_2))(\gamma_3 x_4 + \gamma_4 x_3)\\
\mathrm{P}(\omega_{2,4}|x) &= (1-x_3)x_4(\gamma_1 x_1 + \gamma_2 x_2)(\gamma_3(1-x_4) + \gamma_4(1-x_3))\\
\mathrm{P}(\omega_{3.3}|x) &= x_3 x_4(\gamma_1(1-x_1) + \gamma_2(1-x_2))(\gamma_3 x_4 + \gamma_4 x_3)\\
\mathrm{P}(\omega_{3,4}|x) &= x_3 x_4(\gamma_1(1-x_1) + \gamma_2(1-x_2))(\gamma_3(1-x_4) + \gamma_4(1-x_3))\\
\mathrm{P}(\omega_{4,4}|x) &= x_3 x_4(\gamma_1 x_1 + \gamma_2 x_2)(\gamma_3(1-x_4) + \gamma_4(1-x_3))
\end{aligned}\right| \qquad \text{(S1.3.23)}$$

Further, as in section 3.1.1, we can derive:

$$a_{11}^0 = n_0\left[\frac{1}{6}(\gamma_2\gamma_4 e^{-\tau_3}(e^{-3\tau_2} - 3e^{-\tau_2} + 2) + \gamma_2\gamma_3 e^{-\tau_4}(\gamma_2(-6\tau_2 + e^{-3\tau_2} - \right. \\ - 9e^{-\tau_2} + 8) + 6(\tau_2 + e^{-\tau_2} - 1)))\frac{1}{6}(\gamma_2\gamma_4 e^{-\tau_3}(e^{-3\tau_2} - 3e^{-\tau_2} + \\ + 2) + \gamma_2\gamma_3 e^{-\tau_4}(\gamma_2(-6\tau_2 + e^{-3\tau_2} - 9e^{-\tau_2} + 8) + 6(\tau_2 + e^{-\tau_2} - \\ \left. - 1)))\right] \qquad \text{(S1.3.24.1)}$$

$$a_{12}^{0} = n_0 \left[ \frac{1}{6} (\gamma_2 (\gamma_3 e^{-\tau_1-\tau_2-\tau_4} - \gamma_4 e^{-3\tau_2-\tau_3} + 2\gamma_4 e^{-\tau_2-\tau_3}) + \right.$$
$$\left. + \gamma_1 (\gamma_3 (-e^{-3\tau_1-\tau_4}) + 2\gamma_3 e^{-\tau_1-\tau_4} + \gamma_4 e^{-\tau_1-\tau_2-\tau_3})) \right] \quad \text{(S1.3.24.2)}$$

$$a_{13}^{0} = n_0 \left[ \frac{1}{6} (\gamma_1 \gamma_3 e^{-\tau_1-\tau_4} + \gamma_2 \gamma_3 e^{-\tau_1-\tau_2-\tau_4} + \gamma_1 \gamma_4 (-2e^{-\tau_1} - 4e^{-\tau_2-\tau_3} + \right.$$
$$\left. + e^{-\tau_1-\tau_2-\tau_3} + 6) + \gamma_2 \gamma_4 (4e^{-\tau_2} - e^{-3\tau_2-\tau_3} - 2e^{-\tau_2-\tau_3})) \right] \quad \text{(S1.3.24.3)}$$

$$a_{14}^{0} = n_0 \left[ \frac{1}{6} (\gamma_2 \gamma_3 (-4e^{-\tau_2} - 2e^{-\tau_1-\tau_4} + e^{-\tau_1-\tau_2-\tau_4} + 6) - \gamma_2 \gamma_4 (e^{-3\tau_2-\tau_3} - \right.$$
$$\left. - 2e^{-\tau_2-\tau_3}) + \gamma_1 (2\gamma_3 e^{-\tau_1} - \gamma_3 e^{-\tau_1-\tau_4} + \gamma_4 e^{-\tau_1-\tau_2-\tau_3})) \right] \quad \text{(S1.3.24.4)}$$

$$a_{22}^{0} = n_0 \left[ \frac{1}{6} (\gamma_1 \gamma_3 (-4e^{-\tau_1} + e^{-3\tau_1-\tau_4} - 2e^{-\tau_1-\tau_4} + 6) + \gamma_1 \gamma_4 (2e^{-\tau_2-\tau_3} - \right.$$
$$\left. - e^{-\tau_1-\tau_2-\tau_3}) + \gamma_2 (2\gamma_3 e^{-\tau_2} - \gamma_3 e^{-\tau_1-\tau_2-\tau_4} + \gamma_4 e^{-\tau_2-\tau_3})) \right] \quad \text{(S1.3.24.5)}$$

$$a_{23}^{0} = n_0 \left[ \frac{1}{6} (\gamma_1 \gamma_3 (4e^{-\tau_1} - e^{-3\tau_1-\tau_4} - 2e^{-\tau_1-\tau_4}) + \gamma_2 \gamma_3 (-2e^{-\tau_2} - \right.$$
$$\left. - 4e^{-\tau_1-\tau_4} + e^{-\tau_1-\tau_2-\tau_4} + 6) + \gamma_1 \gamma_4 e^{-\tau_1-\tau_2-\tau_3} + \gamma_2 \gamma_4 e^{-\tau_2-\tau_3}) \right] \quad \text{(S1.3.24.6)}$$

$$a_{24}^{0} = n_0 \left[ \frac{1}{6} (\gamma_1 \gamma_3 (-(e^{-3\tau_1-\tau_4} - 2e^{-\tau_1-\tau_4})) + \gamma_1 \gamma_4 (-4e^{-\tau_1} - 2e^{-\tau_2-\tau_3} + \right.$$
$$+ e^{-\tau_1-\tau_2-\tau_3} + 6) + \gamma_2 (\gamma_3 e^{-\tau_1-\tau_2-\tau_4} + 2\gamma_4 e^{-\tau_2} -$$
$$- \gamma_4 e^{-\tau_2-\tau_3})) \frac{1}{6} (\gamma_1 \gamma_3 (-(e^{-3\tau_1-\tau_4} - 2e^{-\tau_1-\tau_4})) + \gamma_1 \gamma_4 (-4e^{-\tau_1} - \quad \text{(S1.3.24.7)}$$
$$- 2e^{-\tau_2-\tau_3} + e^{-\tau_1-\tau_2-\tau_3} + 6) + \gamma_2 (\gamma_3 e^{-\tau_1-\tau_2-\tau_4} + 2\gamma_4 e^{-\tau_2} -$$
$$\left. - \gamma_4 e^{-\tau_2-\tau_3})) \right]$$

$$a_{33}^{0} = n_0 \left[ \frac{1}{6} (\gamma_2 (\gamma_3 (-e^{-\tau_1-\tau_2-\tau_4}) - \gamma_4 e^{-\tau_2-\tau_3} + 2e^{-\tau_2}) + \gamma_1 (\gamma_3 (-e^{-\tau_1-\tau_4}) - \right.$$
$$\left. - \gamma_4 e^{-\tau_1-\tau_2-\tau_3} + 2e^{-\tau_1})) \right] \quad \text{(S1.3.24.8)}$$

$$a_{34}^{0} = n_0 \left[ \frac{1}{6} (\gamma_1 \gamma_3 e^{-\tau_1-\tau_4} + \gamma_2 \gamma_3 e^{-\tau_1-\tau_2-\tau_4} + \gamma_1 \gamma_4 e^{-\tau_1-\tau_2-\tau_3} + \gamma_2 \gamma_4 e^{-\tau_2-\tau_3}) \right] \quad \text{(S1.3.24.9)}$$

$$a_{44}^{0} = n_0 \left[\frac{1}{6}(\gamma_2(2\gamma_3 e^{-\tau_1-\tau_4} - \gamma_3 e^{-\tau_1-\tau_2-\tau_4} + \gamma_4 e^{-\tau_2-\tau_3}) + \gamma_1(\gamma_3 e^{-\tau_1-\tau_4} + \right.$$
$$\left. + 2\gamma_4 e^{-\tau_2-\tau_3} - \gamma_4 e^{-\tau_1-\tau_2-\tau_3}))\right] \quad \text{(S1.3.24.10)}$$

Here $\tau_1, \tau_2, \tau_3, \tau_4$ are defined according to (S1.3.6).

##### 3.2.2. Branch $B_1$

For a retrotransposon insertion on the branch $B_1$ at $t\epsilon(t_0, t_1)$, its frequencies at branches $B_0$, $B_2$, and $B_3$ are null. Then, as in section 3.1.1, we consider the random vector $\mathcal{X} = (0, X_1, 0, 0, X_4)$.

Then, introducing the (non-random) vector $x = (0, x_1, 0, 0, x_4)$ we see that the probability density function of the new vector $\mathcal{X}$ is:

$$f^1(x) = u_1(t, p, t_1, x_1)u_4(t_1, x_1, t_2, x_4), \quad \text{(S1.3.25)}$$

where $p = (2\, N_1(t))^{-1}$

Note, that if $i$ or $j$ does not equal 2, then all $\mathrm{P}(\omega_{i,j}) = 0$.

If we then substitute in formulas (S1.3.2) $x_2 = x_3 = 0$:

$$\mathrm{P}(\omega_{1,2}|x) = x_1(1 - x_4)x_4\gamma_1\gamma_3$$
$$\mathrm{P}(\omega_{2,2}|x) = x_1 x_4^2\, \gamma_1\gamma_3$$
$$\mathrm{P}(\omega_{2,3}|x) = (\gamma_1(1 - x_1) + \gamma_2)x_4^2\gamma_3$$
$$\mathrm{P}(\omega_{2,4}|x) = x_1 x_4(\gamma_3(1 - x_4) + \gamma_4)\gamma_1$$
(S1.3.26)

Proceeding as in section S1.3.1.2 we can write:

$$a_{12}^{1} = n_1 \frac{1}{6} e^{-\tau_4}(2 + e^{-3\tau_1} - 3e^{-\tau_1})\gamma_1\gamma_3$$
$$a_{22}^{1} = n_1 \frac{1}{6} e^{-\tau_4}(2 + e^{-3\tau_1} - 3e^{-\tau_1})\gamma_1\gamma_3$$
$$a_{23}^{1} = n_1 \left[\frac{1}{6}(-1 + e^{-\tau_1})(e^{-\tau_4}(4 + e^{-2\tau_1} + e^{-\tau_1}) - 6)\gamma_1\gamma_3 + \gamma_2(e^{-\tau_4}(-1 + \right.$$
$$\left. + e^{-\tau_1}) + \tau_1)\gamma_3\right]$$
$$a_{24}^{1} = n_1\gamma_1 \left[\frac{1}{6} e^{-\tau_4}(2 + e^{-3\tau_1} - 3e^{-\tau_1})\gamma_3 + \gamma_4(\tau_1 + e^{-\tau_1} - 1)\right]$$
(S1.3.27)

##### 3.2.3. Branch $B_2$

As we can see by analyzing the 2H2 model, the results for branch $B_2$ can be obtained from the ones for branch $B_2$ if there is a change in the indices $(i, j)$ 1 ↔ 2 and 3 ↔ 4.

Thus, we have:

$$\begin{aligned}
a_{11}^2 &= n_2\gamma_2\gamma_4\left[-\frac{1}{6}e^{-\tau_2-\tau_3}(3-e^{-2\tau_2})+e^{-\tau_2}-\frac{e^{-\tau_3}}{3}+\tau_2-1\right] \\
a_{12}^2 &= n_2\frac{1}{6}e^{-\tau_3}(2+e^{-3\tau_2}-3e^{-\tau_2})\gamma_2\gamma_4 \\
a_{13}^2 &= n_2\left[\frac{1}{6}(-1+e^{-\tau_2})(e^{-\tau_3}(4+e^{-2\tau_2}+e^{-\tau_2})-6)\gamma_2\gamma_4+\gamma_1(e^{-\tau_3}(-1+\right. \\
&\quad \left.+e^{-\tau_2})+\tau_2)\gamma_4\right] \\
a_{14}^2 &= n_2\gamma_2\left[\frac{1}{6}e^{-\tau_3}(2+e^{-3\tau_2}-3e^{-\tau_2})\gamma_4+\gamma_3(\tau_2+e^{-\tau_2}-1)\right]
\end{aligned} \qquad \text{(S1.3.28)}$$

##### 3.2.4. Branch $B_3$

Comparing models 2H1 and 2H2, all the arguments of section S1.3.1.4 remain valid. Therefore, the formula (S1.3.19) is still correct:

$$a_{13}^3 = n_3\gamma_4(\tau_3+e^{-\tau_3}-1) \qquad \text{(S1.3.29)}$$

(all other $a_{i,j}^3 = 0$).

##### 3.2.5. Branch $B_4$

Comparing models 2H1 and 2H2, when i or j are not equal to 2 or 3, then $a_{i,j}^4 = 0$, and $a_{23}^4$ is defined by the right side of the equation (S1.3.22):

$$a_{23}^4 = n_4\gamma_3(\tau_4+e^{-\tau_4}-1) \qquad \text{(S1.3.30)}$$

#### 4. Evaluation of test data

##### 4.1. Parameter estimation

According to equation (S1.1.3), the parameters of the Poisson-distributed, random variables $\xi_{i,j}$ can be found as:

$$a_{i,j} = \sum_{k=0}^{4} a_{i,j}^{k} , \quad \text{(S1.4.1)}$$

where $a_{i,j}^{k}$ are defined with equations (S1.3.7), (S1.3.11), (S1.3.15), (S1.3.19), (S1.3.22) for model 2H1 and with (S1.3.25), (S1.3.28), (S1.3.29), (S1.3.30), (S1.3.31) for model 2H2.

However, because in practice, not all diagnostic markers can be identified (due to limited screening, imperfect 2-way or multiway alignments, or low-quality genome data) we introduce β as the probability that the current locus of insertion can be recognized in all species. It can then be argued that the observed values of $\xi_{i,j},$ as well as the parameters of the Poisson-distributed, random variables, can be obtained from the ancestral parameters by multiplying them by β. In fact, we can use the above equations, taking into account that $n_k$ in equation (S1.1.1) is:

$$n_k = 2N_k \alpha_k \beta \quad \text{(S1.4.2)}$$

(Recall that, when we obtained the above equations, it was assumed that $N_k$ and $\alpha_k$ were constant on the corresponding branches).

We also note that because branches $B_1$ and $B_2$ (as well as $B_3$ and $B_4$) belong to the same period, it is reasonable to assume that the probabilities of mutations are the same for both:

$$\alpha_1 = \alpha_2 , \;\; \alpha_3 = \alpha_4 \;\; . \quad \text{(S1.4.3)}$$

Then we have:

$$\frac{N_1}{N_2} = \frac{n_1}{n_2} \;\; , \qquad \frac{N_3}{N_4} = \frac{n_3}{n_4} \;\; . \quad \text{(S1.4.4)}$$

And, by (S1.3.6):

$$\tau_2 = \frac{n_1}{n_2} \tau_1 \, , \qquad \tau_4 = \frac{n_3}{n_4} \tau_3 \, . \quad \text{(S1.4.5)}$$

Hence, denoting

$$r_k = \frac{n_k}{n_0} \quad (1 \leq k \leq 4), \quad \text{(S1.4.6)}$$

we can write:

$$n_k = r_k n_0 \quad (1 \leq k \leq 4), \quad \text{(S1.4.7)}$$

and

$$\tau_2 = \frac{r_1}{r_2} \tau_1 \, , \qquad \tau_4 = \frac{r_3}{r_4} \tau_3 \, . \quad \text{(S1.4.8)}$$

Thus, if coefficients $r_k$ are fixed, the parameters $a_{i,j}$ depend on five values: $n_0, T_1, T_3, \gamma_1,$ and $\gamma_3$ (noting that $\gamma_2 = 1 - \gamma_1, \;\gamma_4 = 1 - \gamma_3$ ) where,

$$T_1 = r_1 \tau_1 = r_2 \tau_2, \qquad T_3 = r_3 \tau_3 = r_4 \tau_4 . \quad \text{(S1.4.9)}$$

Thus, for given values of $r_k$ $(1 \leq k \leq 4)$, $a_{i,j}$ can be written in the form:

$$a_{i,j} = n_0 \bar{a}_{i,j}(T_1, T_3, \gamma_1, \gamma_3), \quad \text{(S1.4.10)}$$

As we noted above, in each of the two models, one can consider different permutations of the lineages $A_1$, $A_2$, $A_3$, $A_4$. To do so, let $S_4$ be a set of all 4-permutations. Then for any permutation $\sigma = (\sigma_1, \sigma_2, \sigma_3, \sigma_4) \in S_4$ we consider the corresponding scheme with substitution: $A_i \to A_{\sigma_i}$ $(1 \le i \le 4)$. In this case, in equation S1.4.1, we need to replace $a_{i,j}$ with $a_{\sigma_i,\sigma_j}$.

Setting $x = \{x_{i,j}\}_{1\le i,j\le 4}$, where $x_{i,j}$ are arbitrary nonnegative integers with $x_{i,j} = x_{j,i}$, we can write the likelihood function as follows:

$$l(x;\theta) = l(x;\mathrm{m},\sigma,\nu) = \prod_{1\le i\le j\le 4} (a_{i,j})^{x_{\sigma_i,\sigma_j}} e^{-a_{i,j}}\,, \qquad \text{(S1.4.11)}$$

where $\theta = (\mathrm{m},\sigma,\nu)$ is the model parameters vector; $a_{i,j} = a_{i,j}(\mathrm{m},\nu)$; m is a type of the model (2H1 or 2H2), and $\nu = (n_0, T_1, T_3, \gamma_1, \gamma_3)$.

If $x_{i,j}$ are observed values of random variables $\xi_{i,j}$, the estimates of the parameters for the corresponding model can be derived by maximizing $l(x;\theta)$ on the set $\Theta = \mathrm{M} \times S_4 \times V$, where

$$\mathrm{M} = \{2\mathrm{H}1, 2\mathrm{H}2\}$$

$$\mathrm{V} = \{\nu = (n_0, T_1, T_3, \gamma_1, \gamma_3): n_0 \geqslant 0, T_1 \geqslant 0, T_3 \geqslant 0, 0 \leqslant \gamma_1 \leqslant 1, 0 \leqslant \gamma_3 \leqslant 1\}$$

Thus

$$L(x|\Theta) = \sup_{\theta\in\Theta} l(x;\theta) = \max_{\mathrm{m}\in\mathrm{M}} \max_{\sigma\in S_4} \sup_{\nu\in V} l(x;\mathrm{m},\sigma,\nu)$$

or

$$L(x;\Theta) = \max_{\mathrm{m}\in\mathrm{M}} L(x;\mathrm{m}|S_4\times \mathrm{V}) = \max\{L(x;2\mathrm{H}1|S_4\times\mathrm{V}), L(x;2\mathrm{H}2|S_4\times\mathrm{V})\},$$

where

$$L(x;\mathrm{m}|S_4\times\mathrm{V}) = \max_{\sigma\in S_4} L(x;\mathrm{m},\sigma|\mathrm{V}), \text{ and}$$

$$L(x;\mathrm{m},\sigma|\mathrm{V}) = \sup_{\nu\in\mathrm{V}} l(x;\mathrm{m},\sigma,\nu).$$

Practically, for each of the two types of models (m = 2H1 or m = 2H2) and for each permutation $\sigma$, we must find likelihood values $L(x;\mathrm{m},\sigma|\mathrm{V})$ and then select the highest of them.

Note. As can be seen from Fig. 2, if we interchange $A_1$ and $A_2$, we will get the same model as when the value of $\gamma_2$ replaces that of $\gamma_1$, and $\gamma_3$ replaces $\gamma_4$, and the likelihood values $L(x;\mathrm{m},\sigma|\mathrm{V})$ are also identical. Thus, for a model of type 2H2, we can restrict ourselves to only twelve permutations, for example, with the condition: $\sigma_1 < \sigma_2$.

#### 4.2. Special cases

We obtained special cases of our models by fixing some of the parameters $T_1$, $T_3$, $\gamma_1$, and $\gamma_3$. The maximum likelihood estimates for the parameters were found by maximizing $l(x;\mathrm{m},\sigma,\nu)$ on the corresponding subset of the set V. For clarity, we set: V = *TTgg*. Fixing, for example, $\gamma_3 = 0$, we set:

$$TTg0 = \{\nu = (n_0, T_1, T_3, \gamma_1, \gamma_3): n_0 \geqslant 0, T_1 \geqslant 0, T_3 \geqslant 0, 0 \leqslant \gamma_1 \leqslant 1, \gamma_3 = 0\} \subset \mathrm{V},$$

and we calculated $L(x;\mathrm{m},\sigma|TTg0) = \sup_{\nu\in V} l(x;\mathrm{m},\sigma,\nu)$.

Similarly, we considered other subsets. Mathematically replacing all terms of $T$ with zero and $\gamma$ with zero or one, we constructed 2·2·3·3 = 36 combinations. However, some of them lead to equivalent models. Particularly, restricting $T_1 = 0$, leads to the independency of $l(x;\mathrm{m},\sigma,\nu)$ from $\gamma_1$, then 0*Tgg* = 0*T*0*g* = 0*T*1*g* = 0*Tng* (here the symbol *n* indicates no dependence on the corresponding parameter; for example, 00*nn* replacing 00*gg* = 000*g* = 001*g* = 00*g*0 = 00*g*1 = 0000 = 0001 = 0010 = 0011).

Further analysis shows that for model 2H1, with a fixed permutation $\sigma = (\sigma_1, \sigma_2, \sigma_3, \sigma_4)$, there are only 20 different cases; for model 2H2 there are 22 different cases (See Table 1). Thus, mathematically we have 24·20 + 12·22 = 744 variants. However, among them, one can also find topologically equivalent ones. So, for example, the following are equivalent:

$$(2H1{,}1234, TTg1), (2H1{,}1243, TTg1), (2H1{,}2134, TTg1), (2H1{,}2143, TTg1)$$

(in the future we will classify these as 1H4);

as are these

$$(2H1{,}1234, TTg0), (2H1{,}1432, TTg0), (2H2{,}1234, TTg0),$$

$$(2H2{,}1432, TTg0), (2H2{,}2134, TTg1), (2H2{,}4132, TTg1)$$

(in the future we will classify them as 1H1) etc.

**Table 1. Simplified cases for topologies 2H1 and 2H2.**

| | | | |
|---|---|---|---|
| 1 | TTgg | 11 | T0g1 |
| 2 | TTg1 | 12 | T0g0 |
| 3 | TTg0 | 13 | T01g |
| 4 | TT1g | 14 | 0Tng |
| 5 | TT0g | 15 | T00n |
| 6 | T0gg | 16 | T010 |
| 7 | TT01 | 17 | T011 |
| 8 | TT10 | 18 | 0Tn1 |
| 9 | TT11 | 19 | 0Tn0 |
| 10 | TT00 | 20 | 00nn |

Simplified cases for topology 2H1.

| | | | |
|---|---|---|---|
| 1 | TTgg | 12 | T0g0 |
| 2 | TTg1 | 13 | T01g |
| 3 | TTg0 | 14 | T00g |
| 4 | TT1g | 15 | 0Tng |
| 5 | TT0g | 16 | T011 |
| 6 | T0gg | 17 | T010 |
| 7 | TT11 | 18 | T001 |
| 8 | TT10 | 19 | T000 |
| 9 | TT01 | 20 | 0Tn1 |
| 10 | TT00 | 21 | 0Tn0 |
| 11 | T0g1 | 22 | 00nn |

Simplified cases for topology 2H2.

The detailed analysis enabled us to identify a total of 155 variants of models with fixed permutations. We assigned each model to one of 15 types: 2H1, 2H2, 1H1, 1H2, 1H3, 1H4, 2HP, 1HP1, 1HP2, 1HP3, T1, T2, PT, TP, P. Here, 2H or 1H means the presence of two or one hybridizations, respectively; T or P indicates the presence of a tree or (partial) polytomy. (Note, in fact, types 1H1, 1H2, 1H3 were discussed previously (Doronina et al., 2017)).

Depending on the number of free parameters (dimension of the corresponding subspace), all 155 models can be divided into five classes: $\Theta_0$, $\Theta_1$, $\Theta_2$, $\Theta_3$, $\Theta_4$, and the index value is one less than the number of parameters for models in the corresponding class (see Table 2). Note that the class $\Theta_0$ consists of one element - complete polytomy.

These 15 model types will be considered as subclasses of the corresponding classes and denoted $\Theta_j^i$ (where index $0 \le j \le 4$ is number of the class of the model, and $i$ - is number of the model in the related class, see Table 2).

**Table 2. Set of non-redundant complete of trees.**

| Model | Name | Alias | $v$ | Tree variant | σ = $(j_1, j_2, j_3, j_4)$ | Total |
|---|---|---|---|---|---|---|
| $\Theta_4^1$ | 2H1 | H1:TTgg | $(n_0, T_1, T_3, \gamma_1, \gamma_3)$ | 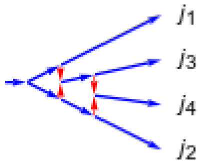  | all permutations | 24 |
| $\Theta_4^2$ | 2H2 | H2:TTgg | $(n_0, T_1, T_3, \gamma_1, \gamma_3)$ | 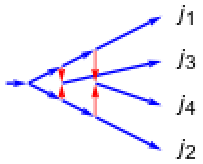  | 1234, 1243, 1324, 1342, 1423, 1432, 2314, 2341, 2413, 2431, 3412, 3421 | 12 |
| $\Theta_3^1$ | 2HP | H2:T0gg | $(n_0, T_1, \gamma_1, \gamma_3)$ | 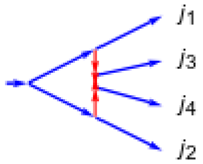  | 1234, 1324, 1423, 2314, 2413, 3412 | 6 |
| $\Theta_3^2$ | 1H1 | H1:TTg0 | $(n_0, T_1, T_3, \gamma_1)$ | 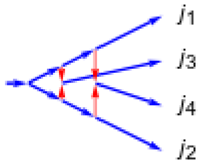  | 1234, 1243, 1324, 2134, 2143, 2314, 3124, 3142, 3214, 4123, 4132, 4213 | 12 |
| $\Theta_3^3$ | 1H2 | H1:TT1g | $(n_0, T_1, T_3, \gamma_3)$ | 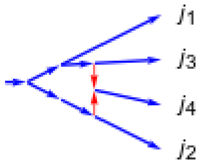  | all permutations | 24 |
| $\Theta_3^4$ | 1H3 | H1:TT0g | $(n_0, T_1, T_3, \gamma_3)$ | 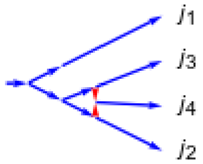  | 1234, 1243, 1342, 2134, 2143, 2341, 3124, 3142, 3241, 4123, 4132, 4231 | 12 |

| | | | | | | |
|---|---|---|---|---|---|---|
| $\Theta_3^5$ | 1H4 | H1:TTg1 | $(n_0, T_1, T_3, \gamma_1)$ | 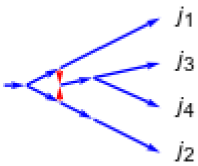 | 1234, 1324, 1423, 2314, 2413, 3412 | 6 |
| $\Theta_2^1$ | 1HP1 | H1:0Tng | $(n_0, T_3, \gamma_3)$ | 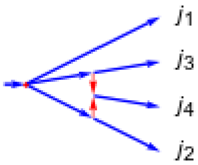 | 1234, 1243, 1342, 2134, 2143, 2341, 3124, 3142, 3241, 4123, 4132, 4231 | 12 |
| $\Theta_2^2$ | 1HP2 | H1:T01g | $(n_0, T_1, \gamma_1)$ | 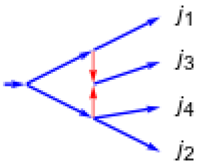 | 1234, 1243, 1324, 2134, 2143, 2314, 3124, 3142, 3214, 4123, 4132, 4213 | 12 |
| $\Theta_2^3$ | 1HP3 | H1:T0g1 | $(n_0, T_1, \gamma_1)$ | 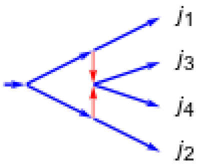 | 1234, 1324, 1423, 2314, 2413, 3412 | 6 |
| $\Theta_2^4$ | T1 | H1:TT10 | $(n_0, T_1, T_3)$ | 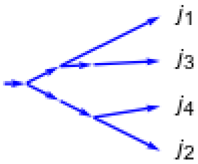 | 1234, 1243, 1324, 2134, 2143, 3142 | 6 |
| $\Theta_2^5$ | T2 | H1:TT01 | $(n_0, T_1, T_3)$ | 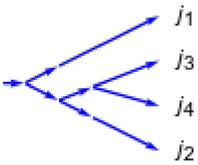 | 1234, 1324, 1423, 2134, 2314, 2413, 3124, 3214, 3412, 4123, 4213, 4312 | 12 |
| $\Theta_1^1$ | PT | H1:0Tn1 | $(n_0, T_3)$ | 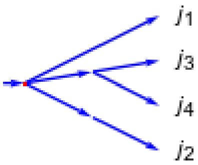 | 1234, 1324, 1423, 2314, 2413, 3412 | 6 |
| $\Theta_1^2$ | TP | H1:T001 | $(n_0, T_1)$ | 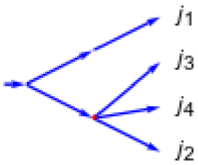 | 1234, 2134, 3124, 4123 | 4 |
| $\Theta_0^1$ | P | H1:00nn | $n_0$ | 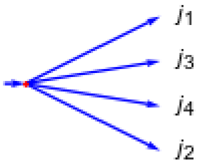 | 1234 | 1 |

#### 5. Statistical testing

With the 15 described phylogenetic models applied on the 155 possible tree topologies we need to find one tree, which represents the most probable phylogenetic relationship. For this tree, we must apply statistical testing of significance.

Therefore, we have a chain of parametric sets:

$$\Theta_j^i \subset \Theta$$

(where every set $\Theta_j^i$ defines a ($j$+1)-dimensional manifold in the nonnegative orthant in 10-dimensional space of parameters $a_{i,j}$).

Maximum likelihood estimates the parameter θ for model $\theta \subset \Theta$. This results in $\hat{\theta}_j = \hat{\theta}_j(x) \in \Theta_j$ (here $x$ is the observed value of $\xi$), where the maximum value of the likelihood function:

$$l\left(x; \hat{\theta}_j^i(x)\right) = \max_{\theta \in \Theta_j^i} l(x; \theta) = L\left(x|\Theta_j^i\right) \quad \text{(S1.5.1)}$$

is achieved (here $\hat{\theta}_j^i$ actually represents one of the models listed in Table 2 with some specific permutation and some values of parameters $(n_0, T_1, T_3, \gamma_1, \gamma_3)$).

For selecting of the optimal model we consider the logarithmic likelihood ratio:

$$\lambda_j^k(x) = 2\log\frac{L(x|\Theta)}{L\left(x|\Theta_j^i\right)}\ (0 \le j \le 3), \quad \text{(S1.5.2)}$$

Note if we have $\Theta = \Theta_4^1 \cup \Theta_4^2$ hence $L(x|\Theta)$ equaled to the largest of the values from: $L(x|\Theta_4^1)$ and $L(x|\Theta_4^2)$.

Corresponding p-value is:

$$p_j^k(x) = \sup_{\theta \in \Theta_j^i} \mathrm{P}_\theta\left(\lambda_j^i(\xi) \ge \lambda_j^i(x)\right). \quad \text{(S1.5.3)}$$

Using, for example, the chi-square approximation, we can then write:

$$p_j^k(x) \approx \mathrm{P}\left(\chi_{4-j}^2 \ge \lambda_j^i(x)\right), \quad \text{(S1.5.4)}$$

where $\chi_s^2$ is a random variable distributed according to the chi-square distribution with $s$ degrees of freedom.

However, due to the limited accuracy of the distribution approximation, at large values of $\lambda_j^i(x)$ using the S1.5.4 approximation for p-value might lead to inappropriate values. Therefore, in practice, it is necessary to check condition: $p_j^i(x) \le \alpha$, where α is the selected significance level.

The last inequality is equivalent to: $\lambda_j^i(x) \ge$ z, where z is defined as (1 – α)·100% quantile of the corresponding statistic distribution $\lambda_j^i(\xi)$. If the real distribution of $\lambda_j^i(\xi)$ is not known, and not fit to the chi-square approximation the quantile of its empirical distribution could be taken as the critical value for statistics after performing computer modeling also known as empirical cumulative distribution function. For this, a sample of sufficient size M sets of 10 random Poisson-distributed variables with parameters $a_{i,j}$ calculated by formula (S1.4.1) is generated, where values of parameters $(n_0, T_1, T_3, \gamma_1, \gamma_3)$ are taken from $\hat{\theta}_j^i$. Next, for each set of values of $L(x|\Theta), L\left(x|\Theta_j^i\right)$, can be derived, and according formula (S1.5.2) $\lambda_j^i$ is calculated. Sorting the obtained values in ascending order, we take z equivalent to (1-α)·M'th value of $\lambda_j^i$.

It should be taken into account that for polytomy maximum likelihood of parameter θ can be calculated analytically. Here, all $a_{i,j} = \frac{n_0}{6}$, whence, according to (S1.4.11), it follows:

$$\log l(x;\theta) = n \cdot log\frac{n_0}{6} - \frac{5n_0}{3}$$

where,

$$n = \sum_{1\le i\le j\le 4} x_{i,j}.$$

Thus $\max_{\theta\in\Theta_0^1} l(x;\theta)$ is achieved when $n_0 = \frac{3n}{5}$, and $\log L(x|\Theta_0^1) = n \cdot \left(\log\frac{n}{10} - 1\right)$.

Due to the high complexity of the expressions for the likelihood function $l(x;\theta)$, its maximization in the cases $\theta \in \Theta_j$ and $j \neq 0$ should be performed computationally.

#### References

Doronina L, Churakov G, Kuritzin A, Shi J, Baertsch R, Clawson H, Schmitz J. (2017) Speciation network in Laurasiatheria: retrophylogenomic signals. Genome Res, 27:997-1003.

Fisher R.A. On the dominance ratio. Proceedings Royal Society Edinburg 1922; 42:321-241.

Kimura M. (1955a) Stochastic processes and distribution of gene frequencies under natural selection. *Cold Spring Harb Symp Quant. Biol* 20: 33–53

Kimura M. (1955b) Solution of a process of random genetic drift with a continuous model. *Proc Natl Acad Sci USA* 41:144-150

Kuritzin A, Kischka T, Schmitz J, Churakov G. (2016) Incomplete Lineage Sorting and Hybridization Statistics for Large-Scale Retroposon Insertion Data. *PLoS Comput Biol.* 12:e1004812.

Tran T., Hofrichter J., Jost J. An introduction to the mathematical structure of the Wright–Fisher model of population genetics. Theory Bioscience 2013; 132:73-82.

Wright S. Evolution in Mendelian populations. Genetics 1931; 16:97-159.

Waxman D. A unified treatment of the probability of fixation when population size and the strength of selection change over time. Genetics 2011; 188:907–913.
