## Supplementary tables (symulations) for "A 4-lineage statistical suite to evaluate the support of large-scale retrotransposon insertion data to reconstruct evolutionary trees"

### Table S1.

**We are comparing the *stepwise* method with chi-square distribution against a 50% chi-square distribution with a 50% point mass 0 mixture criterion for frequency of correct recognition of models in the Poisson randomized sets.** The first column indicates group *Θj*, the following two columns show the trivial name of the model and its alias in TTgg notation, and the following four columns describe the initial values of parameters T1, T3, $\gamma_1$, $\gamma_3$ taken in the Poisson-based randomization process where n0=100. The following eight columns are grouped into four columns according to our selected criterion: *chi-square* indicating chi-square distribution with one degree of freedom; and *50% chi-square plus 50% point mass 0* indicating a mixture consisting from 50% chi-square distribution with one degree of freedom and 50% of point mass 0 distribution. In these groups, we annotated the frequency of the correct model recognition: $\Theta^{i}_{j}$; frequency of error recognition of the model in the direction of a simple model: *Θj-* (second type of error); frequency of error recognition of the model to a complex model: *Θj+* (first type of error); frequency of error recognition of the model because optimization process selects another model from the same group: *Θj*.

| Group | Model | Alias | $T_1$ | $T_3$ | $\gamma_1$ | $\gamma_3$ | *chi-square* | | | | *50% chi-square plus 50% point mass 0* | | | |
|---|---|---|---|---|---|---|---|---|---|---|---|---|---|---|
| | | | | | | | $\Theta^{i}_{j}$ | $\Theta_{j-}$ | $\Theta_{j+}$ | $\Theta_j$ | $\Theta^{i}_{j}$ | $\Theta_{j-}$ | $\Theta_{j+}$ | $\Theta_j$ |
| **$\Theta_0$** | **P** | **H1:00nn** | **0** | **0** | **-** | **-** | 0.76 | 0 | 0.24 | 0 | 0.70 | 0 | 0.30 | 0 |
| **$\Theta_1$** | **TP** | **H1:0Tn1** | **0** | **1** | **-** | **1** | 0.82 | 0 | 0.18 | 0 | 0.81 | 0 | 0.19 | 0 |
| **$\Theta_1$** | **PT** | **H1:T00n** | **1** | **0** | **0** | **-** | 0.85 | 0 | 0.15 | 0 | 0.80 | 0 | 0.19 | 0.01 |
| **$\Theta_2$** | **1HP1** | **H1:0Tng** | **0** | **1** | **-** | **0.5** | 0.92 | 0 | 0.08 | 0 | 0.87 | 0 | 0.13 | 0 |
| **$\Theta_2$** | **1HP2** | **H1:T0g0** | **1** | **0** | **0.5** | **0** | 0.89 | 0 | 0.11 | 0 | 0.85 | 0 | 0.15 | 0 |
| **$\Theta_2$** | **1HP3** | **H1:T0g1** | **1** | **0** | **0.5** | **1** | 0.88 | 0 | 0.12 | 0 | 0.83 | 0 | 0.17 | 0 |
| **$\Theta_2$** | **T1** | **H1:TT10** | **0.5** | **0.5** | **1** | **0** | 0.87 | 0 | 0.12 | 0.01 | 0.82 | 0 | 0.17 | 0.01 |
| **$\Theta_2$** | **T2** | **H1:TT01** | **0.5** | **0.5** | **0** | **1** | 0.87 | 0 | 0.13 | 0 | 0.85 | 0 | 0.15 | 0 |
| **$\Theta_3$** | **1H1** | **H1:TTg0** | **0.5** | **0.5** | **0.5** | **0** | 0.86 | 0.04 | 0.10 | 0 | 0.85 | 0.02 | 0.12 | 0.01 |
| **$\Theta_3$** | **1H2** | **H1:TT1g** | **0.5** | **0.5** | **1** | **0.5** | 0.95 | 0 | 0.05 | 0 | 0.93 | 0.01 | 0.06 | 0 |
| **$\Theta_3$** | **1H3** | **H1:TT0g** | **0.5** | **0.5** | **0** | **0.5** | 0.89 | 0.04 | 0.04 | 0.03 | 0.90 | 0.02 | 0.06 | 0.02 |
| **$\Theta_3$** | **1H4** | **H1:TTg1** | **0.5** | **0.5** | **0.5** | **1** | 0.90 | 0.01 | 0.08 | 0.01 | 0.91 | 0 | 0.08 | 0.01 |
| **$\Theta_3$** | **2PH** | **H2:T0gg** | **1** | **0** | **0.25** | **0.5** | 0.93 | 0 | 0.07 | 0 | 0.91 | 0.01 | 0.08 | 0 |
| **$\Theta_4$** | **2H1** | **H1:TTgg** | **0.5** | **0.5** | **0.5** | **0.5** | 0.92 | 0.08 | 0 | 0 | 0.93 | 0.07 | 0 | 0 |
| **$\Theta_4$** | **2H2** | **H2:TTgg** | **0.5** | **0.5** | **0.5** | **0.5** | 0.94 | 0.06 | 0 | 0 | 0.94 | 0.06 | 0 | 0 |

### Table S2.

**Comparing the *reverse* method and chi-square distribution against binary chi-square distribution according to degrees of freedom plus point mass 0 mixture and binary chi-square distribution according to degrees of freedom plus point mass 1 mixture criteria for frequency of correct recognition of models in the Poisson randomized set.** The first column indicates group *Θj*, the following two columns show the trivial name of the model and its alias in TTgg notation, and the following four columns describe initial values of parameters T1, T3, $\gamma_1$, $\gamma_3$ taken in Poisson based randomization process, where n0=100. Next, twelve columns are grouped into four columns according to the selected criterion: *chi-square* indicating chi-square distribution with different degrees of freedom; *chi-squares mixture with 0 p.m.* annotating mixture of several chi-square distributions with different degrees of freedom and binary coefficients plus point mass 0; *chi-squares mixture with 1 p.m.* indicating mixture of several chi-square distributions with different degrees of freedom and binary coefficients plus point mass 1. In these groups, we annotated the frequency of the correct model recognition: $\Theta_j^i$; frequency of error recognition of the model in the direction of a simple model: *Θj-* (second type of error); frequency of error recognition of the model to a complex model: *Θj+* (first type of error); frequency of error recognition of the model because optimization process selects another model from the same group: *Θj*.

| Group | Model | Alias | $T_1$ | $T_3$ | $\gamma_1$ | $\gamma_3$ | ***chi-square*** | | | | ***chi-squares mixture with 0 p.m.*** | | | | ***chi-squares mixture with 1 p.m.*** | | | |
|---|---|---|---|---|---|---|---|---|---|---|---|---|---|---|---|---|---|---|
| | | | | | | | $\Theta_j^i$ | $\Theta_{j-}$ | $\Theta_{j+}$ | $\Theta_j$ | $\Theta_j^i$ | $\Theta_{j-}$ | $\Theta_{j+}$ | $\Theta_j$ | $\Theta_j^i$ | $\Theta_{j-}$ | $\Theta_{j+}$ | $\Theta_j$ |
| $\Theta_0$ | P | H1:00nn | 0 | 0 | - | - | 0.98 | 0 | 0.02 | 0 | 0.89 | 0 | 0.11 | 0 | 0.90 | 0 | 0.10 | 0 |
| $\Theta_1$ | TP | H1:0Tn1 | 0 | 1 | - | 1 | 0.91 | 0 | 0.09 | 0 | 0.87 | 0 | 0.13 | 0 | 0.88 | 0 | 0.12 | 0 |
| $\Theta_1$ | PT | H1:T00n | 1 | 0 | 0 | - | 0.93 | 0 | 0.07 | 0 | 0.88 | 0 | 0.12 | 0 | 0.89 | 0 | 0.11 | 0 |
| $\Theta_2$ | 1HP1 | H1:0Tng | 0 | 0.5 | - | 0.5 | 0.91 | 0.01 | 0.07 | 0.01 | 0.85 | 0 | 0.14 | 0.01 | 0.87 | 0 | 0.12 | 0.01 |
| $\Theta_2$ | 1HP2 | H1:T0g0 | 1 | 0 | 0.5 | 0 | 0.93 | 0 | 0.07 | 0 | 0.88 | 0 | 0.12 | 0 | 0.88 | 0 | 0.12 | 0 |
| $\Theta_2$ | 1HP3 | H1:T0g1 | 1 | 0 | 0.5 | 1 | 0.93 | 0 | 0.07 | 0 | 0.84 | 0 | 0.15 | 0.01 | 0.86 | 0 | 0.13 | 0.01 |
| $\Theta_2$ | T1 | H1:TT10 | 0.5 | 0.5 | 1 | 0 | 0.92 | 0 | 0.07 | 0.01 | 0.87 | 0 | 0.12 | 0.01 | 0.87 | 0 | 0.13 | 0 |
| $\Theta_2$ | T2 | H1:TT01 | 0.5 | 0.5 | 0 | 1 | 0.94 | 0 | 0.06 | 0 | 0.86 | 0 | 0.14 | 0 | 0.88 | 0 | 0.12 | 0 |
| $\Theta_3$ | 1H1 | H1:TTg0 | 0.5 | 0.5 | 0.5 | 0 | 0.83 | 0.07 | 0.10 | 0 | 0.82 | 0.05 | 0.13 | 0 | 0.81 | 0.04 | 0.15 | 0 |
| $\Theta_3$ | 1H2 | H1:TT1g | 0.5 | 0.5 | 1 | 0.5 | 0.95 | 0 | 0.05 | 0 | 0.92 | 0 | 0.07 | 0.01 | 0,94 | 0 | 0.05 | 0.01 |
| $\Theta_3$ | 1H3 | H1:TT0g | 0.5 | 0.5 | 0 | 0.5 | 0.86 | 0.08 | 0.04 | 0.02 | 0.87 | 0.06 | 0.06 | 0.01 | 0.86 | 0.03 | 0.11 | 0 |
| $\Theta_3$ | 1H4 | H1:TTg1 | 0.5 | 0.5 | 0.5 | 1 | 0.89 | 0.03 | 0.07 | 0.01 | 0.88 | 0.02 | 0.09 | 0.01 | 0.88 | 0.01 | 0.10 | 0.01 |
| $\Theta_3$ | 2PH | H2:T0gg | 1 | 0 | 0.25 | 0.75 | 0.89 | 0.01 | 0.10 | 0 | 0.86 | 0 | 0.14 | 0 | 0.87 | 0.01 | 0.12 | 0 |
| $\Theta_4$ | 2H1 | H1:TTgg | 0.5 | 0.5 | 0.5 | 0.5 | 0.91 | 0.09 | 0 | 0 | 0.92 | 0.08 | 0 | 0 | 0.93 | 0.07 | 0 | 0 |
| $\Theta_4$ | 2H2 | H2:TTgg | 0.5 | 0.5 | 0.5 | 0.5 | 0.94 | 0.06 | 0 | 0 | 0.95 | 0.05 | 0 | 0 | 0.93 | 0.07 | 0 | 0 |

### Table S3.

**Testing the *reverse* method and chi-square distribution criterion for frequency of correct recognition of models in the Poisson randomized set.** The first column indicates group $\Theta j$, the following two columns show the trivial name of the model and its alias in TTgg notation, and the following four columns describe the initial values of parameters T1, T3, $\gamma_1$, $\gamma_3$ taken in the Poisson-distribution based randomization process. The last four columns present the frequency of the correct model recognition: $\theta_j^i$; frequency of error recognition of the model in the direction of a simple model:$\Theta j$-; frequency of error recognition of the model to a complex model: $\Theta j$+; frequency of error recognition of the model because optimization process selects another model from the same group: $\Theta j$.

| Group | Model | Alias | $T_1$ | $T_3$ | $\gamma_1$ | $\gamma_3$ | $\theta_j^i$ | $\Theta_{j-}$ | $\Theta_{j+}$ | $\Theta_j$ |
|---|---|---|---|---|---|---|---|---|---|---|
| $\Theta_0$ | P | H1:00nn | 0 | 0 | - | - | 0.93 | 0 | 0.07 | 0 |
| $\Theta_1$ | TP | H1:0Tn1 | 0 | 0.5 | - | 1 | 0.94 | 0 | 0.06 | 0 |
| $\Theta_1$ | TP | H1:0Tn1 | 0 | 1 | - | 1 | 0.91 | 0 | 0.09 | 0 |
| $\Theta_1$ | TP | H1:0Tn1 | 0 | 1.5 | - | 1 | 0.93 | 0 | 0.07 | 0 |
| $\Theta_1$ | PT | H1:T00n | 0.5 | 0 | 0 | - | 0.95 | 0 | 0.05 | 0 |
| $\Theta_1$ | PT | H1:T00n | 1 | 0 | 0 | - | 0.93 | 0 | 0.07 | 0 |
| $\Theta_1$ | PT | H1:T00n | 1.5 | 0 | 0 | - | 0.95 | 0 | 0.05 | 0 |
| $\Theta_2$ | 1HP1 | H1:0Tng | 0 | 0.5 | - | 0.25 | 0.65 | 0.27 | 0.05 | 0.03 |
| $\Theta_2$ | 1HP1 | H1:0Tng | 0 | 0.5 | - | 0.5 | 0.91 | 0.01 | 0.07 | 0.01 |
| $\Theta_2$ | 1HP1 | H1:0Tng | 0 | 1 | - | 0.25 | 0.96 | 0 | 0.04 | 0 |
| $\Theta_2$ | 1HP1 | H1:0Tng | 0 | 1 | - | 0.5 | 0.96 | 0 | 0.04 | 0 |
| $\Theta_2$ | 1HP1 | H1:0Tng | 0 | 1.5 | - | 0.25 | 0.96 | 0 | 0.04 | 0 |
| $\Theta_2$ | 1HP1 | H1:0Tng | 0 | 1.5 | - | 0.5 | 0.95 | 0 | 0.05 | 0 |
| $\Theta_2$ | 1HP2 | H1:T0g0 | 0.5 | 0 | 0.25 | 0 | 0.83 | 0.02 | 0.05 | 0.1 |
| $\Theta_2$ | 1HP2 | H1:T0g0 | 0.5 | 0 | 0.5 | 0 | 0.90 | 0 | 0.09 | 0.01 |
| $\Theta_2$ | 1HP2 | H1:T0g0 | 0.5 | 0 | 0.75 | 0 | 0.88 | 0 | 0.06 | 0.06 |
| $\Theta_2$ | 1HP2 | H1:T0g0 | 1 | 0 | 0.25 | 0 | 0.92 | 0 | 0.08 | 0 |
| $\Theta_2$ | 1HP2 | H1:T0g0 | 1 | 0 | 0.5 | 0 | 0.93 | 0 | 0.07 | 0 |
| $\Theta_2$ | 1HP2 | H1:T0g0 | 1 | 0 | 0.75 | 0 | 0.92 | 0 | 0.08 | 0 |
| $\Theta_2$ | 1HP2 | H1:T0g0 | 1.5 | 0 | 0.25 | 0 | 0.92 | 0 | 0.08 | 0 |
| $\Theta_2$ | 1HP2 | H1:T0g0 | 1.5 | 0 | 0.5 | 0 | 0.94 | 0 | 0.06 | 0 |
| $\Theta_2$ | 1HP2 | H1:T0g0 | 1.5 | 0 | 0.75 | 0 | 0.93 | 0 | 0.07 | 0 |
| $\Theta_2$ | 1HP3 | H1:T0g1 | 0.5 | 0 | 0.5 | 1 | 0.81 | 0.08 | 0.06 | 0.05 |
| $\Theta_2$ | 1HP3 | H1:T0g1 | 0.5 | 0 | 0.25 | 1 | 0.92 | 0.01 | 0.06 | 0.01 |
| $\Theta_2$ | 1HP3 | H1:T0g1 | 1 | 0 | 0.5 | 1 | 0.93 | 0 | 0.07 | 0 |
| $\Theta_2$ | 1HP3 | H1:T0g1 | 1 | 0 | 0.25 | 1 | 0.93 | 0 | 0.07 | 0 |
| $\Theta_2$ | 1HP3 | H1:T0g1 | 1.5 | 0 | 0.5 | 1 | 0.94 | 0 | 0.06 | 0 |
| $\Theta_2$ | 1HP3 | H1:T0g1 | 1.5 | 0 | 0.25 | 1 | 0.93 | 0 | 0.07 | 0 |
| $\Theta_2$ | T1 | H1:TT10 | 0.125 | 0.375 | 1 | 0 | 0.45 | 0.45 | 0.05 | 0.05 |
| $\Theta_2$ | T1 | H1:TT10 | 0.25 | 0.25 | 1 | 0 | 0.76 | 0.02 | 0.06 | 0.16 |
| $\Theta_2$ | T1 | H1:TT10 | 0.25 | 0.75 | 1 | 0 | 0.87 | 0.03 | 0.10 | 0 |
| $\Theta_2$ | T1 | H1:TT10 | 0.375 | 0.125 | 1 | 0 | 0.51 | 0 | 0.03 | 0.46 |
| $\Theta_2$ | T1 | H1:TT10 | 0.375 | 1.125 | 1 | 0 | 0.93 | 0 | 0.07 | 0 |
| $\Theta_2$ | T1 | H1:TT10 | 0.5 | 0.5 | 1 | 0 | 0.92 | 0 | 0.07 | 0.01 |
| $\Theta_2$ | T1 | H1:TT10 | 0.75 | 0.25 | 1 | 0 | 0.67 | 0 | 0.05 | 0.28 |
| $\Theta_2$ | T1 | H1:TT10 | 0.75 | 0.75 | 1 | 0 | 0.92 | 0 | 0.08 | 0 |
| $\Theta_2$ | T1 | H1:TT10 | 1.125 | 0.375 | 1 | 0 | 0.77 | 0 | 0.04 | 0.19 |
| $\Theta_2$ | T2 | H1:TT01 | 0.125 | 0.375 | 0 | 1 | 0.43 | 0.46 | 0.04 | 0.07 |
| $\Theta_2$ | T2 | H1:TT01 | 0.25 | 0.25 | 0 | 1 | 0.87 | 0.02 | 0.05 | 0.06 |

| | | | | | | | | | | |
|---|---|---|---|---|---|---|---|---|---|---|
| **$\Theta_2$** | **T2** | **H1:TT01** | **0.25** | **0.75** | **0** | **1** | 0.92 | 0.03 | 0.05 | 0 |
| **$\Theta_2$** | **T2** | **H1:TT01** | **0.375** | **0.125** | **0** | **1** | 0.39 | 0.45 | 0.03 | 0.13 |
| **$\Theta_2$** | **T2** | **H1:TT01** | **0.375** | **1.125** | **0** | **1** | 0.95 | 0 | 0.05 | 0 |
| **$\Theta_2$** | **T2** | **H1:TT01** | **0.5** | **0.5** | **0** | **1** | 0.94 | 0 | 0.06 | 0 |
| **$\Theta_2$** | **T2** | **H1:TT01** | **0.75** | **0.25** | **0** | **1** | 0.89 | 0.02 | 0.06 | 0.03 |
| **$\Theta_2$** | **T2** | **H1:TT01** | **0.75** | **0.75** | **0** | **1** | 0.94 | 0 | 0.06 | 0 |
| **$\Theta_3$** | **T2** | **H1:TT01** | **1.125** | **0.375** | **0** | **0** | 0.94 | 0 | 0.06 | 0 |
| **$\Theta_3$** | **1H1** | **H1:TTg0** | **0.125** | **0.375** | **0.25** | **0** | 0.02 | 0.96 | 0.01 | 0.01 |
| **$\Theta_3$** | **1H1** | **H1:TTg0** | **0.125** | **0.375** | **0.5** | **0** | 0.03 | 0.95 | 0.02 | 0 |
| **$\Theta_3$** | **1H1** | **H1:TTg0** | **0.125** | **0.375** | **0.75** | **0** | 0.01 | 0.96 | 0.01 | 0.02 |
| **$\Theta_3$** | **1H1** | **H1:TTg0** | **0.25** | **0.25** | **0.25** | **0** | 0.05 | 0.90 | 0.03 | 0.02 |
| **$\Theta_3$** | **1H1** | **H1:TTg0** | **0.25** | **0.25** | **0.5** | **0** | 0.11 | 0.82 | 0.04 | 0.03 |
| **$\Theta_3$** | **1H1** | **H1:TTg0** | **0.25** | **0.25** | **0.75** | **0** | 0.04 | 0.90 | 0.02 | 0.04 |
| **$\Theta_3$** | **1H1** | **H1:TTg0** | **0.25** | **0.75** | **0.25** | **0** | 0.12 | 0.83 | 0.03 | 0.02 |
| **$\Theta_3$** | **1H1** | **H1:TTg0** | **0.25** | **0.75** | **0.5** | **0** | 0.25 | 0.70 | 0.03 | 0.02 |
| **$\Theta_3$** | **1H1** | **H1:TTg0** | **0.25** | **0.75** | **0.75** | **0** | 0.11 | 0.84 | 0.01 | 0.04 |
| **$\Theta_3$** | **1H1** | **H1:TTg0** | **0.375** | **0.125** | **0.25** | **0** | 0.02 | 0.92 | 0.03 | 0.03 |
| **$\Theta_3$** | **1H1** | **H1:TTg0** | **0.375** | **0.125** | **0.5** | **0** | 0.07 | 0.85 | 0.05 | 0.03 |
| **$\Theta_3$** | **1H1** | **H1:TTg0** | **0.375** | **0.125** | **0.75** | **0** | 0.02 | 0.91 | 0.02 | 0.05 |
| **$\Theta_3$** | **1H1** | **H1:TTg0** | **0.375** | **1.125** | **0.25** | **0** | 0.24 | 0.70 | 0.03 | 0.03 |
| **$\Theta_3$** | **1H1** | **H1:TTg0** | **0.375** | **1.125** | **0.5** | **0** | 0.53 | 0.40 | 0.05 | 0.02 |
| **$\Theta_3$** | **1H1** | **H1:TTg0** | **0.375** | **1.125** | **0.75** | **0** | 0.20 | 0.72 | 0.04 | 0.04 |
| **$\Theta_3$** | **1H1** | **H1:TTg0** | **0.5** | **0.5** | **0.25** | **0** | 0.50 | 0.42 | 0.07 | 0.01 |
| **$\Theta_3$** | **1H1** | **H1:TTg0** | **0.5** | **0.5** | **0.5** | **0** | 0.83 | 0.07 | 0.10 | 0 |
| **$\Theta_3$** | **1H1** | **H1:TTg0** | **0.5** | **0.5** | **0.75** | **0** | 0.45 | 0.44 | 0.07 | 0.04 |
| **$\Theta_3$** | **1H1** | **H1:TTg0** | **0.75** | **0.25** | **0.25** | **0** | 0.55 | 0.36 | 0.08 | 0.01 |
| **$\Theta_3$** | **1H1** | **H1:TTg0** | **0.75** | **0.25** | **0.5** | **0** | 0.47 | 0.45 | 0.08 | 0 |
| **$\Theta_3$** | **1H1** | **H1:TTg0** | **0.75** | **0.25** | **0.75** | **0** | 0.34 | 0.58 | 0.06 | 0.02 |
| **$\Theta_3$** | **1H1** | **H1:TTg0** | **0.75** | **0.75** | **0.25** | **0** | 0.80 | 0.09 | 0.10 | 0.01 |
| **$\Theta_3$** | **1H1** | **H1:TTg0** | **0.75** | **0.75** | **0.5** | **0** | 0.91 | 0 | 0.09 | 0 |
| **$\Theta_3$** | **1H1** | **H1:TTg0** | **0.75** | **0.75** | **0.75** | **0** | 0.75 | 0.17 | 0.07 | 0.01 |
| **$\Theta_3$** | **1H1** | **H1:TTg0** | **1.125** | **0.375** | **0.25** | **0** | 0.81 | 0.09 | 0.10 | 0 |
| **$\Theta_3$** | **1H1** | **H1:TTg0** | **1.125** | **0.375** | **0.5** | **0** | 0.69 | 0.23 | 0.07 | 0.01 |
| **$\Theta_3$** | **1H1** | **H1:TTg0** | **1.125** | **0.375** | **0.75** | **0** | 0.55 | 0.37 | 0.07 | 0.01 |
| **$\Theta_3$** | **1H2** | **H1:TT1g** | **0.125** | **0.375** | **1** | **0.25** | 0.31 | 0.66 | 0.02 | 0.01 |
| **$\Theta_3$** | **1H2** | **H1:TT1g** | **0.125** | **0.375** | **1** | **0.5** | 0.47 | 0.49 | 0.04 | 0 |
| **$\Theta_3$** | **1H2** | **H1:TT1g** | **0.125** | **0.375** | **1** | **0.75** | 0.34 | 0.61 | 0.03 | 0.02 |
| **$\Theta_3$** | **1H2** | **H1:TT1g** | **0.25** | **0.25** | **1** | **0.25** | 0.40 | 0.55 | 0.02 | 0.03 |
| **$\Theta_3$** | **1H2** | **H1:TT1g** | **0.25** | **0.25** | **1** | **0.5** | 0.73 | 0.2 | 0.03 | 0.04 |
| **$\Theta_3$** | **1H2** | **H1:TT1g** | **0.25** | **0.25** | **1** | **0.75** | 0.55 | 0.41 | 0.01 | 0.03 |
| **$\Theta_3$** | **1H2** | **H1:TT1g** | **0.25** | **0.75** | **1** | **0.25** | 0.93 | 0.04 | 0.03 | 0 |
| **$\Theta_3$** | **1H2** | **H1:TT1g** | **0.25** | **0.75** | **1** | **0.5** | 0.91 | 0.04 | 0.05 | 0 |
| **$\Theta_3$** | **1H2** | **H1:TT1g** | **0.25** | **0.75** | **1** | **0.75** | 0.91 | 0.04 | 0.05 | 0 |
| **$\Theta_3$** | **1H2** | **H1:TT1g** | **0.375** | **0.125** | **1** | **0.25** | 0.16 | 0.81 | 0 | 0.03 |
| **$\Theta_3$** | **1H2** | **H1:TT1g** | **0.375** | **0.125** | **1** | **0.5** | 0.21 | 0.73 | 0.02 | 0.04 |

| | | | | | | | | | | |
|---|---|---|---|---|---|---|---|---|---|---|
| **$\Theta_3$** | **1H2** | **H1:TT1g** | **0.375** | **0.125** | **1** | **0.75** | 0.24 | 0.72 | 0.01 | 0.03 |
| **$\Theta_3$** | **1H2** | **H1:TT1g** | **0.375** | **1.125** | **1** | **0.25** | 0.95 | 0 | 0.05 | 0 |
| **$\Theta_3$** | **1H2** | **H1:TT1g** | **0.375** | **1.125** | **1** | **0.5** | 0.95 | 0 | 0.05 | 0 |
| **$\Theta_3$** | **1H2** | **H1:TT1g** | **0.375** | **1.125** | **1** | **0.75** | 0.96 | 0 | 0.04 | 0 |
| **$\Theta_3$** | **1H2** | **H1:TT1g** | **0.5** | **0.5** | **1** | **0.25** | 0.94 | 0.01 | 0.05 | 0 |
| **$\Theta_3$** | **1H2** | **H1:TT1g** | **0.5** | **0.5** | **1** | **0.5** | 0.95 | 0 | 0.05 | 0 |
| **$\Theta_3$** | **1H2** | **H1:TT1g** | **0.5** | **0.5** | **1** | **0.75** | 0.95 | 0 | 0.05 | 0 |
| **$\Theta_3$** | **1H2** | **H1:TT1g** | **0.75** | **0.25** | **1** | **0.25** | 0.51 | 0.42 | 0.02 | 0.05 |
| **$\Theta_3$** | **1H2** | **H1:TT1g** | **0.75** | **0.25** | **1** | **0.5** | 0.68 | 0.26 | 0.03 | 0.03 |
| **$\Theta_3$** | **1H2** | **H1:TT1g** | **0.75** | **0.25** | **1** | **0.75** | 0.80 | 0.11 | 0.05 | 0.04 |
| **$\Theta_3$** | **1H2** | **H1:TT1g** | **0.75** | **0.75** | **1** | **0.25** | 0.96 | 0 | 0.04 | 0 |
| **$\Theta_3$** | **1H2** | **H1:TT1g** | **0.75** | **0.75** | **1** | **0.5** | 0.94 | 0 | 0.06 | 0 |
| **$\Theta_3$** | **1H2** | **H1:TT1g** | **0.75** | **0.75** | **1** | **0.75** | 0.95 | 0 | 0.05 | 0 |
| **$\Theta_3$** | **1H2** | **H1:TT1g** | **1.125** | **0.375** | **1** | **0.25** | 0.71 | 0.24 | 0.03 | 0.02 |
| **$\Theta_3$** | **1H2** | **H1:TT1g** | **1.125** | **0.375** | **1** | **0.5** | 0.87 | 0.06 | 0.06 | 0.01 |
| **$\Theta_3$** | **1H2** | **H1:TT1g** | **1.125** | **0.375** | **1** | **0.75** | 0.93 | 0.01 | 0.05 | 0.01 |
| **$\Theta_3$** | **1H3** | **H1:TT0g** | **0.125** | **0.375** | **0** | **0.25** | 0.09 | 0.85 | 0.02 | 0.04 |
| **$\Theta_3$** | **1H3** | **H1:TT0g** | **0.125** | **0.375** | **0** | **0.5** | 0.26 | 0.68 | 0.02 | 0.04 |
| **$\Theta_3$** | **1H3** | **H1:TT0g** | **0.25** | **0.25** | **0** | **0.25** | 0.08 | 0.85 | 0.02 | 0.05 |
| **$\Theta_3$** | **1H3** | **H1:TT0g** | **0.25** | **0.25** | **0** | **0.5** | 0.18 | 0.76 | 0.01 | 0.05 |
| **$\Theta_3$** | **1H3** | **H1:TT0g** | **0.25** | **0.75** | **0** | **0.25** | 0.85 | 0.09 | 0.04 | 0.02 |
| **$\Theta_3$** | **1H3** | **H1:TT0g** | **0.25** | **0.75** | **0** | **0.5** | 0.93 | 0.02 | 0.05 | 0 |
| **$\Theta_3$** | **1H3** | **H1:TT0g** | **0.375** | **0.125** | **0** | **0.25** | 0.02 | 0.96 | 0 | 0.02 |
| **$\Theta_3$** | **1H3** | **H1:TT0g** | **0.375** | **0.125** | **0** | **0.5** | 0.02 | 0.96 | 0 | 0.02 |
| **$\Theta_3$** | **1H3** | **H1:TT0g** | **0.375** | **1.125** | **0** | **0.25** | 0.94 | 0 | 0.06 | 0 |
| **$\Theta_3$** | **1H3** | **H1:TT0g** | **0.375** | **1.125** | **0** | **0.5** | 0.94 | 0 | 0.06 | 0 |
| **$\Theta_3$** | **1H3** | **H1:TT0g** | **0.5** | **0.5** | **0** | **0.25** | 0.49 | 0.42 | 0.04 | 0.05 |
| **$\Theta_3$** | **1H3** | **H1:TT0g** | **0.5** | **0.5** | **0** | **0.5** | 0.86 | 0.08 | 0.04 | 0.02 |
| **$\Theta_3$** | **1H3** | **H1:TT0g** | **0.75** | **0.25** | **0** | **0.25** | 0.10 | 0.85 | 0.01 | 0.04 |
| **$\Theta_3$** | **1H3** | **H1:TT0g** | **0.75** | **0.25** | **0** | **0.5** | 0.18 | 0.77 | 0.02 | 0.03 |
| **$\Theta_3$** | **1H3** | **H1:TT0g** | **0.75** | **0.75** | **0** | **0.25** | 0.81 | 0.11 | 0.05 | 0.03 |
| **$\Theta_3$** | **1H3** | **H1:TT0g** | **0.75** | **0.75** | **0** | **0.5** | 0.95 | 0 | 0.05 | 0 |
| **$\Theta_3$** | **1H3** | **H1:TT0g** | **1.125** | **0.375** | **0** | **0.25** | 0.19 | 0.74 | 0.03 | 0.04 |
| **$\Theta_3$** | **1H3** | **H1:TT0g** | **1.125** | **0.375** | **0** | **0.5** | 0.47 | 0.46 | 0.03 | 0.04 |
| **$\Theta_3$** | **1H4** | **H1:TTg1** | **0.125** | **0.375** | **0.25** | **1** | 0.04 | 0.95 | 0 | 0.01 |
| **$\Theta_3$** | **1H4** | **H1:TTg1** | **0.125** | **0.375** | **0.5** | **1** | 0.04 | 0.95 | 0 | 0.01 |
| **$\Theta_3$** | **1H4** | **H1:TTg1** | **0.25** | **0.25** | **0.25** | **1** | 0.13 | 0.82 | 0.01 | 0.04 |
| **$\Theta_3$** | **1H4** | **H1:TTg1** | **0.25** | **0.25** | **0.5** | **1** | 0.32 | 0.63 | 0.01 | 0.04 |
| **$\Theta_3$** | **1H4** | **H1:TTg1** | **0.25** | **0.75** | **0.25** | **1** | 0.11 | 0.84 | 0 | 0.05 |
| **$\Theta_3$** | **1H4** | **H1:TTg1** | **0.25** | **0.75** | **0.5** | **1** | 0.23 | 0.72 | 0.02 | 0.03 |
| **$\Theta_3$** | **1H4** | **H1:TTg1** | **0.375** | **0.125** | **0.25** | **1** | 0.25 | 0.70 | 0.01 | 0.04 |
| **$\Theta_3$** | **1H4** | **H1:TTg1** | **0.375** | **0.125** | **0.5** | **1** | 0.57 | 0.38 | 0.04 | 0.01 |
| **$\Theta_3$** | **1H4** | **H1:TTg1** | **0.375** | **1.125** | **0.25** | **1** | 0.21 | 0.71 | 0.01 | 0.07 |
| **$\Theta_3$** | **1H4** | **H1:TTg1** | **0.375** | **1.125** | **0.5** | **1** | 0.47 | 0.47 | 0.03 | 0.03 |
| **$\Theta_3$** | **1H4** | **H1:TTg1** | **0.5** | **0.5** | **0.25** | **1** | 0.55 | 0.35 | 0.03 | 0.07 |
| **$\Theta_3$** | **1H4** | **H1:TTg1** | **0.5** | **0.5** | **0.5** | **1** | 0.89 | 0.03 | 0.07 | 0.01 |

| | | | | | | | | | | |
|---|---|---|---|---|---|---|---|---|---|---|
| **$\Theta_3$** | **1H4** | **H1:TTg1** | **0.75** | **0.25** | **0.25** | **1** | 0.87 | 0.05 | 0.06 | 0.02 |
| **$\Theta_3$** | **1H4** | **H1:TTg1** | **0.75** | **0.25** | **0.5** | **1** | 0.93 | 0 | 0.07 | 0 |
| **$\Theta_3$** | **1H4** | **H1:TTg1** | **0.75** | **0.75** | **0.25** | **1** | 0.83 | 0.09 | 0.06 | 0.02 |
| **$\Theta_3$** | **1H4** | **H1:TTg1** | **0.75** | **0.75** | **0.5** | **1** | 0.92 | 0 | 0.08 | 0 |
| **$\Theta_3$** | **1H4** | **H1:TTg1** | **1.125** | **0.375** | **0.25** | **1** | 0.93 | 0 | 0.07 | 0 |
| **$\Theta_3$** | **1H4** | **H1:TTg1** | **1.125** | **0.375** | **0.5** | **1** | 0.91 | 0 | 0.09 | 0 |
| **$\Theta_3$** | **2PH** | **H2:T0gg** | **0.5** | **0** | **0.25** | **0.5** | 0.51 | 0.40 | 0.03 | 0.06 |
| **$\Theta_3$** | **2PH** | **H2:T0gg** | **0.5** | **0** | **0.25** | **0.75** | 0.39 | 0.50 | 0.03 | 0.08 |
| **$\Theta_3$** | **2PH** | **H2:T0gg** | **1** | **0** | **0.25** | **0.5** | 0.92 | 0 | 0.08 | 0 |
| **$\Theta_3$** | **2PH** | **H2:T0gg** | **1** | **0** | **0.25** | **0.75** | 0.89 | 0.01 | 0.10 | 0 |
| **$\Theta_3$** | **2PH** | **H2:T0gg** | **1.5** | **0** | **0.25** | **0.5** | 0.92 | 0 | 0.08 | 0 |
| **$\Theta_3$** | **2PH** | **H2:T0gg** | **1.5** | **0** | **0.25** | **0.75** | 0.92 | 0 | 0.08 | 0 |
| **$\Theta_4$** | **2H1** | **H1:TTgg** | **0.125** | **0.375** | **0.25** | **0.25** | 0.02 | 0.97 | 0 | 0.01 |
| **$\Theta_4$** | **2H1** | **H1:TTgg** | **0.125** | **0.375** | **0.25** | **0.5** | 0.04 | 0.96 | 0 | 0 |
| **$\Theta_4$** | **2H1** | **H1:TTgg** | **0.125** | **0.375** | **0.25** | **0.75** | 0.03 | 0.97 | 0 | 0 |
| **$\Theta_4$** | **2H1** | **H1:TTgg** | **0.125** | **0.375** | **0.5** | **0.25** | 0.02 | 0.97 | 0 | 0.01 |
| **$\Theta_4$** | **2H1** | **H1:TTgg** | **0.125** | **0.375** | **0.5** | **0.5** | 0.04 | 0.96 | 0 | 0 |
| **$\Theta_4$** | **2H1** | **H1:TTgg** | **0.125** | **0.375** | **0.5** | **0.75** | 0.04 | 0.96 | 0 | 0 |
| **$\Theta_4$** | **2H1** | **H1:TTgg** | **0.125** | **0.375** | **0.75** | **0.25** | 0.03 | 0.96 | 0 | 0.01 |
| **$\Theta_4$** | **2H1** | **H1:TTgg** | **0.125** | **0.375** | **0.75** | **0.5** | 0.03 | 0.97 | 0 | 0 |
| **$\Theta_4$** | **2H1** | **H1:TTgg** | **0.125** | **0.375** | **0.75** | **0.75** | 0.02 | 0.97 | 0 | 0.01 |
| **$\Theta_4$** | **2H1** | **H1:TTgg** | **0.25** | **0.25** | **0.25** | **0.25** | 0.03 | 0.96 | 0 | 0.01 |
| **$\Theta_4$** | **2H1** | **H1:TTgg** | **0.25** | **0.25** | **0.25** | **0.5** | 0.05 | 0.95 | 0 | 0 |
| **$\Theta_4$** | **2H1** | **H1:TTgg** | **0.25** | **0.25** | **0.25** | **0.75** | 0.05 | 0.95 | 0 | 0 |
| **$\Theta_4$** | **2H1** | **H1:TTgg** | **0.25** | **0.25** | **0.5** | **0.25** | 0.06 | 0.93 | 0 | 0.01 |
| **$\Theta_4$** | **2H1** | **H1:TTgg** | **0.25** | **0.25** | **0.5** | **0.5** | 0.18 | 0.82 | 0 | 0 |
| **$\Theta_4$** | **2H1** | **H1:TTgg** | **0.25** | **0.25** | **0.5** | **0.75** | 0.14 | 0.86 | 0 | 0 |
| **$\Theta_4$** | **2H1** | **H1:TTgg** | **0.25** | **0.25** | **0.75** | **0.25** | 0.03 | 0.96 | 0 | 0.01 |
| **$\Theta_4$** | **2H1** | **H1:TTgg** | **0.25** | **0.25** | **0.75** | **0.5** | 0.13 | 0.87 | 0 | 0 |
| **$\Theta_4$** | **2H1** | **H1:TTgg** | **0.25** | **0.25** | **0.75** | **0.75** | 0.13 | 0.87 | 0 | 0 |
| **$\Theta_4$** | **2H1** | **H1:TTgg** | **0.25** | **0.75** | **0.25** | **0.25** | 0.17 | 0.82 | 0 | 0.01 |
| **$\Theta_4$** | **2H1** | **H1:TTgg** | **0.25** | **0.75** | **0.25** | **0.5** | 0.16 | 0.84 | 0 | 0 |
| **$\Theta_4$** | **2H1** | **H1:TTgg** | **0.25** | **0.75** | **0.25** | **0.75** | 0.18 | 0.82 | 0 | 0 |
| **$\Theta_4$** | **2H1** | **H1:TTgg** | **0.25** | **0.75** | **0.5** | **0.25** | 0.29 | 0.71 | 0 | 0 |
| **$\Theta_4$** | **2H1** | **H1:TTgg** | **0.25** | **0.75** | **0.5** | **0.5** | 0.30 | 0.70 | 0 | 0 |
| **$\Theta_4$** | **2H1** | **H1:TTgg** | **0.25** | **0.75** | **0.5** | **0.75** | 0.32 | 0.68 | 0 | 0 |
| **$\Theta_4$** | **2H1** | **H1:TTgg** | **0.25** | **0.75** | **0.75** | **0.25** | 0.15 | 0.85 | 0 | 0 |
| **$\Theta_4$** | **2H1** | **H1:TTgg** | **0.25** | **0.75** | **0.75** | **0.5** | 0.17 | 0.83 | 0 | 0 |
| **$\Theta_4$** | **2H1** | **H1:TTgg** | **0.25** | **0.75** | **0.75** | **0.75** | 0.15 | 0.85 | 0 | 0 |
| **$\Theta_4$** | **2H1** | **H1:TTgg** | **0.375** | **0.125** | **0.25** | **0.25** | 0.04 | 0.96 | 0 | 0 |
| **$\Theta_4$** | **2H1** | **H1:TTgg** | **0.375** | **0.125** | **0.25** | **0.5** | 0.05 | 0.95 | 0 | 0 |
| **$\Theta_4$** | **2H1** | **H1:TTgg** | **0.375** | **0.125** | **0.25** | **0.75** | 0.04 | 0.96 | 0 | 0 |
| **$\Theta_4$** | **2H1** | **H1:TTgg** | **0.375** | **0.125** | **0.5** | **0.25** | 0.06 | 0.93 | 0 | 0.01 |
| **$\Theta_4$** | **2H1** | **H1:TTgg** | **0.375** | **0.125** | **0.5** | **0.5** | 0.17 | 0.83 | 0 | 0 |
| **$\Theta_4$** | **2H1** | **H1:TTgg** | **0.375** | **0.125** | **0.5** | **0.75** | 0.19 | 0.81 | 0 | 0 |
| **$\Theta_4$** | **2H1** | **H1:TTgg** | **0.375** | **0.125** | **0.75** | **0.25** | 0.03 | 0.96 | 0 | 0.01 |
| **$\Theta_4$** | **2H1** | **H1:TTgg** | **0.375** | **0.125** | **0.75** | **0.5** | 0.15 | 0.85 | 0 | 0 |
| **$\Theta_4$** | **2H1** | **H1:TTgg** | **0.375** | **0.125** | **0.75** | **0.75** | 0.22 | 0.78 | 0 | 0 |
| **$\Theta_4$** | **2H1** | **H1:TTgg** | **0.375** | **1.125** | **0.25** | **0.25** | 0.30 | 0.70 | 0 | 0 |
| **$\Theta_4$** | **2H1** | **H1:TTgg** | **0.375** | **1.125** | **0.25** | **0.5** | 0.33 | 0.67 | 0 | 0 |

| | | | | | | | | | | |
|---|---|---|---|---|---|---|---|---|---|---|
| **$\Theta_4$** | **2H1** | **H1:TTgg** | **0.375** | **1.125** | **0.25** | **0.75** | 0.30 | 0.70 | 0 | 0 |
| **$\Theta_4$** | **2H1** | **H1:TTgg** | **0.375** | **1.125** | **0.5** | **0.25** | 0.61 | 0.39 | 0 | 0 |
| **$\Theta_4$** | **2H1** | **H1:TTgg** | **0.375** | **1.125** | **0.5** | **0.5** | 0.61 | 0.39 | 0 | 0 |
| **$\Theta_4$** | **2H1** | **H1:TTgg** | **0.375** | **1.125** | **0.5** | **0.75** | 0.65 | 0.35 | 0 | 0 |
| **$\Theta_4$** | **2H1** | **H1:TTgg** | **0.375** | **1.125** | **0.75** | **0.25** | 0.30 | 0.70 | 0 | 0 |
| **$\Theta_4$** | **2H1** | **H1:TTgg** | **0.375** | **1.125** | **0.75** | **0.5** | 0.32 | 0.68 | 0 | 0 |
| **$\Theta_4$** | **2H1** | **H1:TTgg** | **0.375** | **1.125** | **0.75** | **0.75** | 0.29 | 0.71 | 0 | 0 |
| **$\Theta_4$** | **2H1** | **H1:TTgg** | **0.5** | **0.5** | **0.25** | **0.25** | 0.40 | 0.57 | 0 | 0.03 |
| **$\Theta_4$** | **2H1** | **H1:TTgg** | **0.5** | **0.5** | **0.25** | **0.5** | 0.50 | 0.50 | 0 | 0 |
| **$\Theta_4$** | **2H1** | **H1:TTgg** | **0.5** | **0.5** | **0.25** | **0.75** | 0.44 | 0.56 | 0 | 0 |
| **$\Theta_4$** | **2H1** | **H1:TTgg** | **0.5** | **0.5** | **0.5** | **0.25** | 0.74 | 0.25 | 0 | 0.01 |
| **$\Theta_4$** | **2H1** | **H1:TTgg** | **0.5** | **0.5** | **0.5** | **0.5** | 0.91 | 0.09 | 0 | 0 |
| **$\Theta_4$** | **2H1** | **H1:TTgg** | **0.5** | **0.5** | **0.5** | **0.75** | 0.90 | 0.10 | 0 | 0 |
| **$\Theta_4$** | **2H1** | **H1:TTgg** | **0.5** | **0.5** | **0.75** | **0.25** | 0.53 | 0.47 | 0 | 0 |
| **$\Theta_4$** | **2H1** | **H1:TTgg** | **0.5** | **0.5** | **0.75** | **0.5** | 0.59 | 0.41 | 0 | 0 |
| **$\Theta_4$** | **2H1** | **H1:TTgg** | **0.5** | **0.5** | **0.75** | **0.75** | 0.60 | 0.40 | 0 | 0 |
| **$\Theta_4$** | **2H1** | **H1:TTgg** | **0.75** | **0.25** | **0.25** | **0.25** | 0.19 | 0.78 | 0 | 0.03 |
| **$\Theta_4$** | **2H1** | **H1:TTgg** | **0.75** | **0.25** | **0.25** | **0.5** | 0.43 | 0.56 | 0 | 0.01 |
| **$\Theta_4$** | **2H1** | **H1:TTgg** | **0.75** | **0.25** | **0.25** | **0.75** | 0.43 | 0.57 | 0 | 0 |
| **$\Theta_4$** | **2H1** | **H1:TTgg** | **0.75** | **0.25** | **0.5** | **0.25** | 0.35 | 0.60 | 0 | 0.05 |
| **$\Theta_4$** | **2H1** | **H1:TTgg** | **0.75** | **0.25** | **0.5** | **0.5** | 0.79 | 0.20 | 0 | 0.01 |
| **$\Theta_4$** | **2H1** | **H1:TTgg** | **0.75** | **0.25** | **0.5** | **0.75** | 0.86 | 0.14 | 0 | 0 |
| **$\Theta_4$** | **2H1** | **H1:TTgg** | **0.75** | **0.25** | **0.75** | **0.25** | 0.49 | 0.47 | 0 | 0.04 |
| **$\Theta_4$** | **2H1** | **H1:TTgg** | **0.75** | **0.25** | **0.75** | **0.5** | 0.84 | 0.16 | 0 | 0 |
| **$\Theta_4$** | **2H1** | **H1:TTgg** | **0.75** | **0.25** | **0.75** | **0.75** | 0.93 | 0.07 | 0 | 0 |
| **$\Theta_4$** | **2H1** | **H1:TTgg** | **0.75** | **0.75** | **0.25** | **0.25** | 0.83 | 0.16 | 0 | 0.01 |
| **$\Theta_4$** | **2H1** | **H1:TTgg** | **0.75** | **0.75** | **0.25** | **0.5** | 0.88 | 0.12 | 0 | 0 |
| **$\Theta_4$** | **2H1** | **H1:TTgg** | **0.75** | **0.75** | **0.25** | **0.75** | 0.85 | 0.15 | 0 | 0 |
| **$\Theta_4$** | **2H1** | **H1:TTgg** | **0.75** | **0.75** | **0.5** | **0.25** | 0.98 | 0.02 | 0 | 0 |
| **$\Theta_4$** | **2H1** | **H1:TTgg** | **0.75** | **0.75** | **0.5** | **0.5** | 1 | 0 | 0 | 0 |
| **$\Theta_4$** | **2H1** | **H1:TTgg** | **0.75** | **0.75** | **0.5** | **0.75** | 1 | 0 | 0 | 0 |
| **$\Theta_4$** | **2H1** | **H1:TTgg** | **0.75** | **0.75** | **0.75** | **0.25** | 0.86 | 0.14 | 0 | 0 |
| **$\Theta_4$** | **2H1** | **H1:TTgg** | **0.75** | **0.75** | **0.75** | **0.5** | 0.87 | 0.13 | 0 | 0 |
| **$\Theta_4$** | **2H1** | **H1:TTgg** | **0.75** | **0.75** | **0.75** | **0.75** | 0.88 | 0.12 | 0 | 0 |
| **$\Theta_4$** | **2H1** | **H1:TTgg** | **1.125** | **0.375** | **0.25** | **0.25** | 0.47 | 0.49 | 0 | 0.04 |
| **$\Theta_4$** | **2H1** | **H1:TTgg** | **1.125** | **0.375** | **0.25** | **0.5** | 0.83 | 0.17 | 0 | 0 |
| **$\Theta_4$** | **2H1** | **H1:TTgg** | **1.125** | **0.375** | **0.25** | **0.75** | 0.78 | 0.22 | 0 | 0 |
| **$\Theta_4$** | **2H1** | **H1:TTgg** | **1.125** | **0.375** | **0.5** | **0.25** | 0.71 | 0.25 | 0 | 0.04 |
| **$\Theta_4$** | **2H1** | **H1:TTgg** | **1.125** | **0.375** | **0.5** | **0.5** | 0.99 | 0.01 | 0 | 0 |
| **$\Theta_4$** | **2H1** | **H1:TTgg** | **1.125** | **0.375** | **0.5** | **0.75** | 0.97 | 0.03 | 0 | 0 |
| **$\Theta_4$** | **2H1** | **H1:TTgg** | **1.125** | **0.375** | **0.75** | **0.25** | 0.82 | 0.16 | 0 | 0.02 |
| **$\Theta_4$** | **2H1** | **H1:TTgg** | **1.125** | **0.375** | **0.75** | **0.5** | 0.99 | 0.01 | 0 | 0 |
| **$\Theta_4$** | **2H1** | **H1:TTgg** | **1.125** | **0.375** | **0.75** | **0.75** | 1 | 0 | 0 | 0 |
| **$\Theta_4$** | **2H2** | **H2:TTgg** | **0.125** | **0.375** | **0.25** | **0.25** | 0.05 | 0.95 | 0 | 0 |
| **$\Theta_4$** | **2H2** | **H2:TTgg** | **0.125** | **0.375** | **0.25** | **0.5** | 0.05 | 0.95 | 0 | 0 |
| **$\Theta_4$** | **2H2** | **H2:TTgg** | **0.125** | **0.375** | **0.25** | **0.75** | 0.05 | 0.95 | 0 | 0 |
| **$\Theta_4$** | **2H2** | **H2:TTgg** | **0.125** | **0.375** | **0.5** | **0.25** | 0.05 | 0.95 | 0 | 0 |

| | | | | | | | | | | |
|---|---|---|---|---|---|---|---|---|---|---|
| **$\Theta_4$** | **2H2** | **H2:TTgg** | **0.125** | **0.375** | **0.5** | **0.5** | 0.06 | 0.94 | 0 | 0 |
| **$\Theta_4$** | **2H2** | **H2:TTgg** | **0.125** | **0.375** | **0.5** | **0.75** | 0.07 | 0.93 | 0 | 0 |
| **$\Theta_4$** | **2H2** | **H2:TTgg** | **0.25** | **0.25** | **0.25** | **0.25** | 0.07 | 0.93 | 0 | 0 |
| **$\Theta_4$** | **2H2** | **H2:TTgg** | **0.25** | **0.25** | **0.25** | **0.5** | 0.09 | 0.91 | 0 | 0 |
| **$\Theta_4$** | **2H2** | **H2:TTgg** | **0.25** | **0.25** | **0.25** | **0.75** | 0.09 | 0.91 | 0 | 0 |
| **$\Theta_4$** | **2H2** | **H2:TTgg** | **0.25** | **0.25** | **0.5** | **0.25** | 0.20 | 0.80 | 0 | 0 |
| **$\Theta_4$** | **2H2** | **H2:TTgg** | **0.25** | **0.25** | **0.5** | **0.5** | 0.18 | 0.82 | 0 | 0 |
| **$\Theta_4$** | **2H2** | **H2:TTgg** | **0.25** | **0.25** | **0.5** | **0.75** | 0.2 | 0.8 | 0 | 0 |
| **$\Theta_4$** | **2H2** | **H2:TTgg** | **0.25** | **0.75** | **0.25** | **0.25** | 0.16 | 0.84 | 0 | 0 |
| **$\Theta_4$** | **2H2** | **H2:TTgg** | **0.25** | **0.75** | **0.25** | **0.5** | 0.17 | 0.83 | 0 | 0 |
| **$\Theta_4$** | **2H2** | **H2:TTgg** | **0.25** | **0.75** | **0.25** | **0.75** | 0.16 | 0.84 | 0 | 0 |
| **$\Theta_4$** | **2H2** | **H2:TTgg** | **0.25** | **0.75** | **0.5** | **0.25** | 0.3 | 0.7 | 0 | 0 |
| **$\Theta_4$** | **2H2** | **H2:TTgg** | **0.25** | **0.75** | **0.5** | **0.5** | 0.32 | 0.68 | 0 | 0 |
| **$\Theta_4$** | **2H2** | **H2:TTgg** | **0.25** | **0.75** | **0.5** | **0.75** | 0.3 | 0.7 | 0 | 0 |
| **$\Theta_4$** | **2H2** | **H2:TTgg** | **0.375** | **0.125** | **0.25** | **0.25** | 0.03 | 0.97 | 0 | 0 |
| **$\Theta_4$** | **2H2** | **H2:TTgg** | **0.375** | **0.125** | **0.25** | **0.5** | 0.05 | 0.94 | 0 | 0.01 |
| **$\Theta_4$** | **2H2** | **H2:TTgg** | **0.375** | **0.125** | **0.25** | **0.75** | 0.06 | 0.93 | 0 | 0.01 |
| **$\Theta_4$** | **2H2** | **H2:TTgg** | **0.375** | **0.125** | **0.5** | **0.25** | 0.14 | 0.86 | 0 | 0 |
| **$\Theta_4$** | **2H2** | **H2:TTgg** | **0.375** | **0.125** | **0.5** | **0.5** | 0.14 | 0.86 | 0 | 0 |
| **$\Theta_4$** | **2H2** | **H2:TTgg** | **0.375** | **0.125** | **0.5** | **0.75** | 0.14 | 0.85 | 0 | 0.01 |
| **$\Theta_4$** | **2H2** | **H2:TTgg** | **0.375** | **1.125** | **0.25** | **0.25** | 0.29 | 0.71 | 0 | 0 |
| **$\Theta_4$** | **2H2** | **H2:TTgg** | **0.375** | **1.125** | **0.25** | **0.5** | 0.29 | 0.71 | 0 | 0 |
| **$\Theta_4$** | **2H2** | **H2:TTgg** | **0.375** | **1.125** | **0.25** | **0.75** | 0.29 | 0.71 | 0 | 0 |
| **$\Theta_4$** | **2H2** | **H2:TTgg** | **0.375** | **1.125** | **0.5** | **0.25** | 0.64 | 0.36 | 0 | 0 |
| **$\Theta_4$** | **2H2** | **H2:TTgg** | **0.375** | **1.125** | **0.5** | **0.5** | 0.62 | 0.38 | 0 | 0 |
| **$\Theta_4$** | **2H2** | **H2:TTgg** | **0.375** | **1.125** | **0.5** | **0.75** | 0.6 | 0.4 | 0 | 0 |
| **$\Theta_4$** | **2H2** | **H2:TTgg** | **0.5** | **0.5** | **0.25** | **0.25** | 0.58 | 0.42 | 0 | 0 |
| **$\Theta_4$** | **2H2** | **H2:TTgg** | **0.5** | **0.5** | **0.25** | **0.5** | 0.57 | 0.43 | 0 | 0 |
| **$\Theta_4$** | **2H2** | **H2:TTgg** | **0.5** | **0.5** | **0.25** | **0.75** | 0.55 | 0.45 | 0 | 0 |
| **$\Theta_4$** | **2H2** | **H2:TTgg** | **0.5** | **0.5** | **0.5** | **0.25** | 0.94 | 0.06 | 0 | 0 |
| **$\Theta_4$** | **2H2** | **H2:TTgg** | **0.5** | **0.5** | **0.5** | **0.5** | 0.94 | 0.06 | 0 | 0 |
| **$\Theta_4$** | **2H2** | **H2:TTgg** | **0.5** | **0.5** | **0.5** | **0.75** | 0.93 | 0.07 | 0 | 0 |
| **$\Theta_4$** | **2H2** | **H2:TTgg** | **0.75** | **0.25** | **0.25** | **0.25** | 0.68 | 0.32 | 0 | 0 |
| **$\Theta_4$** | **2H2** | **H2:TTgg** | **0.75** | **0.25** | **0.25** | **0.5** | 0.61 | 0.39 | 0 | 0 |
| **$\Theta_4$** | **2H2** | **H2:TTgg** | **0.75** | **0.25** | **0.25** | **0.75** | 0.52 | 0.47 | 0 | 0.01 |
| **$\Theta_4$** | **2H2** | **H2:TTgg** | **0.75** | **0.25** | **0.5** | **0.25** | 0.68 | 0.32 | 0 | 0 |
| **$\Theta_4$** | **2H2** | **H2:TTgg** | **0.75** | **0.25** | **0.5** | **0.5** | 0.68 | 0.32 | 0 | 0 |
| **$\Theta_4$** | **2H2** | **H2:TTgg** | **0.75** | **0.25** | **0.5** | **0.75** | 0.66 | 0.33 | 0 | 0.01 |
| **$\Theta_4$** | **2H2** | **H2:TTgg** | **0.75** | **0.75** | **0.25** | **0.25** | 0.87 | 0.13 | 0 | 0 |
| **$\Theta_4$** | **2H2** | **H2:TTgg** | **0.75** | **0.75** | **0.25** | **0.5** | 0.87 | 0.13 | 0 | 0 |
| **$\Theta_4$** | **2H2** | **H2:TTgg** | **0.75** | **0.75** | **0.25** | **0.75** | 0.82 | 0.18 | 0 | 0 |
| **$\Theta_4$** | **2H2** | **H2:TTgg** | **0.75** | **0.75** | **0.5** | **0.25** | 1 | 0 | 0 | 0 |
| **$\Theta_4$** | **2H2** | **H2:TTgg** | **0.75** | **0.75** | **0.5** | **0.5** | 1 | 0 | 0 | 0 |
| **$\Theta_4$** | **2H2** | **H2:TTgg** | **0.75** | **0.75** | **0.5** | **0.75** | 0.94 | 0.06 | 0 | 0 |
| **$\Theta_4$** | **2H2** | **H2:TTgg** | **1.125** | **0.375** | **0.25** | **0.25** | 0.91 | 0.09 | 0 | 0 |
| **$\Theta_4$** | **2H2** | **H2:TTgg** | **1.125** | **0.375** | **0.25** | **0.5** | 0.89 | 0.11 | 0 | 0 |
| **$\Theta_4$** | **2H2** | **H2:TTgg** | **1.125** | **0.375** | **0.25** | **0.75** | 0.83 | 0.16 | 0 | 0.01 |
| **$\Theta_4$** | **2H2** | **H2:TTgg** | **1.125** | **0.375** | **0.5** | **0.25** | 0.87 | 0.13 | 0 | 0 |
| **$\Theta_4$** | **2H2** | **H2:TTgg** | **1.125** | **0.375** | **0.5** | **0.5** | 0.87 | 0.13 | 0 | 0 |
| **$\Theta_4$** | **2H2** | **H2:TTgg** | **1.125** | **0.375** | **0.5** | **0.75** | 0.85 | 0.15 | 0 | 0 |

### Table S4.

**Testing the *stepwise* method and chi-square distribution criterion for frequency of correct recognition of models in the Poisson randomized set.** The first column indicates group *Θj*, the following two columns show the trivial name of the model and its alias in TTgg notation, and the following four columns describe the initial values of parameters T1, T3, $\gamma_1$, $\gamma_3$ taken in a Poisson-distribution based randomization process. The last four columns present the frequency of the correct model recognition: $\Theta_j^i$; frequency of error recognition of the model in the direction of a simple model: *Θj-*; frequency of error recognition of the model to a complex model: *Θj+*; frequency of error recognition of the model because optimization process selects another model from the same group: *Θj*.

| Group | Model | Alias | $T_1$ | $T_3$ | $\gamma_1$ | $\gamma_3$ | $\Theta_j^i$ | $\Theta_{j-}$ | $\Theta_{j+}$ | $\Theta_j$ |
|---|---|---|---|---|---|---|---|---|---|---|
| $\Theta_0$ | P | H1:00nn | 0 | 0 | - | - | 0.76 | 0 | 0.24 | 0 |
| $\Theta_1$ | TP | H1:0Tn1 | 0 | 0.5 | - | 1 | 0.85 | 0 | 0.15 | 0 |
| $\Theta_1$ | TP | H1:0Tn1 | 0 | 1 | - | 1 | 0.82 | 0 | 0.18 | 0 |
| $\Theta_1$ | TP | H1:0Tn1 | 0 | 1.5 | - | 1 | 0.82 | 0 | 0.18 | 0 |
| $\Theta_1$ | PT | H1:T00n | 0.5 | 0 | 0 | - | 0.85 | 0 | 0.15 | 0 |
| $\Theta_1$ | PT | H1:T00n | 1 | 0 | 0 | - | 0.85 | 0 | 0.15 | 0 |
| $\Theta_1$ | PT | H1:T00n | 1.5 | 0 | 0 | - | 0.84 | 0 | 0.16 | 0 |
| $\Theta_2$ | 1HP1 | H1:0Tng | 0 | 0.5 | - | 0.25 | 0.76 | 0.13 | 0.08 | 0.03 |
| $\Theta_2$ | 1HP1 | H1:0Tng | 0 | 0.5 | - | 0.5 | 0.88 | 0 | 0.11 | 0.01 |
| $\Theta_2$ | 1HP1 | H1:0Tng | 0 | 1 | - | 0.25 | 0.92 | 0 | 0.08 | 0 |
| $\Theta_2$ | 1HP1 | H1:0Tng | 0 | 1 | - | 0.5 | 0.92 | 0 | 0.08 | 0 |
| $\Theta_2$ | 1HP1 | H1:0Tng | 0 | 1.5 | - | 0.25 | 0.91 | 0 | 0.09 | 0 |
| $\Theta_2$ | 1HP1 | H1:0Tng | 0 | 1.5 | - | 0.5 | 0.91 | 0 | 0.09 | 0 |
| $\Theta_2$ | 1HP2 | H1:T0g0 | 0.5 | 0 | 0.25 | 0 | 0.80 | 0.01 | 0.09 | 0.10 |
| $\Theta_2$ | 1HP2 | H1:T0g0 | 0.5 | 0 | 0.5 | 0 | 0.85 | 0 | 0.15 | 0 |
| $\Theta_2$ | 1HP2 | H1:T0g0 | 0.5 | 0 | 0.75 | 0 | 0.83 | 0 | 0.12 | 0.05 |
| $\Theta_2$ | 1HP2 | H1:T0g0 | 1 | 0 | 0.25 | 0 | 0.86 | 0 | 0.14 | 0 |
| $\Theta_2$ | 1HP2 | H1:T0g0 | 1 | 0 | 0.5 | 0 | 0.89 | 0 | 0.11 | 0 |
| $\Theta_2$ | 1HP2 | H1:T0g0 | 1 | 0 | 0.75 | 0 | 0.86 | 0 | 0.14 | 0 |
| $\Theta_2$ | 1HP2 | H1:T0g0 | 1.5 | 0 | 0.25 | 0 | 0.86 | 0 | 0.14 | 0 |
| $\Theta_2$ | 1HP2 | H1:T0g0 | 1.5 | 0 | 0.5 | 0 | 0.89 | 0 | 0.11 | 0 |
| $\Theta_2$ | 1HP2 | H1:T0g0 | 1.5 | 0 | 0.75 | 0 | 0.88 | 0 | 0.12 | 0 |
| $\Theta_2$ | 1HP3 | H1:T0g1 | 0.5 | 0 | 0.5 | 1 | 0.83 | 0.03 | 0.08 | 0.06 |
| $\Theta_2$ | 1HP3 | H1:T0g1 | 0.5 | 0 | 0.25 | 1 | 0.88 | 0 | 0.11 | 0.01 |
| $\Theta_2$ | 1HP3 | H1:T0g1 | 1 | 0 | 0.5 | 1 | 0.88 | 0 | 0.12 | 0 |
| $\Theta_2$ | 1HP3 | H1:T0g1 | 1 | 0 | 0.25 | 1 | 0.88 | 0 | 0.12 | 0 |
| $\Theta_2$ | 1HP3 | H1:T0g1 | 1.5 | 0 | 0.5 | 1 | 0.88 | 0 | 0.12 | 0 |
| $\Theta_2$ | 1HP3 | H1:T0g1 | 1.5 | 0 | 0.25 | 1 | 0.88 | 0 | 0.12 | 0 |
| $\Theta_2$ | T1 | H1:TT10 | 0.125 | 0.375 | 1 | 0 | 0.53 | 0.31 | 0.11 | 0.05 |
| $\Theta_2$ | T1 | H1:TT10 | 0.25 | 0.25 | 1 | 0 | 0.74 | 0 | 0.10 | 0.16 |
| $\Theta_2$ | T1 | H1:TT10 | 0.25 | 0.75 | 1 | 0 | 0.84 | 0.01 | 0.15 | 0 |
| $\Theta_2$ | T1 | H1:TT10 | 0.375 | 0.125 | 1 | 0 | 0.49 | 0 | 0.05 | 0.46 |
| $\Theta_2$ | T1 | H1:TT10 | 0.375 | 1.125 | 1 | 0 | 0.88 | 0 | 0.12 | 0 |
| $\Theta_2$ | T1 | H1:TT10 | 0.5 | 0.5 | 1 | 0 | 0.87 | 0 | 0.12 | 0.01 |
| $\Theta_2$ | T1 | H1:TT10 | 0.75 | 0.25 | 1 | 0 | 0.63 | 0 | 0.09 | 0.28 |
| $\Theta_2$ | T1 | H1:TT10 | 0.75 | 0.75 | 1 | 0 | 0.85 | 0 | 0.15 | 0 |
| $\Theta_2$ | T1 | H1:TT10 | 1.125 | 0.375 | 1 | 0 | 0.75 | 0 | 0.07 | 0.18 |
| $\Theta_2$ | T2 | H1:TT01 | 0.125 | 0.375 | 0 | 1 | 0.56 | 0.28 | 0.08 | 0.08 |

| | | | | | | | | | | |
|---|---|---|---|---|---|---|---|---|---|---|
| $\Theta_2$ | T2 | H1:TT01 | 0.25 | 0.25 | 0 | 1 | 0.84 | 0 | 0.10 | 0.06 |
| $\Theta_2$ | T2 | H1:TT01 | 0.25 | 0.75 | 0 | 1 | 0.87 | 0.02 | 0.11 | 0 |
| $\Theta_2$ | T2 | H1:TT01 | 0.375 | 0.125 | 0 | 1 | 0.52 | 0.23 | 0.08 | 0.17 |
| $\Theta_2$ | T2 | H1:TT01 | 0.375 | 1.125 | 0 | 1 | 0.89 | 0 | 0.11 | 0 |
| $\Theta_2$ | T2 | H1:TT01 | 0.5 | 0.5 | 0 | 1 | 0.87 | 0 | 0.13 | 0 |
| $\Theta_2$ | T2 | H1:TT01 | 0.75 | 0.25 | 0 | 1 | 0.86 | 0.01 | 0.10 | 0.03 |
| $\Theta_2$ | T2 | H1:TT01 | 0.75 | 0.75 | 0 | 1 | 0.87 | 0 | 0.13 | 0 |
| $\Theta_3$ | T2 | H1:TT01 | 1.125 | 0.375 | 0 | 0 | 0.87 | 0 | 0.13 | 0 |
| $\Theta_3$ | 1H1 | H1:TTg0 | 0.125 | 0.375 | 0.25 | 0 | 0.04 | 0.93 | 0.01 | 0.02 |
| $\Theta_3$ | 1H1 | H1:TTg0 | 0.125 | 0.375 | 0.5 | 0 | 0.06 | 0.91 | 0.02 | 0.01 |
| $\Theta_3$ | 1H1 | H1:TTg0 | 0.125 | 0.375 | 0.75 | 0 | 0.04 | 0.91 | 0.01 | 0.04 |
| $\Theta_3$ | 1H1 | H1:TTg0 | 0.25 | 0.25 | 0.25 | 0 | 0.07 | 0.86 | 0.04 | 0.03 |
| $\Theta_3$ | 1H1 | H1:TTg0 | 0.25 | 0.25 | 0.5 | 0 | 0.19 | 0.73 | 0.04 | 0.04 |
| $\Theta_3$ | 1H1 | H1:TTg0 | 0.25 | 0.25 | 0.75 | 0 | 0.08 | 0.82 | 0.02 | 0.08 |
| $\Theta_3$ | 1H1 | H1:TTg0 | 0.25 | 0.75 | 0.25 | 0 | 0.19 | 0.75 | 0.03 | 0.03 |
| $\Theta_3$ | 1H1 | H1:TTg0 | 0.25 | 0.75 | 0.5 | 0 | 0.33 | 0.62 | 0.03 | 0.02 |
| $\Theta_3$ | 1H1 | H1:TTg0 | 0.25 | 0.75 | 0.75 | 0 | 0.15 | 0.76 | 0.02 | 0.07 |
| $\Theta_3$ | 1H1 | H1:TTg0 | 0.375 | 0.125 | 0.25 | 0 | 0.02 | 0.92 | 0.02 | 0.04 |
| $\Theta_3$ | 1H1 | H1:TTg0 | 0.375 | 0.125 | 0.5 | 0 | 0.12 | 0.79 | 0.06 | 0.03 |
| $\Theta_3$ | 1H1 | H1:TTg0 | 0.375 | 0.125 | 0.75 | 0 | 0.05 | 0.87 | 0.02 | 0.06 |
| $\Theta_3$ | 1H1 | H1:TTg0 | 0.375 | 1.125 | 0.25 | 0 | 0.31 | 0.62 | 0.03 | 0.04 |
| $\Theta_3$ | 1H1 | H1:TTg0 | 0.375 | 1.125 | 0.5 | 0 | 0.63 | 0.30 | 0.05 | 0.02 |
| $\Theta_3$ | 1H1 | H1:TTg0 | 0.375 | 1.125 | 0.75 | 0 | 0.28 | 0.62 | 0.04 | 0.06 |
| $\Theta_3$ | 1H1 | H1:TTg0 | 0.5 | 0.5 | 0.25 | 0 | 0.57 | 0.35 | 0.07 | 0.01 |
| $\Theta_3$ | 1H1 | H1:TTg0 | 0.5 | 0.5 | 0.5 | 0 | 0.86 | 0.04 | 0.10 | 0 |
| $\Theta_3$ | 1H1 | H1:TTg0 | 0.5 | 0.5 | 0.75 | 0 | 0.55 | 0.33 | 0.06 | 0.06 |
| $\Theta_3$ | 1H1 | H1:TTg0 | 0.75 | 0.25 | 0.25 | 0 | 0.63 | 0.27 | 0.09 | 0.01 |
| $\Theta_3$ | 1H1 | H1:TTg0 | 0.75 | 0.25 | 0.5 | 0 | 0.57 | 0.35 | 0.08 | 0 |
| $\Theta_3$ | 1H1 | H1:TTg0 | 0.75 | 0.25 | 0.75 | 0 | 0.45 | 0.47 | 0.06 | 0.02 |
| $\Theta_3$ | 1H1 | H1:TTg0 | 0.75 | 0.75 | 0.25 | 0 | 0.84 | 0.06 | 0.09 | 0.01 |
| $\Theta_3$ | 1H1 | H1:TTg0 | 0.75 | 0.75 | 0.5 | 0 | 0.91 | 0 | 0.09 | 0 |
| $\Theta_3$ | 1H1 | H1:TTg0 | 0.75 | 0.75 | 0.75 | 0 | 0.80 | 0.11 | 0.07 | 0.02 |
| $\Theta_3$ | 1H1 | H1:TTg0 | 1.125 | 0.375 | 0.25 | 0 | 0.85 | 0.06 | 0.09 | 0 |
| $\Theta_3$ | 1H1 | H1:TTg0 | 1.125 | 0.375 | 0.5 | 0 | 0.76 | 0.16 | 0.08 | 0 |
| $\Theta_3$ | 1H1 | H1:TTg0 | 1.125 | 0.375 | 0.75 | 0 | 0.63 | 0.29 | 0.07 | 0.01 |
| $\Theta_3$ | 1H2 | H1:TT1g | 0.125 | 0.375 | 1 | 0.25 | 0.43 | 0.53 | 0.03 | 0.01 |
| $\Theta_3$ | 1H2 | H1:TT1g | 0.125 | 0.375 | 1 | 0.5 | 0.59 | 0.37 | 0.04 | 0 |
| $\Theta_3$ | 1H2 | H1:TT1g | 0.125 | 0.375 | 1 | 0.75 | 0.47 | 0.47 | 0.04 | 0.02 |
| $\Theta_3$ | 1H2 | H1:TT1g | 0.25 | 0.25 | 1 | 0.25 | 0.55 | 0.39 | 0.02 | 0.04 |
| $\Theta_3$ | 1H2 | H1:TT1g | 0.25 | 0.25 | 1 | 0.5 | 0.82 | 0.11 | 0.03 | 0.04 |
| $\Theta_3$ | 1H2 | H1:TT1g | 0.25 | 0.25 | 1 | 0.75 | 0.67 | 0.28 | 0.02 | 0.03 |
| $\Theta_3$ | 1H2 | H1:TT1g | 0.25 | 0.75 | 1 | 0.25 | 0.95 | 0.02 | 0.03 | 0 |
| $\Theta_3$ | 1H2 | H1:TT1g | 0.25 | 0.75 | 1 | 0.5 | 0.93 | 0.02 | 0.05 | 0 |
| $\Theta_3$ | 1H2 | H1:TT1g | 0.25 | 0.75 | 1 | 0.75 | 0.93 | 0.02 | 0.05 | 0 |
| $\Theta_3$ | 1H2 | H1:TT1g | 0.375 | 0.125 | 1 | 0.25 | 0.24 | 0.70 | 0 | 0.06 |
| $\Theta_3$ | 1H2 | H1:TT1g | 0.375 | 0.125 | 1 | 0.5 | 0.31 | 0.59 | 0.03 | 0.07 |
| $\Theta_3$ | 1H2 | H1:TT1g | 0.375 | 0.125 | 1 | 0.75 | 0.37 | 0.56 | 0.02 | 0.05 |
| $\Theta_3$ | 1H2 | H1:TT1g | 0.375 | 1.125 | 1 | 0.25 | 0.96 | 0 | 0.04 | 0 |
| $\Theta_3$ | 1H2 | H1:TT1g | 0.375 | 1.125 | 1 | 0.5 | 0.95 | 0 | 0.05 | 0 |
| $\Theta_3$ | 1H2 | H1:TT1g | 0.375 | 1.125 | 1 | 0.75 | 0.96 | 0 | 0.04 | 0 |

| | | | | | | | | | | |
|---|---|---|---|---|---|---|---|---|---|---|
| $\Theta_3$ | 1H2 | H1:TT1g | 0.5 | 0.5 | 1 | 0.25 | 0.95 | 0 | 0.05 | 0 |
| $\Theta_3$ | 1H2 | H1:TT1g | 0.5 | 0.5 | 1 | 0.5 | 0.95 | 0 | 0.05 | 0 |
| $\Theta_3$ | 1H2 | H1:TT1g | 0.5 | 0.5 | 1 | 0.75 | 0.95 | 0 | 0.05 | 0 |
| $\Theta_3$ | 1H2 | H1:TT1g | 0.75 | 0.25 | 1 | 0.25 | 0.61 | 0.3 | 0.02 | 0.07 |
| $\Theta_3$ | 1H2 | H1:TT1g | 0.75 | 0.25 | 1 | 0.5 | 0.77 | 0.16 | 0.03 | 0.04 |
| $\Theta_3$ | 1H2 | H1:TT1g | 0.75 | 0.25 | 1 | 0.75 | 0.86 | 0.04 | 0.05 | 0.05 |
| $\Theta_3$ | 1H2 | H1:TT1g | 0.75 | 0.75 | 1 | 0.25 | 0.96 | 0 | 0.04 | 0 |
| $\Theta_3$ | 1H2 | H1:TT1g | 0.75 | 0.75 | 1 | 0.5 | 0.94 | 0 | 0.06 | 0 |
| $\Theta_3$ | 1H2 | H1:TT1g | 0.75 | 0.75 | 1 | 0.75 | 0.95 | 0 | 0.05 | 0 |
| $\Theta_3$ | 1H2 | H1:TT1g | 1.125 | 0.375 | 1 | 0.25 | 0.81 | 0.14 | 0.03 | 0.02 |
| $\Theta_3$ | 1H2 | H1:TT1g | 1.125 | 0.375 | 1 | 0.5 | 0.90 | 0.03 | 0.06 | 0.01 |
| $\Theta_3$ | 1H2 | H1:TT1g | 1.125 | 0.375 | 1 | 0.75 | 0.93 | 0.01 | 0.05 | 0.01 |
| $\Theta_3$ | 1H3 | H1:TT0g | 0.125 | 0.375 | 0 | 0.25 | 0.19 | 0.71 | 0.02 | 0.08 |
| $\Theta_3$ | 1H3 | H1:TT0g | 0.125 | 0.375 | 0 | 0.5 | 0.40 | 0.51 | 0.03 | 0.06 |
| $\Theta_3$ | 1H3 | H1:TT0g | 0.25 | 0.25 | 0 | 0.25 | 0.16 | 0.75 | 0.01 | 0.08 |
| $\Theta_3$ | 1H3 | H1:TT0g | 0.25 | 0.25 | 0 | 0.5 | 0.28 | 0.63 | 0.01 | 0.08 |
| $\Theta_3$ | 1H3 | H1:TT0g | 0.25 | 0.75 | 0 | 0.25 | 0.89 | 0.05 | 0.04 | 0.02 |
| $\Theta_3$ | 1H3 | H1:TT0g | 0.25 | 0.75 | 0 | 0.5 | 0.94 | 0.01 | 0.05 | 0 |
| $\Theta_3$ | 1H3 | H1:TT0g | 0.375 | 0.125 | 0 | 0.25 | 0.05 | 0.90 | 0 | 0.05 |
| $\Theta_3$ | 1H3 | H1:TT0g | 0.375 | 0.125 | 0 | 0.5 | 0.05 | 0.91 | 0 | 0.04 |
| $\Theta_3$ | 1H3 | H1:TT0g | 0.375 | 1.125 | 0 | 0.25 | 0.94 | 0 | 0.06 | 0 |
| $\Theta_3$ | 1H3 | H1:TT0g | 0.375 | 1.125 | 0 | 0.5 | 0.94 | 0 | 0.06 | 0 |
| $\Theta_3$ | 1H3 | H1:TT0g | 0.5 | 0.5 | 0 | 0.25 | 0.61 | 0.29 | 0.03 | 0.07 |
| $\Theta_3$ | 1H3 | H1:TT0g | 0.5 | 0.5 | 0 | 0.5 | 0.89 | 0.04 | 0.04 | 0.03 |
| $\Theta_3$ | 1H3 | H1:TT0g | 0.75 | 0.25 | 0 | 0.25 | 0.15 | 0.76 | 0.01 | 0.08 |
| $\Theta_3$ | 1H3 | H1:TT0g | 0.75 | 0.25 | 0 | 0.5 | 0.29 | 0.63 | 0.02 | 0.06 |
| $\Theta_3$ | 1H3 | H1:TT0g | 0.75 | 0.75 | 0 | 0.25 | 0.87 | 0.05 | 0.04 | 0.04 |
| $\Theta_3$ | 1H3 | H1:TT0g | 0.75 | 0.75 | 0 | 0.5 | 0.95 | 0 | 0.05 | 0 |
| $\Theta_3$ | 1H3 | H1:TT0g | 1.125 | 0.375 | 0 | 0.25 | 0.28 | 0.62 | 0.02 | 0.08 |
| $\Theta_3$ | 1H3 | H1:TT0g | 1.125 | 0.375 | 0 | 0.5 | 0.60 | 0.31 | 0.03 | 0.06 |
| $\Theta_3$ | 1H4 | H1:TTg1 | 0.125 | 0.375 | 0.25 | 1 | 0.06 | 0.91 | 0 | 0.03 |
| $\Theta_3$ | 1H4 | H1:TTg1 | 0.125 | 0.375 | 0.5 | 1 | 0.08 | 0.9 | 0 | 0.02 |
| $\Theta_3$ | 1H4 | H1:TTg1 | 0.25 | 0.25 | 0.25 | 1 | 0.20 | 0.70 | 0.01 | 0.09 |
| $\Theta_3$ | 1H4 | H1:TTg1 | 0.25 | 0.25 | 0.5 | 1 | 0.44 | 0.50 | 0.01 | 0.05 |
| $\Theta_3$ | 1H4 | H1:TTg1 | 0.25 | 0.75 | 0.25 | 1 | 0.17 | 0.72 | 0 | 0.11 |
| $\Theta_3$ | 1H4 | H1:TTg1 | 0.25 | 0.75 | 0.5 | 1 | 0.33 | 0.61 | 0.02 | 0.04 |
| $\Theta_3$ | 1H4 | H1:TTg1 | 0.375 | 0.125 | 0.25 | 1 | 0.36 | 0.57 | 0.01 | 0.06 |
| $\Theta_3$ | 1H4 | H1:TTg1 | 0.375 | 0.125 | 0.5 | 1 | 0.69 | 0.26 | 0.03 | 0.02 |
| $\Theta_3$ | 1H4 | H1:TTg1 | 0.375 | 1.125 | 0.25 | 1 | 0.31 | 0.55 | 0.01 | 0.13 |
| $\Theta_3$ | 1H4 | H1:TTg1 | 0.375 | 1.125 | 0.5 | 1 | 0.65 | 0.28 | 0.03 | 0.04 |
| $\Theta_3$ | 1H4 | H1:TTg1 | 0.5 | 0.5 | 0.25 | 1 | 0.63 | 0.25 | 0.03 | 0.09 |
| $\Theta_3$ | 1H4 | H1:TTg1 | 0.5 | 0.5 | 0.5 | 1 | 0.90 | 0.01 | 0.08 | 0.01 |
| $\Theta_3$ | 1H4 | H1:TTg1 | 0.75 | 0.25 | 0.25 | 1 | 0.90 | 0.03 | 0.05 | 0.02 |
| $\Theta_3$ | 1H4 | H1:TTg1 | 0.75 | 0.25 | 0.5 | 1 | 0.93 | 0 | 0.07 | 0 |
| $\Theta_3$ | 1H4 | H1:TTg1 | 0.75 | 0.75 | 0.25 | 1 | 0.87 | 0.05 | 0.05 | 0.03 |
| $\Theta_3$ | 1H4 | H1:TTg1 | 0.75 | 0.75 | 0.5 | 1 | 0.92 | 0 | 0.08 | 0 |
| $\Theta_3$ | 1H4 | H1:TTg1 | 1.125 | 0.375 | 0.25 | 1 | 0.93 | 0 | 0.07 | 0 |
| $\Theta_3$ | 1H4 | H1:TTg1 | 1.125 | 0.375 | 0.5 | 1 | 0.91 | 0 | 0.09 | 0 |
| $\Theta_3$ | 2PH | H2:T0gg | 0.5 | 0 | 0.25 | 0.5 | 0.62 | 0.27 | 0.04 | 0.07 |
| $\Theta_3$ | 2PH | H2:T0gg | 0.5 | 0 | 0.25 | 0.75 | 0.48 | 0.39 | 0.03 | 0.1 |

| | | | | | | | | | | |
|---|---|---|---|---|---|---|---|---|---|---|
| **$\Theta_3$** | **2PH** | **H2:T0gg** | **1** | **0** | **0.25** | **0.5** | 0.93 | 0 | 0.07 | 0 |
| **$\Theta_3$** | **2PH** | **H2:T0gg** | **1** | **0** | **0.25** | **0.75** | 0.89 | 0 | 0.11 | 0 |
| **$\Theta_3$** | **2PH** | **H2:T0gg** | **1.5** | **0** | **0.25** | **0.5** | 0.92 | 0 | 0.08 | 0 |
| **$\Theta_3$** | **2PH** | **H2:T0gg** | **1.5** | **0** | **0.25** | **0.75** | 0.92 | 0 | 0.08 | 0 |
| **$\Theta_4$** | **2H1** | **H1:TTgg** | **0.125** | **0.375** | **0.25** | **0.25** | 0.02 | 0.97 | 0 | 0.01 |
| **$\Theta_4$** | **2H1** | **H1:TTgg** | **0.125** | **0.375** | **0.25** | **0.5** | 0.04 | 0.96 | 0 | 0 |
| **$\Theta_4$** | **2H1** | **H1:TTgg** | **0.125** | **0.375** | **0.25** | **0.75** | 0.03 | 0.97 | 0 | 0 |
| **$\Theta_4$** | **2H1** | **H1:TTgg** | **0.125** | **0.375** | **0.5** | **0.25** | 0.02 | 0.97 | 0 | 0.01 |
| **$\Theta_4$** | **2H1** | **H1:TTgg** | **0.125** | **0.375** | **0.5** | **0.5** | 0.04 | 0.96 | 0 | 0 |
| **$\Theta_4$** | **2H1** | **H1:TTgg** | **0.125** | **0.375** | **0.5** | **0.75** | 0.04 | 0.96 | 0 | 0 |
| **$\Theta_4$** | **2H1** | **H1:TTgg** | **0.125** | **0.375** | **0.75** | **0.25** | 0.03 | 0.96 | 0 | 0.01 |
| **$\Theta_4$** | **2H1** | **H1:TTgg** | **0.125** | **0.375** | **0.75** | **0.5** | 0.03 | 0.97 | 0 | 0 |
| **$\Theta_4$** | **2H1** | **H1:TTgg** | **0.125** | **0.375** | **0.75** | **0.75** | 0.02 | 0.97 | 0 | 0.01 |
| **$\Theta_4$** | **2H1** | **H1:TTgg** | **0.25** | **0.25** | **0.25** | **0.25** | 0.04 | 0.95 | 0 | 0.01 |
| **$\Theta_4$** | **2H1** | **H1:TTgg** | **0.25** | **0.25** | **0.25** | **0.5** | 0.05 | 0.95 | 0 | 0 |
| **$\Theta_4$** | **2H1** | **H1:TTgg** | **0.25** | **0.25** | **0.25** | **0.75** | 0.05 | 0.95 | 0 | 0 |
| **$\Theta_4$** | **2H1** | **H1:TTgg** | **0.25** | **0.25** | **0.5** | **0.25** | 0.07 | 0.92 | 0 | 0.01 |
| **$\Theta_4$** | **2H1** | **H1:TTgg** | **0.25** | **0.25** | **0.5** | **0.5** | 0.20 | 0.80 | 0 | 0 |
| **$\Theta_4$** | **2H1** | **H1:TTgg** | **0.25** | **0.25** | **0.5** | **0.75** | 0.14 | 0.86 | 0 | 0 |
| **$\Theta_4$** | **2H1** | **H1:TTgg** | **0.25** | **0.25** | **0.75** | **0.25** | 0.04 | 0.95 | 0 | 0.01 |
| **$\Theta_4$** | **2H1** | **H1:TTgg** | **0.25** | **0.25** | **0.75** | **0.5** | 0.14 | 0.86 | 0 | 0 |
| **$\Theta_4$** | **2H1** | **H1:TTgg** | **0.25** | **0.25** | **0.75** | **0.75** | 0.13 | 0.87 | 0 | 0 |
| **$\Theta_4$** | **2H1** | **H1:TTgg** | **0.25** | **0.75** | **0.25** | **0.25** | 0.17 | 0.82 | 0 | 0.01 |
| **$\Theta_4$** | **2H1** | **H1:TTgg** | **0.25** | **0.75** | **0.25** | **0.5** | 0.16 | 0.84 | 0 | 0 |
| **$\Theta_4$** | **2H1** | **H1:TTgg** | **0.25** | **0.75** | **0.25** | **0.75** | 0.18 | 0.82 | 0 | 0 |
| **$\Theta_4$** | **2H1** | **H1:TTgg** | **0.25** | **0.75** | **0.5** | **0.25** | 0.29 | 0.71 | 0 | 0 |
| **$\Theta_4$** | **2H1** | **H1:TTgg** | **0.25** | **0.75** | **0.5** | **0.5** | 0.31 | 0.69 | 0 | 0 |
| **$\Theta_4$** | **2H1** | **H1:TTgg** | **0.25** | **0.75** | **0.5** | **0.75** | 0.34 | 0.66 | 0 | 0 |
| **$\Theta_4$** | **2H1** | **H1:TTgg** | **0.25** | **0.75** | **0.75** | **0.25** | 0.16 | 0.84 | 0 | 0 |
| **$\Theta_4$** | **2H1** | **H1:TTgg** | **0.25** | **0.75** | **0.75** | **0.5** | 0.17 | 0.83 | 0 | 0 |
| **$\Theta_4$** | **2H1** | **H1:TTgg** | **0.25** | **0.75** | **0.75** | **0.75** | 0.16 | 0.84 | 0 | 0 |
| **$\Theta_4$** | **2H1** | **H1:TTgg** | **0.375** | **0.125** | **0.25** | **0.25** | 0.04 | 0.96 | 0 | 0 |
| **$\Theta_4$** | **2H1** | **H1:TTgg** | **0.375** | **0.125** | **0.25** | **0.5** | 0.06 | 0.94 | 0 | 0 |
| **$\Theta_4$** | **2H1** | **H1:TTgg** | **0.375** | **0.125** | **0.25** | **0.75** | 0.04 | 0.96 | 0 | 0 |
| **$\Theta_4$** | **2H1** | **H1:TTgg** | **0.375** | **0.125** | **0.5** | **0.25** | 0.07 | 0.92 | 0 | 0.01 |
| **$\Theta_4$** | **2H1** | **H1:TTgg** | **0.375** | **0.125** | **0.5** | **0.5** | 0.18 | 0.82 | 0 | 0 |
| **$\Theta_4$** | **2H1** | **H1:TTgg** | **0.375** | **0.125** | **0.5** | **0.75** | 0.20 | 0.80 | 0 | 0 |
| **$\Theta_4$** | **2H1** | **H1:TTgg** | **0.375** | **0.125** | **0.75** | **0.25** | 0.04 | 0.95 | 0 | 0.01 |
| **$\Theta_4$** | **2H1** | **H1:TTgg** | **0.375** | **0.125** | **0.75** | **0.5** | 0.15 | 0.85 | 0 | 0 |
| **$\Theta_4$** | **2H1** | **H1:TTgg** | **0.375** | **0.125** | **0.75** | **0.75** | 0.22 | 0.78 | 0 | 0 |
| **$\Theta_4$** | **2H1** | **H1:TTgg** | **0.375** | **1.125** | **0.25** | **0.25** | 0.30 | 0.70 | 0 | 0 |
| **$\Theta_4$** | **2H1** | **H1:TTgg** | **0.375** | **1.125** | **0.25** | **0.5** | 0.33 | 0.67 | 0 | 0 |
| **$\Theta_4$** | **2H1** | **H1:TTgg** | **0.375** | **1.125** | **0.25** | **0.75** | 0.30 | 0.70 | 0 | 0 |
| **$\Theta_4$** | **2H1** | **H1:TTgg** | **0.375** | **1.125** | **0.5** | **0.25** | 0.62 | 0.38 | 0 | 0 |
| **$\Theta_4$** | **2H1** | **H1:TTgg** | **0.375** | **1.125** | **0.5** | **0.5** | 0.63 | 0.37 | 0 | 0 |
| **$\Theta_4$** | **2H1** | **H1:TTgg** | **0.375** | **1.125** | **0.5** | **0.75** | 0.67 | 0.33 | 0 | 0 |
| **$\Theta_4$** | **2H1** | **H1:TTgg** | **0.375** | **1.125** | **0.75** | **0.25** | 0.31 | 0.69 | 0 | 0 |
| **$\Theta_4$** | **2H1** | **H1:TTgg** | **0.375** | **1.125** | **0.75** | **0.5** | 0.32 | 0.68 | 0 | 0 |
| **$\Theta_4$** | **2H1** | **H1:TTgg** | **0.375** | **1.125** | **0.75** | **0.75** | 0.29 | 0.71 | 0 | 0 |
| **$\Theta_4$** | **2H1** | **H1:TTgg** | **0.5** | **0.5** | **0.25** | **0.25** | 0.40 | 0.57 | 0 | 0.03 |

| | | | | | | | | | | |
|---|---|---|---|---|---|---|---|---|---|---|
| $\Theta_4$ | **2H1** | **H1:TTgg** | **0.5** | **0.5** | **0.25** | **0.5** | 0.50 | 0.50 | 0 | 0 |
| $\Theta_4$ | **2H1** | **H1:TTgg** | **0.5** | **0.5** | **0.25** | **0.75** | 0.44 | 0.56 | 0 | 0 |
| $\Theta_4$ | **2H1** | **H1:TTgg** | **0.5** | **0.5** | **0.5** | **0.25** | 0.74 | 0.25 | 0 | 0.01 |
| $\Theta_4$ | **2H1** | **H1:TTgg** | **0.5** | **0.5** | **0.5** | **0.5** | 0.92 | 0.08 | 0 | 0 |
| $\Theta_4$ | **2H1** | **H1:TTgg** | **0.5** | **0.5** | **0.5** | **0.75** | 0.90 | 0.10 | 0 | 0 |
| $\Theta_4$ | **2H1** | **H1:TTgg** | **0.5** | **0.5** | **0.75** | **0.25** | 0.54 | 0.46 | 0 | 0 |
| $\Theta_4$ | **2H1** | **H1:TTgg** | **0.5** | **0.5** | **0.75** | **0.5** | 0.59 | 0.41 | 0 | 0 |
| $\Theta_4$ | **2H1** | **H1:TTgg** | **0.5** | **0.5** | **0.75** | **0.75** | 0.60 | 0.40 | 0 | 0 |
| $\Theta_4$ | **2H1** | **H1:TTgg** | **0.75** | **0.25** | **0.25** | **0.25** | 0.21 | 0.76 | 0 | 0.03 |
| $\Theta_4$ | **2H1** | **H1:TTgg** | **0.75** | **0.25** | **0.25** | **0.5** | 0.45 | 0.54 | 0 | 0.01 |
| $\Theta_4$ | **2H1** | **H1:TTgg** | **0.75** | **0.25** | **0.25** | **0.75** | 0.44 | 0.56 | 0 | 0 |
| $\Theta_4$ | **2H1** | **H1:TTgg** | **0.75** | **0.25** | **0.5** | **0.25** | 0.37 | 0.58 | 0 | 0.05 |
| $\Theta_4$ | **2H1** | **H1:TTgg** | **0.75** | **0.25** | **0.5** | **0.5** | 0.80 | 0.19 | 0 | 0.01 |
| $\Theta_4$ | **2H1** | **H1:TTgg** | **0.75** | **0.25** | **0.5** | **0.75** | 0.86 | 0.14 | 0 | 0 |
| $\Theta_4$ | **2H1** | **H1:TTgg** | **0.75** | **0.25** | **0.75** | **0.25** | 0.49 | 0.47 | 0 | 0.04 |
| $\Theta_4$ | **2H1** | **H1:TTgg** | **0.75** | **0.25** | **0.75** | **0.5** | 0.84 | 0.16 | 0 | 0 |
| $\Theta_4$ | **2H1** | **H1:TTgg** | **0.75** | **0.25** | **0.75** | **0.75** | 0.93 | 0.07 | 0 | 0 |
| $\Theta_4$ | **2H1** | **H1:TTgg** | **0.75** | **0.75** | **0.25** | **0.25** | 0.83 | 0.16 | 0 | 0.01 |
| $\Theta_4$ | **2H1** | **H1:TTgg** | **0.75** | **0.75** | **0.25** | **0.5** | 0.88 | 0.12 | 0 | 0 |
| $\Theta_4$ | **2H1** | **H1:TTgg** | **0.75** | **0.75** | **0.25** | **0.75** | 0.85 | 0.15 | 0 | 0 |
| $\Theta_4$ | **2H1** | **H1:TTgg** | **0.75** | **0.75** | **0.5** | **0.25** | 0.98 | 0.02 | 0 | 0 |
| $\Theta_4$ | **2H1** | **H1:TTgg** | **0.75** | **0.75** | **0.5** | **0.5** | 1 | 0 | 0 | 0 |
| $\Theta_4$ | **2H1** | **H1:TTgg** | **0.75** | **0.75** | **0.5** | **0.75** | 1 | 0 | 0 | 0 |
| $\Theta_4$ | **2H1** | **H1:TTgg** | **0.75** | **0.75** | **0.75** | **0.25** | 0.86 | 0.14 | 0 | 0 |
| $\Theta_4$ | **2H1** | **H1:TTgg** | **0.75** | **0.75** | **0.75** | **0.5** | 0.87 | 0.13 | 0 | 0 |
| $\Theta_4$ | **2H1** | **H1:TTgg** | **0.75** | **0.75** | **0.75** | **0.75** | 0.88 | 0.12 | 0 | 0 |
| $\Theta_4$ | **2H1** | **H1:TTgg** | **1.125** | **0.375** | **0.25** | **0.25** | 0.48 | 0.48 | 0 | 0.04 |
| $\Theta_4$ | **2H1** | **H1:TTgg** | **1.125** | **0.375** | **0.25** | **0.5** | 0.84 | 0.16 | 0 | 0 |
| $\Theta_4$ | **2H1** | **H1:TTgg** | **1.125** | **0.375** | **0.25** | **0.75** | 0.78 | 0.22 | 0 | 0 |
| $\Theta_4$ | **2H1** | **H1:TTgg** | **1.125** | **0.375** | **0.5** | **0.25** | 0.71 | 0.25 | 0 | 0.04 |
| $\Theta_4$ | **2H1** | **H1:TTgg** | **1.125** | **0.375** | **0.5** | **0.5** | 0.99 | 0.01 | 0 | 0 |
| $\Theta_4$ | **2H1** | **H1:TTgg** | **1.125** | **0.375** | **0.5** | **0.75** | 0.97 | 0.03 | 0 | 0 |
| $\Theta_4$ | **2H1** | **H1:TTgg** | **1.125** | **0.375** | **0.75** | **0.25** | 0.82 | 0.16 | 0 | 0.02 |
| $\Theta_4$ | **2H1** | **H1:TTgg** | **1.125** | **0.375** | **0.75** | **0.5** | 0.99 | 0.01 | 0 | 0 |
| $\Theta_4$ | **2H1** | **H1:TTgg** | **1.125** | **0.375** | **0.75** | **0.75** | 1 | 0 | 0 | 0 |
| $\Theta_4$ | **2H2** | **H2:TTgg** | **0.125** | **0.375** | **0.25** | **0.25** | 0.05 | 0.95 | 0 | 0 |
| $\Theta_4$ | **2H2** | **H2:TTgg** | **0.125** | **0.375** | **0.25** | **0.5** | 0.05 | 0.95 | 0 | 0 |
| $\Theta_4$ | **2H2** | **H2:TTgg** | **0.125** | **0.375** | **0.25** | **0.75** | 0.05 | 0.94 | 0 | 0.01 |
| $\Theta_4$ | **2H2** | **H2:TTgg** | **0.125** | **0.375** | **0.5** | **0.25** | 0.05 | 0.95 | 0 | 0 |
| $\Theta_4$ | **2H2** | **H2:TTgg** | **0.125** | **0.375** | **0.5** | **0.5** | 0.06 | 0.94 | 0 | 0 |
| $\Theta_4$ | **2H2** | **H2:TTgg** | **0.125** | **0.375** | **0.5** | **0.75** | 0.07 | 0.93 | 0 | 0 |
| $\Theta_4$ | **2H2** | **H2:TTgg** | **0.25** | **0.25** | **0.25** | **0.25** | 0.07 | 0.93 | 0 | 0 |
| $\Theta_4$ | **2H2** | **H2:TTgg** | **0.25** | **0.25** | **0.25** | **0.5** | 0.09 | 0.91 | 0 | 0 |
| $\Theta_4$ | **2H2** | **H2:TTgg** | **0.25** | **0.25** | **0.25** | **0.75** | 0.09 | 0.91 | 0 | 0 |
| $\Theta_4$ | **2H2** | **H2:TTgg** | **0.25** | **0.25** | **0.5** | **0.25** | 0.21 | 0.79 | 0 | 0 |
| $\Theta_4$ | **2H2** | **H2:TTgg** | **0.25** | **0.25** | **0.5** | **0.5** | 0.18 | 0.82 | 0 | 0 |
| $\Theta_4$ | **2H2** | **H2:TTgg** | **0.25** | **0.25** | **0.5** | **0.75** | 0.20 | 0.80 | 0 | 0 |
| $\Theta_4$ | **2H2** | **H2:TTgg** | **0.25** | **0.75** | **0.25** | **0.25** | 0.16 | 0.84 | 0 | 0 |
| $\Theta_4$ | **2H2** | **H2:TTgg** | **0.25** | **0.75** | **0.25** | **0.5** | 0.17 | 0.83 | 0 | 0 |
| $\Theta_4$ | **2H2** | **H2:TTgg** | **0.25** | **0.75** | **0.25** | **0.75** | 0.16 | 0.84 | 0 | 0 |

| | | | | | | | | | | |
|---|---|---|---|---|---|---|---|---|---|---|
| **$\Theta_4$** | **2H2** | **H2:TTgg** | **0.25** | **0.75** | **0.5** | **0.25** | 0.30 | 0.70 | 0 | 0 |
| **$\Theta_4$** | **2H2** | **H2:TTgg** | **0.25** | **0.75** | **0.5** | **0.5** | 0.32 | 0.68 | 0 | 0 |
| **$\Theta_4$** | **2H2** | **H2:TTgg** | **0.25** | **0.75** | **0.5** | **0.75** | 0.30 | 0.70 | 0 | 0 |
| **$\Theta_4$** | **2H2** | **H2:TTgg** | **0.375** | **0.125** | **0.25** | **0.25** | 0.03 | 0.97 | 0 | 0 |
| **$\Theta_4$** | **2H2** | **H2:TTgg** | **0.375** | **0.125** | **0.25** | **0.5** | 0.05 | 0.94 | 0 | 0.01 |
| **$\Theta_4$** | **2H2** | **H2:TTgg** | **0.375** | **0.125** | **0.25** | **0.75** | 0.06 | 0.93 | 0 | 0.01 |
| **$\Theta_4$** | **2H2** | **H2:TTgg** | **0.375** | **0.125** | **0.5** | **0.25** | 0.14 | 0.85 | 0 | 0.01 |
| **$\Theta_4$** | **2H2** | **H2:TTgg** | **0.375** | **0.125** | **0.5** | **0.5** | 0.14 | 0.86 | 0 | 0 |
| **$\Theta_4$** | **2H2** | **H2:TTgg** | **0.375** | **0.125** | **0.5** | **0.75** | 0.14 | 0.85 | 0 | 0.01 |
| **$\Theta_4$** | **2H2** | **H2:TTgg** | **0.375** | **1.125** | **0.25** | **0.25** | 0.29 | 0.71 | 0 | 0 |
| **$\Theta_4$** | **2H2** | **H2:TTgg** | **0.375** | **1.125** | **0.25** | **0.5** | 0.29 | 0.71 | 0 | 0 |
| **$\Theta_4$** | **2H2** | **H2:TTgg** | **0.375** | **1.125** | **0.25** | **0.75** | 0.29 | 0.71 | 0 | 0 |
| **$\Theta_4$** | **2H2** | **H2:TTgg** | **0.375** | **1.125** | **0.5** | **0.25** | 0.64 | 0.36 | 0 | 0 |
| **$\Theta_4$** | **2H2** | **H2:TTgg** | **0.375** | **1.125** | **0.5** | **0.5** | 0.62 | 0.38 | 0 | 0 |
| **$\Theta_4$** | **2H2** | **H2:TTgg** | **0.375** | **1.125** | **0.5** | **0.75** | 0.60 | 0.40 | 0 | 0 |
| **$\Theta_4$** | **2H2** | **H2:TTgg** | **0.5** | **0.5** | **0.25** | **0.25** | 0.58 | 0.42 | 0 | 0 |
| **$\Theta_4$** | **2H2** | **H2:TTgg** | **0.5** | **0.5** | **0.25** | **0.5** | 0.57 | 0.43 | 0 | 0 |
| **$\Theta_4$** | **2H2** | **H2:TTgg** | **0.5** | **0.5** | **0.25** | **0.75** | 0.55 | 0.45 | 0 | 0 |
| **$\Theta_4$** | **2H2** | **H2:TTgg** | **0.5** | **0.5** | **0.5** | **0.25** | 0.94 | 0.06 | 0 | 0 |
| **$\Theta_4$** | **2H2** | **H2:TTgg** | **0.5** | **0.5** | **0.5** | **0.5** | 0.94 | 0.06 | 0 | 0 |
| **$\Theta_4$** | **2H2** | **H2:TTgg** | **0.5** | **0.5** | **0.5** | **0.75** | 0.93 | 0.07 | 0 | 0 |
| **$\Theta_4$** | **2H2** | **H2:TTgg** | **0.75** | **0.25** | **0.25** | **0.25** | 0.68 | 0.32 | 0 | 0 |
| **$\Theta_4$** | **2H2** | **H2:TTgg** | **0.75** | **0.25** | **0.25** | **0.5** | 0.61 | 0.39 | 0 | 0 |
| **$\Theta_4$** | **2H2** | **H2:TTgg** | **0.75** | **0.25** | **0.25** | **0.75** | 0.52 | 0.47 | 0 | 0.01 |
| **$\Theta_4$** | **2H2** | **H2:TTgg** | **0.75** | **0.25** | **0.5** | **0.25** | 0.68 | 0.32 | 0 | 0 |
| **$\Theta_4$** | **2H2** | **H2:TTgg** | **0.75** | **0.25** | **0.5** | **0.5** | 0.68 | 0.32 | 0 | 0 |
| **$\Theta_4$** | **2H2** | **H2:TTgg** | **0.75** | **0.25** | **0.5** | **0.75** | 0.66 | 0.33 | 0 | 0.01 |
| **$\Theta_4$** | **2H2** | **H2:TTgg** | **0.75** | **0.75** | **0.25** | **0.25** | 0.87 | 0.13 | 0 | 0 |
| **$\Theta_4$** | **2H2** | **H2:TTgg** | **0.75** | **0.75** | **0.25** | **0.5** | 0.87 | 0.13 | 0 | 0 |
| **$\Theta_4$** | **2H2** | **H2:TTgg** | **0.75** | **0.75** | **0.25** | **0.75** | 0.82 | 0.18 | 0 | 0 |
| **$\Theta_4$** | **2H2** | **H2:TTgg** | **0.75** | **0.75** | **0.5** | **0.25** | 1 | 0 | 0 | 0 |
| **$\Theta_4$** | **2H2** | **H2:TTgg** | **0.75** | **0.75** | **0.5** | **0.5** | 1 | 0 | 0 | 0 |
| **$\Theta_4$** | **2H2** | **H2:TTgg** | **0.75** | **0.75** | **0.5** | **0.75** | 0.94 | 0.06 | 0 | 0 |
| **$\Theta_4$** | **2H2** | **H2:TTgg** | **1.125** | **0.375** | **0.25** | **0.25** | 0.91 | 0.09 | 0 | 0 |
| **$\Theta_4$** | **2H2** | **H2:TTgg** | **1.125** | **0.375** | **0.25** | **0.5** | 0.89 | 0.11 | 0 | 0 |
| **$\Theta_4$** | **2H2** | **H2:TTgg** | **1.125** | **0.375** | **0.25** | **0.75** | 0.83 | 0.16 | 0 | 0.01 |
| **$\Theta_4$** | **2H2** | **H2:TTgg** | **1.125** | **0.375** | **0.5** | **0.25** | 0.87 | 0.13 | 0 | 0 |
| **$\Theta_4$** | **2H2** | **H2:TTgg** | **1.125** | **0.375** | **0.5** | **0.5** | 0.87 | 0.13 | 0 | 0 |
| **$\Theta_4$** | **2H2** | **H2:TTgg** | **1.125** | **0.375** | **0.5** | **0.75** | 0.85 | 0.15 | 0 | 0 |

### Table S5.

**Testing the *reverse* method and empirical distribution criterion (eCDF) for frequency of correct recognition of models in the Poisson randomized set.** The first column indicates group *Θj*, the following two columns show the trivial name of the model and its alias in TTgg notation, and the following four columns describe the initial values of parameters T1, T3, $\gamma_1$, $\gamma_3$ taken in a Poisson-distribution based randomization process. The last four columns present the frequency of the correct model recognition: $\boldsymbol{\Theta_j^i}$; frequency of error recognition of the model in the direction of a simple model:*Θj-*; frequency of error recognition of the model to a complex model: *Θj+*; frequency of error recognition of the model because optimization process selects another model from the same group: *Θj*.

| Group | Model | Alias | $T_1$ | $T_3$ | $\gamma_1$ | $\gamma_3$ | $\Theta_j^i$ | $\Theta_{j-}$ | $\Theta_{j+}$ | $\Theta_j$ |
|---|---|---|---|---|---|---|---|---|---|---|
| $\Theta_0$ | P | H1:00nn | 0 | 0 | - | - | 0.98 | 0 | 0.02 | 0 |
| $\Theta_1$ | TP | H1:0Tn1 | 0 | 0.5 | - | 1 | 0.98 | 0 | 0.02 | 0 |
| $\Theta_1$ | TP | H1:0Tn1 | 0 | 1 | - | 1 | 0.98 | 0 | 0.02 | 0 |
| $\Theta_1$ | TP | H1:0Tn1 | 0 | 1.5 | - | 1 | 0.97 | 0 | 0.03 | 0 |
| $\Theta_1$ | PT | H1:T00n | 0.5 | 0 | 0 | - | 0.97 | 0 | 0.03 | 0 |
| $\Theta_1$ | PT | H1:T00n | 1 | 0 | 0 | - | 0.97 | 0 | 0.03 | 0 |
| $\Theta_1$ | PT | H1:T00n | 1.5 | 0 | 0 | - | 0.97 | 0 | 0.03 | 0 |
| $\Theta_2$ | 1HP1 | H1:0Tng | 0 | 0.5 | - | 0.25 | 0.88 | 0.05 | 0.05 | 0.02 |
| $\Theta_2$ | 1HP1 | H1:0Tng | 0 | 0.5 | - | 0.5 | 0.95 | 0.02 | 0.02 | 0.01 |
| $\Theta_2$ | 1HP1 | H1:0Tng | 0 | 1 | - | 0.25 | 0.98 | 0 | 0.02 | 0 |
| $\Theta_2$ | 1HP1 | H1:0Tng | 0 | 1 | - | 0.5 | 0.98 | 0 | 0.02 | 0 |
| $\Theta_2$ | 1HP1 | H1:0Tng | 0 | 1.5 | - | 0.25 | 0.98 | 0 | 0.02 | 0 |
| $\Theta_2$ | 1HP1 | H1:0Tng | 0 | 1.5 | - | 0.5 | 0.97 | 0 | 0.03 | 0 |
| $\Theta_2$ | 1HP2 | H1:T0g0 | 0.5 | 0 | 0.25 | 0 | 0.82 | 0.12 | 0.05 | 0.01 |
| $\Theta_2$ | 1HP2 | H1:T0g0 | 0.5 | 0 | 0.5 | 0 | 0.97 | 0.01 | 0.02 | 0 |
| $\Theta_2$ | 1HP2 | H1:T0g0 | 0.5 | 0 | 0.75 | 0 | 0.90 | 0 | 0.05 | 0.05 |
| $\Theta_2$ | 1HP2 | H1:T0g0 | 1 | 0 | 0.25 | 0 | 0.97 | 0 | 0.03 | 0 |
| $\Theta_2$ | 1HP2 | H1:T0g0 | 1 | 0 | 0.5 | 0 | 0.97 | 0 | 0.03 | 0 |
| $\Theta_2$ | 1HP2 | H1:T0g0 | 1 | 0 | 0.75 | 0 | 0.98 | 0 | 0.02 | 0 |
| $\Theta_2$ | 1HP2 | H1:T0g0 | 1.5 | 0 | 0.25 | 0 | 0.97 | 0 | 0.03 | 0 |
| $\Theta_2$ | 1HP2 | H1:T0g0 | 1.5 | 0 | 0.5 | 0 | 0.97 | 0 | 0.03 | 0 |
| $\Theta_2$ | 1HP2 | H1:T0g0 | 1.5 | 0 | 0.75 | 0 | 0.97 | 0 | 0.03 | 0 |
| $\Theta_2$ | 1HP3 | H1:T0g1 | 0.5 | 0 | 0.5 | 1 | 0.86 | 0.04 | 0.04 | 0.06 |
| $\Theta_2$ | 1HP3 | H1:T0g1 | 0.5 | 0 | 0.25 | 1 | 0.94 | 0.02 | 0.03 | 0.01 |
| $\Theta_2$ | 1HP3 | H1:T0g1 | 1 | 0 | 0.5 | 1 | 0.98 | 0 | 0.02 | 0 |
| $\Theta_2$ | 1HP3 | H1:T0g1 | 1 | 0 | 0.25 | 1 | 0.97 | 0 | 0.03 | 0 |
| $\Theta_2$ | 1HP3 | H1:T0g1 | 1.5 | 0 | 0.5 | 1 | 0.97 | 0 | 0.03 | 0 |
| $\Theta_2$ | 1HP3 | H1:T0g1 | 1.5 | 0 | 0.25 | 1 | 0.97 | 0 | 0.03 | 0 |
| $\Theta_2$ | T1 | H1:TT10 | 0.125 | 0.375 | 1 | 0 | 0.77 | 0.11 | 0.07 | 0.05 |
| $\Theta_2$ | T1 | H1:TT10 | 0.25 | 0.25 | 1 | 0 | 0.84 | 0 | 0 | 0.16 |
| $\Theta_2$ | T1 | H1:TT10 | 0.25 | 0.75 | 1 | 0 | 0.92 | 0.04 | 0.04 | 0 |
| $\Theta_2$ | T1 | H1:TT10 | 0.375 | 0.125 | 1 | 0 | 0.65 | 0 | 0 | 0.35 |
| $\Theta_2$ | T1 | H1:TT10 | 0.375 | 1.125 | 1 | 0 | 0.97 | 0 | 0.03 | 0 |
| $\Theta_2$ | T1 | H1:TT10 | 0.5 | 0.5 | 1 | 0 | 0.96 | 0 | 0.03 | 0.01 |
| $\Theta_2$ | T1 | H1:TT10 | 0.75 | 0.25 | 1 | 0 | 0.73 | 0 | 0 | 0.27 |
| $\Theta_2$ | T1 | H1:TT10 | 0.75 | 0.75 | 1 | 0 | 0.96 | 0 | 0.04 | 0 |
| $\Theta_2$ | T1 | H1:TT10 | 1.125 | 0.375 | 1 | 0 | 0.72 | 0 | 0.10 | 0.18 |
| $\Theta_2$ | T2 | H1:TT01 | 0.125 | 0.375 | 0 | 1 | 0.62 | 0.16 | 0.14 | 0.08 |

| | | | | | | | | | | |
|---|---|---|---|---|---|---|---|---|---|---|
| $\Theta_2$ | T2 | H1:TT01 | 0.25 | 0.25 | 0 | 1 | 0.89 | 0.04 | 0.01 | 0.06 |
| $\Theta_2$ | T2 | H1:TT01 | 0.25 | 0.75 | 0 | 1 | 0.93 | 0.03 | 0.04 | 0 |
| $\Theta_2$ | T2 | H1:TT01 | 0.375 | 0.125 | 0 | 1 | 0.55 | 0.21 | 0.07 | 0.17 |
| $\Theta_2$ | T2 | H1:TT01 | 0.375 | 1.125 | 0 | 1 | 0.97 | 0 | 0.03 | 0 |
| $\Theta_2$ | T2 | H1:TT01 | 0.5 | 0.5 | 0 | 1 | 0.97 | 0 | 0.03 | 0 |
| $\Theta_2$ | T2 | H1:TT01 | 0.75 | 0.25 | 0 | 1 | 0.90 | 0.05 | 0.02 | 0.03 |
| $\Theta_2$ | T2 | H1:TT01 | 0.75 | 0.75 | 0 | 1 | 0.97 | 0 | 0.03 | 0 |
| $\Theta_3$ | T2 | H1:TT01 | 1.125 | 0.375 | 0 | 0 | 0.97 | 0 | 0.03 | 0 |
| $\Theta_3$ | 1H1 | H1:TTg0 | 0.125 | 0.375 | 0.25 | 0 | 0.20 | 0.48 | 0.30 | 0.02 |
| $\Theta_3$ | 1H1 | H1:TTg0 | 0.125 | 0.375 | 0.5 | 0 | 0.21 | 0.48 | 0.30 | 0.01 |
| $\Theta_3$ | 1H1 | H1:TTg0 | 0.125 | 0.375 | 0.75 | 0 | 0.20 | 0.49 | 0.27 | 0.04 |
| $\Theta_3$ | 1H1 | H1:TTg0 | 0.25 | 0.25 | 0.25 | 0 | 0.22 | 0.47 | 0.28 | 0.03 |
| $\Theta_3$ | 1H1 | H1:TTg0 | 0.25 | 0.25 | 0.5 | 0 | 0.29 | 0.31 | 0.36 | 0.04 |
| $\Theta_3$ | 1H1 | H1:TTg0 | 0.25 | 0.25 | 0.75 | 0 | 0.23 | 0.31 | 0.38 | 0.08 |
| $\Theta_3$ | 1H1 | H1:TTg0 | 0.25 | 0.75 | 0.25 | 0 | 0.29 | 0.30 | 0.38 | 0.03 |
| $\Theta_3$ | 1H1 | H1:TTg0 | 0.25 | 0.75 | 0.5 | 0 | 0.38 | 0.28 | 0.32 | 0.02 |
| $\Theta_3$ | 1H1 | H1:TTg0 | 0.25 | 0.75 | 0.75 | 0 | 0.27 | 0.30 | 0.36 | 0.07 |
| $\Theta_3$ | 1H1 | H1:TTg0 | 0.375 | 0.125 | 0.25 | 0 | 0.19 | 0.32 | 0.45 | 0.04 |
| $\Theta_3$ | 1H1 | H1:TTg0 | 0.375 | 0.125 | 0.5 | 0 | 0.25 | 0.31 | 0.41 | 0.03 |
| $\Theta_3$ | 1H1 | H1:TTg0 | 0.375 | 0.125 | 0.75 | 0 | 0.21 | 0.32 | 0.41 | 0.06 |
| $\Theta_3$ | 1H1 | H1:TTg0 | 0.375 | 1.125 | 0.25 | 0 | 0.38 | 0.27 | 0.31 | 0.04 |
| $\Theta_3$ | 1H1 | H1:TTg0 | 0.375 | 1.125 | 0.5 | 0 | 0.66 | 0.15 | 0.17 | 0.02 |
| $\Theta_3$ | 1H1 | H1:TTg0 | 0.375 | 1.125 | 0.75 | 0 | 0.36 | 0.28 | 0.30 | 0.06 |
| $\Theta_3$ | 1H1 | H1:TTg0 | 0.5 | 0.5 | 0.25 | 0 | 0.62 | 0.17 | 0.20 | 0.01 |
| $\Theta_3$ | 1H1 | H1:TTg0 | 0.5 | 0.5 | 0.5 | 0 | 0.88 | 0.06 | 0.06 | 0 |
| $\Theta_3$ | 1H1 | H1:TTg0 | 0.5 | 0.5 | 0.75 | 0 | 0.60 | 0.17 | 0.17 | 0.06 |
| $\Theta_3$ | 1H1 | H1:TTg0 | 0.75 | 0.25 | 0.25 | 0 | 0.65 | 0.17 | 0.17 | 0.01 |
| $\Theta_3$ | 1H1 | H1:TTg0 | 0.75 | 0.25 | 0.5 | 0 | 0.63 | 0.16 | 0.21 | 0 |
| $\Theta_3$ | 1H1 | H1:TTg0 | 0.75 | 0.25 | 0.75 | 0 | 0.46 | 0.26 | 0.26 | 0.02 |
| $\Theta_3$ | 1H1 | H1:TTg0 | 0.75 | 0.75 | 0.25 | 0 | 0.85 | 0.07 | 0.07 | 0.01 |
| $\Theta_3$ | 1H1 | H1:TTg0 | 0.75 | 0.75 | 0.5 | 0 | 0.96 | 0.01 | 0.03 | 0 |
| $\Theta_3$ | 1H1 | H1:TTg0 | 0.75 | 0.75 | 0.75 | 0 | 0.81 | 0.09 | 0.08 | 0.02 |
| $\Theta_3$ | 1H1 | H1:TTg0 | 1.125 | 0.375 | 0.25 | 0 | 0.87 | 0.06 | 0.07 | 0 |
| $\Theta_3$ | 1H1 | H1:TTg0 | 1.125 | 0.375 | 0.5 | 0 | 0.77 | 0.11 | 0.12 | 0 |
| $\Theta_3$ | 1H1 | H1:TTg0 | 1.125 | 0.375 | 0.75 | 0 | 0.67 | 0.15 | 0.17 | 0.01 |
| $\Theta_3$ | 1H2 | H1:TT1g | 0.125 | 0.375 | 1 | 0.25 | 0.44 | 0.27 | 0.28 | 0.01 |
| $\Theta_3$ | 1H2 | H1:TT1g | 0.125 | 0.375 | 1 | 0.5 | 0.62 | 0.18 | 0.20 | 0 |
| $\Theta_3$ | 1H2 | H1:TT1g | 0.125 | 0.375 | 1 | 0.75 | 0.54 | 0.2 | 0.24 | 0.02 |
| $\Theta_3$ | 1H2 | H1:TT1g | 0.25 | 0.25 | 1 | 0.25 | 0.59 | 0.19 | 0.18 | 0.04 |
| $\Theta_3$ | 1H2 | H1:TT1g | 0.25 | 0.25 | 1 | 0.5 | 0.83 | 0.08 | 0.05 | 0.04 |
| $\Theta_3$ | 1H2 | H1:TT1g | 0.25 | 0.25 | 1 | 0.75 | 0.70 | 0.13 | 0.14 | 0.03 |
| $\Theta_3$ | 1H2 | H1:TT1g | 0.25 | 0.75 | 1 | 0.25 | 0.95 | 0.03 | 0.02 | 0 |
| $\Theta_3$ | 1H2 | H1:TT1g | 0.25 | 0.75 | 1 | 0.5 | 0.94 | 0.03 | 0.03 | 0 |
| $\Theta_3$ | 1H2 | H1:TT1g | 0.25 | 0.75 | 1 | 0.75 | 0.94 | 0.03 | 0.03 | 0 |
| $\Theta_3$ | 1H2 | H1:TT1g | 0.375 | 0.125 | 1 | 0.25 | 0.44 | 0.18 | 0.32 | 0.06 |
| $\Theta_3$ | 1H2 | H1:TT1g | 0.375 | 0.125 | 1 | 0.5 | 0.49 | 0.17 | 0.27 | 0.07 |
| $\Theta_3$ | 1H2 | H1:TT1g | 0.375 | 0.125 | 1 | 0.75 | 0.51 | 0.17 | 0.27 | 0.05 |
| $\Theta_3$ | 1H2 | H1:TT1g | 0.375 | 1.125 | 1 | 0.25 | 0.98 | 0 | 0.02 | 0 |
| $\Theta_3$ | 1H2 | H1:TT1g | 0.375 | 1.125 | 1 | 0.5 | 0.97 | 0 | 0.03 | 0 |
| $\Theta_3$ | 1H2 | H1:TT1g | 0.375 | 1.125 | 1 | 0.75 | 0.98 | 0 | 0.02 | 0 |

| | | | | | | | | | | |
|---|---|---|---|---|---|---|---|---|---|---|
| $\Theta_3$ | 1H2 | H1:TT1g | 0.5 | 0.5 | 1 | 0.25 | 0.96 | 0.02 | 0.02 | 0 |
| $\Theta_3$ | 1H2 | H1:TT1g | 0.5 | 0.5 | 1 | 0.5 | 0.97 | 0 | 0.03 | 0 |
| $\Theta_3$ | 1H2 | H1:TT1g | 0.5 | 0.5 | 1 | 0.75 | 0.97 | 0 | 0.03 | 0 |
| $\Theta_3$ | 1H2 | H1:TT1g | 0.75 | 0.25 | 1 | 0.25 | 0.68 | 0.12 | 0.13 | 0.07 |
| $\Theta_3$ | 1H2 | H1:TT1g | 0.75 | 0.25 | 1 | 0.5 | 0.79 | 0.09 | 0.08 | 0.04 |
| $\Theta_3$ | 1H2 | H1:TT1g | 0.75 | 0.25 | 1 | 0.75 | 0.87 | 0.06 | 0.02 | 0.05 |
| $\Theta_3$ | 1H2 | H1:TT1g | 0.75 | 0.75 | 1 | 0.25 | 0.98 | 0 | 0.02 | 0 |
| $\Theta_3$ | 1H2 | H1:TT1g | 0.75 | 0.75 | 1 | 0.5 | 0.97 | 0 | 0.03 | 0 |
| $\Theta_3$ | 1H2 | H1:TT1g | 0.75 | 0.75 | 1 | 0.75 | 0.98 | 0 | 0.02 | 0 |
| $\Theta_3$ | 1H2 | H1:TT1g | 1.125 | 0.375 | 1 | 0.25 | 0.82 | 0.08 | 0.08 | 0.02 |
| $\Theta_3$ | 1H2 | H1:TT1g | 1.125 | 0.375 | 1 | 0.5 | 0.92 | 0.03 | 0.04 | 0.01 |
| $\Theta_3$ | 1H2 | H1:TT1g | 1.125 | 0.375 | 1 | 0.75 | 0.96 | 0.01 | 0.02 | 0.01 |
| $\Theta_3$ | 1H3 | H1:TT0g | 0.125 | 0.375 | 0 | 0.25 | 0.41 | 0.18 | 0.33 | 0.08 |
| $\Theta_3$ | 1H3 | H1:TT0g | 0.125 | 0.375 | 0 | 0.5 | 0.53 | 0.17 | 0.24 | 0.06 |
| $\Theta_3$ | 1H3 | H1:TT0g | 0.25 | 0.25 | 0 | 0.25 | 0.39 | 0.19 | 0.34 | 0.08 |
| $\Theta_3$ | 1H3 | H1:TT0g | 0.25 | 0.25 | 0 | 0.5 | 0.47 | 0.17 | 0.28 | 0.08 |
| $\Theta_3$ | 1H3 | H1:TT0g | 0.25 | 0.75 | 0 | 0.25 | 0.90 | 0.05 | 0.03 | 0.02 |
| $\Theta_3$ | 1H3 | H1:TT0g | 0.25 | 0.75 | 0 | 0.5 | 0.95 | 0.02 | 0.03 | 0 |
| $\Theta_3$ | 1H3 | H1:TT0g | 0.375 | 0.125 | 0 | 0.25 | 0.34 | 0.19 | 0.42 | 0.05 |
| $\Theta_3$ | 1H3 | H1:TT0g | 0.375 | 0.125 | 0 | 0.5 | 0.33 | 0.19 | 0.44 | 0.04 |
| $\Theta_3$ | 1H3 | H1:TT0g | 0.375 | 1.125 | 0 | 0.25 | 0.97 | 0 | 0.03 | 0 |
| $\Theta_3$ | 1H3 | H1:TT0g | 0.375 | 1.125 | 0 | 0.5 | 0.97 | 0 | 0.03 | 0 |
| $\Theta_3$ | 1H3 | H1:TT0g | 0.5 | 0.5 | 0 | 0.25 | 0.68 | 0.13 | 0.12 | 0.07 |
| $\Theta_3$ | 1H3 | H1:TT0g | 0.5 | 0.5 | 0 | 0.5 | 0.89 | 0.05 | 0.03 | 0.03 |
| $\Theta_3$ | 1H3 | H1:TT0g | 0.75 | 0.25 | 0 | 0.25 | 0.39 | 0.19 | 0.34 | 0.08 |
| $\Theta_3$ | 1H3 | H1:TT0g | 0.75 | 0.25 | 0 | 0.5 | 0.47 | 0.17 | 0.30 | 0.06 |
| $\Theta_3$ | 1H3 | H1:TT0g | 0.75 | 0.75 | 0 | 0.25 | 0.88 | 0.06 | 0.02 | 0.04 |
| $\Theta_3$ | 1H3 | H1:TT0g | 0.75 | 0.75 | 0 | 0.5 | 0.98 | 0 | 0.02 | 0 |
| $\Theta_3$ | 1H3 | H1:TT0g | 1.125 | 0.375 | 0 | 0.25 | 0.48 | 0.16 | 0.28 | 0.08 |
| $\Theta_3$ | 1H3 | H1:TT0g | 1.125 | 0.375 | 0 | 0.5 | 0.67 | 0.13 | 0.14 | 0.06 |
| $\Theta_3$ | 1H4 | H1:TTg1 | 0.125 | 0.375 | 0.25 | 1 | 0.34 | 0.19 | 0.44 | 0.03 |
| $\Theta_3$ | 1H4 | H1:TTg1 | 0.125 | 0.375 | 0.5 | 1 | 0.35 | 0.19 | 0.44 | 0.02 |
| $\Theta_3$ | 1H4 | H1:TTg1 | 0.25 | 0.25 | 0.25 | 1 | 0.42 | 0.18 | 0.31 | 0.09 |
| $\Theta_3$ | 1H4 | H1:TTg1 | 0.25 | 0.25 | 0.5 | 1 | 0.56 | 0.16 | 0.23 | 0.05 |
| $\Theta_3$ | 1H4 | H1:TTg1 | 0.25 | 0.75 | 0.25 | 1 | 0.40 | 0.18 | 0.31 | 0.11 |
| $\Theta_3$ | 1H4 | H1:TTg1 | 0.25 | 0.75 | 0.5 | 1 | 0.49 | 0.18 | 0.29 | 0.04 |
| $\Theta_3$ | 1H4 | H1:TTg1 | 0.375 | 0.125 | 0.25 | 1 | 0.51 | 0.17 | 0.26 | 0.06 |
| $\Theta_3$ | 1H4 | H1:TTg1 | 0.375 | 0.125 | 0.5 | 1 | 0.72 | 0.12 | 0.14 | 0.02 |
| $\Theta_3$ | 1H4 | H1:TTg1 | 0.375 | 1.125 | 0.25 | 1 | 0.48 | 0.17 | 0.22 | 0.13 |
| $\Theta_3$ | 1H4 | H1:TTg1 | 0.375 | 1.125 | 0.5 | 1 | 0.69 | 0.13 | 0.14 | 0.04 |
| $\Theta_3$ | 1H4 | H1:TTg1 | 0.5 | 0.5 | 0.25 | 1 | 0.70 | 0.11 | 0.10 | 0.09 |
| $\Theta_3$ | 1H4 | H1:TTg1 | 0.5 | 0.5 | 0.5 | 1 | 0.93 | 0.02 | 0.04 | 0.01 |
| $\Theta_3$ | 1H4 | H1:TTg1 | 0.75 | 0.25 | 0.25 | 1 | 0.91 | 0.04 | 0.03 | 0.02 |
| $\Theta_3$ | 1H4 | H1:TTg1 | 0.75 | 0.25 | 0.5 | 1 | 0.97 | 0 | 0.03 | 0 |
| $\Theta_3$ | 1H4 | H1:TTg1 | 0.75 | 0.75 | 0.25 | 1 | 0.88 | 0.05 | 0.04 | 0.03 |
| $\Theta_3$ | 1H4 | H1:TTg1 | 0.75 | 0.75 | 0.5 | 1 | 0.96 | 0 | 0.04 | 0 |
| $\Theta_3$ | 1H4 | H1:TTg1 | 1.125 | 0.375 | 0.25 | 1 | 0.98 | 0 | 0.02 | 0 |
| $\Theta_3$ | 1H4 | H1:TTg1 | 1.125 | 0.375 | 0.5 | 1 | 0.97 | 0 | 0.03 | 0 |
| $\Theta_3$ | 2PH | H2:T0gg | 0.5 | 0 | 0.25 | 0.5 | 0.67 | 0.14 | 0.12 | 0.07 |
| $\Theta_3$ | 2PH | H2:T0gg | 0.5 | 0 | 0.25 | 0.75 | 0.59 | 0.15 | 0.16 | 0.1 |

| | | | | | | | | | | |
|---|---|---|---|---|---|---|---|---|---|---|
| $\Theta_3$ | 2PH | H2:T0gg | 1 | 0 | 0.25 | 0.5 | 0.95 | 0.01 | 0.04 | 0 |
| $\Theta_3$ | 2PH | H2:T0gg | 1 | 0 | 0.25 | 0.75 | 0.94 | 0.01 | 0.05 | 0 |
| $\Theta_3$ | 2PH | H2:T0gg | 1.5 | 0 | 0.25 | 0.5 | 0.96 | 0 | 0.04 | 0 |
| $\Theta_3$ | 2PH | H2:T0gg | 1.5 | 0 | 0.25 | 0.75 | 0.97 | 0 | 0.03 | 0 |
| $\Theta_4$ | 2H1 | H1:TTgg | 0.125 | 0.375 | 0.25 | 0.25 | 0.51 | 0.48 | 0 | 0.01 |
| $\Theta_4$ | 2H1 | H1:TTgg | 0.125 | 0.375 | 0.25 | 0.5 | 0.52 | 0.48 | 0 | 0 |
| $\Theta_4$ | 2H1 | H1:TTgg | 0.125 | 0.375 | 0.25 | 0.75 | 0.52 | 0.48 | 0 | 0 |
| $\Theta_4$ | 2H1 | H1:TTgg | 0.125 | 0.375 | 0.5 | 0.25 | 0.51 | 0.48 | 0 | 0.01 |
| $\Theta_4$ | 2H1 | H1:TTgg | 0.125 | 0.375 | 0.5 | 0.5 | 0.52 | 0.48 | 0 | 0 |
| $\Theta_4$ | 2H1 | H1:TTgg | 0.125 | 0.375 | 0.5 | 0.75 | 0.52 | 0.48 | 0 | 0 |
| $\Theta_4$ | 2H1 | H1:TTgg | 0.125 | 0.375 | 0.75 | 0.25 | 0.52 | 0.47 | 0 | 0.01 |
| $\Theta_4$ | 2H1 | H1:TTgg | 0.125 | 0.375 | 0.75 | 0.5 | 0.52 | 0.48 | 0 | 0 |
| $\Theta_4$ | 2H1 | H1:TTgg | 0.125 | 0.375 | 0.75 | 0.75 | 0.51 | 0.48 | 0 | 0.01 |
| $\Theta_4$ | 2H1 | H1:TTgg | 0.25 | 0.25 | 0.25 | 0.25 | 0.52 | 0.47 | 0 | 0.01 |
| $\Theta_4$ | 2H1 | H1:TTgg | 0.25 | 0.25 | 0.25 | 0.5 | 0.53 | 0.47 | 0 | 0 |
| $\Theta_4$ | 2H1 | H1:TTgg | 0.25 | 0.25 | 0.25 | 0.75 | 0.52 | 0.48 | 0 | 0 |
| $\Theta_4$ | 2H1 | H1:TTgg | 0.25 | 0.25 | 0.5 | 0.25 | 0.54 | 0.45 | 0 | 0.01 |
| $\Theta_4$ | 2H1 | H1:TTgg | 0.25 | 0.25 | 0.5 | 0.5 | 0.60 | 0.40 | 0 | 0 |
| $\Theta_4$ | 2H1 | H1:TTgg | 0.25 | 0.25 | 0.5 | 0.75 | 0.57 | 0.43 | 0 | 0 |
| $\Theta_4$ | 2H1 | H1:TTgg | 0.25 | 0.25 | 0.75 | 0.25 | 0.52 | 0.47 | 0 | 0.01 |
| $\Theta_4$ | 2H1 | H1:TTgg | 0.25 | 0.25 | 0.75 | 0.5 | 0.57 | 0.43 | 0 | 0 |
| $\Theta_4$ | 2H1 | H1:TTgg | 0.25 | 0.25 | 0.75 | 0.75 | 0.56 | 0.44 | 0 | 0 |
| $\Theta_4$ | 2H1 | H1:TTgg | 0.25 | 0.75 | 0.25 | 0.25 | 0.58 | 0.41 | 0 | 0.01 |
| $\Theta_4$ | 2H1 | H1:TTgg | 0.25 | 0.75 | 0.25 | 0.5 | 0.58 | 0.42 | 0 | 0 |
| $\Theta_4$ | 2H1 | H1:TTgg | 0.25 | 0.75 | 0.25 | 0.75 | 0.59 | 0.41 | 0 | 0 |
| $\Theta_4$ | 2H1 | H1:TTgg | 0.25 | 0.75 | 0.5 | 0.25 | 0.65 | 0.35 | 0 | 0 |
| $\Theta_4$ | 2H1 | H1:TTgg | 0.25 | 0.75 | 0.5 | 0.5 | 0.65 | 0.35 | 0 | 0 |
| $\Theta_4$ | 2H1 | H1:TTgg | 0.25 | 0.75 | 0.5 | 0.75 | 0.67 | 0.33 | 0 | 0 |
| $\Theta_4$ | 2H1 | H1:TTgg | 0.25 | 0.75 | 0.75 | 0.25 | 0.58 | 0.42 | 0 | 0 |
| $\Theta_4$ | 2H1 | H1:TTgg | 0.25 | 0.75 | 0.75 | 0.5 | 0.58 | 0.42 | 0 | 0 |
| $\Theta_4$ | 2H1 | H1:TTgg | 0.25 | 0.75 | 0.75 | 0.75 | 0.58 | 0.42 | 0 | 0 |
| $\Theta_4$ | 2H1 | H1:TTgg | 0.375 | 0.125 | 0.25 | 0.25 | 0.52 | 0.48 | 0 | 0 |
| $\Theta_4$ | 2H1 | H1:TTgg | 0.375 | 0.125 | 0.25 | 0.5 | 0.53 | 0.47 | 0 | 0 |
| $\Theta_4$ | 2H1 | H1:TTgg | 0.375 | 0.125 | 0.25 | 0.75 | 0.52 | 0.48 | 0 | 0 |
| $\Theta_4$ | 2H1 | H1:TTgg | 0.375 | 0.125 | 0.5 | 0.25 | 0.54 | 0.45 | 0 | 0.01 |
| $\Theta_4$ | 2H1 | H1:TTgg | 0.375 | 0.125 | 0.5 | 0.5 | 0.59 | 0.41 | 0 | 0 |
| $\Theta_4$ | 2H1 | H1:TTgg | 0.375 | 0.125 | 0.5 | 0.75 | 0.60 | 0.40 | 0 | 0 |
| $\Theta_4$ | 2H1 | H1:TTgg | 0.375 | 0.125 | 0.75 | 0.25 | 0.52 | 0.47 | 0 | 0.01 |
| $\Theta_4$ | 2H1 | H1:TTgg | 0.375 | 0.125 | 0.75 | 0.5 | 0.58 | 0.42 | 0 | 0 |
| $\Theta_4$ | 2H1 | H1:TTgg | 0.375 | 0.125 | 0.75 | 0.75 | 0.61 | 0.39 | 0 | 0 |
| $\Theta_4$ | 2H1 | H1:TTgg | 0.375 | 1.125 | 0.25 | 0.25 | 0.65 | 0.35 | 0 | 0 |
| $\Theta_4$ | 2H1 | H1:TTgg | 0.375 | 1.125 | 0.25 | 0.5 | 0.66 | 0.34 | 0 | 0 |
| $\Theta_4$ | 2H1 | H1:TTgg | 0.375 | 1.125 | 0.25 | 0.75 | 0.65 | 0.35 | 0 | 0 |
| $\Theta_4$ | 2H1 | H1:TTgg | 0.375 | 1.125 | 0.5 | 0.25 | 0.81 | 0.19 | 0 | 0 |
| $\Theta_4$ | 2H1 | H1:TTgg | 0.375 | 1.125 | 0.5 | 0.5 | 0.81 | 0.19 | 0 | 0 |
| $\Theta_4$ | 2H1 | H1:TTgg | 0.375 | 1.125 | 0.5 | 0.75 | 0.83 | 0.17 | 0 | 0 |
| $\Theta_4$ | 2H1 | H1:TTgg | 0.375 | 1.125 | 0.75 | 0.25 | 0.65 | 0.35 | 0 | 0 |
| $\Theta_4$ | 2H1 | H1:TTgg | 0.375 | 1.125 | 0.75 | 0.5 | 0.66 | 0.34 | 0 | 0 |
| $\Theta_4$ | 2H1 | H1:TTgg | 0.375 | 1.125 | 0.75 | 0.75 | 0.64 | 0.36 | 0 | 0 |
| $\Theta_4$ | 2H1 | H1:TTgg | 0.5 | 0.5 | 0.25 | 0.25 | 0.70 | 0.27 | 0 | 0.03 |

| | | | | | | | | | | |
|---|---|---|---|---|---|---|---|---|---|---|
| **$\Theta_4$** | **2H1** | **H1:TTgg** | **0.5** | **0.5** | **0.25** | **0.5** | 0.75 | 0.25 | 0 | 0 |
| **$\Theta_4$** | **2H1** | **H1:TTgg** | **0.5** | **0.5** | **0.25** | **0.75** | 0.72 | 0.28 | 0 | 0 |
| **$\Theta_4$** | **2H1** | **H1:TTgg** | **0.5** | **0.5** | **0.5** | **0.25** | 0.87 | 0.12 | 0 | 0.01 |
| **$\Theta_4$** | **2H1** | **H1:TTgg** | **0.5** | **0.5** | **0.5** | **0.5** | 0.96 | 0.04 | 0 | 0 |
| **$\Theta_4$** | **2H1** | **H1:TTgg** | **0.5** | **0.5** | **0.5** | **0.75** | 0.95 | 0.05 | 0 | 0 |
| **$\Theta_4$** | **2H1** | **H1:TTgg** | **0.5** | **0.5** | **0.75** | **0.25** | 0.77 | 0.23 | 0 | 0 |
| **$\Theta_4$** | **2H1** | **H1:TTgg** | **0.5** | **0.5** | **0.75** | **0.5** | 0.80 | 0.20 | 0 | 0 |
| **$\Theta_4$** | **2H1** | **H1:TTgg** | **0.5** | **0.5** | **0.75** | **0.75** | 0.80 | 0.20 | 0 | 0 |
| **$\Theta_4$** | **2H1** | **H1:TTgg** | **0.75** | **0.25** | **0.25** | **0.25** | 0.60 | 0.37 | 0 | 0.03 |
| **$\Theta_4$** | **2H1** | **H1:TTgg** | **0.75** | **0.25** | **0.25** | **0.5** | 0.73 | 0.26 | 0 | 0.01 |
| **$\Theta_4$** | **2H1** | **H1:TTgg** | **0.75** | **0.25** | **0.25** | **0.75** | 0.72 | 0.28 | 0 | 0 |
| **$\Theta_4$** | **2H1** | **H1:TTgg** | **0.75** | **0.25** | **0.5** | **0.25** | 0.69 | 0.26 | 0 | 0.05 |
| **$\Theta_4$** | **2H1** | **H1:TTgg** | **0.75** | **0.25** | **0.5** | **0.5** | 0.90 | 0.10 | 0 | 0 |
| **$\Theta_4$** | **2H1** | **H1:TTgg** | **0.75** | **0.25** | **0.5** | **0.75** | 0.93 | 0.07 | 0 | 0 |
| **$\Theta_4$** | **2H1** | **H1:TTgg** | **0.75** | **0.25** | **0.75** | **0.25** | 0.75 | 0.21 | 0 | 0.04 |
| **$\Theta_4$** | **2H1** | **H1:TTgg** | **0.75** | **0.25** | **0.75** | **0.5** | 0.92 | 0.08 | 0 | 0 |
| **$\Theta_4$** | **2H1** | **H1:TTgg** | **0.75** | **0.25** | **0.75** | **0.75** | 0.96 | 0.04 | 0 | 0 |
| **$\Theta_4$** | **2H1** | **H1:TTgg** | **0.75** | **0.75** | **0.25** | **0.25** | 0.92 | 0.07 | 0 | 0.01 |
| **$\Theta_4$** | **2H1** | **H1:TTgg** | **0.75** | **0.75** | **0.25** | **0.5** | 0.94 | 0.06 | 0 | 0 |
| **$\Theta_4$** | **2H1** | **H1:TTgg** | **0.75** | **0.75** | **0.25** | **0.75** | 0.92 | 0.08 | 0 | 0 |
| **$\Theta_4$** | **2H1** | **H1:TTgg** | **0.75** | **0.75** | **0.5** | **0.25** | 0.99 | 0.01 | 0 | 0 |
| **$\Theta_4$** | **2H1** | **H1:TTgg** | **0.75** | **0.75** | **0.5** | **0.5** | 1 | 0 | 0 | 0 |
| **$\Theta_4$** | **2H1** | **H1:TTgg** | **0.75** | **0.75** | **0.5** | **0.75** | 1 | 0 | 0 | 0 |
| **$\Theta_4$** | **2H1** | **H1:TTgg** | **0.75** | **0.75** | **0.75** | **0.25** | 0.93 | 0.07 | 0 | 0 |
| **$\Theta_4$** | **2H1** | **H1:TTgg** | **0.75** | **0.75** | **0.75** | **0.5** | 0.94 | 0.06 | 0 | 0 |
| **$\Theta_4$** | **2H1** | **H1:TTgg** | **0.75** | **0.75** | **0.75** | **0.75** | 0.94 | 0.06 | 0 | 0 |
| **$\Theta_4$** | **2H1** | **H1:TTgg** | **1.125** | **0.375** | **0.25** | **0.25** | 0.74 | 0.22 | 0 | 0.04 |
| **$\Theta_4$** | **2H1** | **H1:TTgg** | **1.125** | **0.375** | **0.25** | **0.5** | 0.92 | 0.08 | 0 | 0 |
| **$\Theta_4$** | **2H1** | **H1:TTgg** | **1.125** | **0.375** | **0.25** | **0.75** | 0.89 | 0.11 | 0 | 0 |
| **$\Theta_4$** | **2H1** | **H1:TTgg** | **1.125** | **0.375** | **0.5** | **0.25** | 0.85 | 0.11 | 0 | 0.04 |
| **$\Theta_4$** | **2H1** | **H1:TTgg** | **1.125** | **0.375** | **0.5** | **0.5** | 0.99 | 0.01 | 0 | 0 |
| **$\Theta_4$** | **2H1** | **H1:TTgg** | **1.125** | **0.375** | **0.5** | **0.75** | 0.99 | 0.01 | 0 | 0 |
| **$\Theta_4$** | **2H1** | **H1:TTgg** | **1.125** | **0.375** | **0.75** | **0.25** | 0.91 | 0.07 | 0 | 0.02 |
| **$\Theta_4$** | **2H1** | **H1:TTgg** | **1.125** | **0.375** | **0.75** | **0.5** | 0.99 | 0.01 | 0 | 0 |
| **$\Theta_4$** | **2H1** | **H1:TTgg** | **1.125** | **0.375** | **0.75** | **0.75** | 1 | 0 | 0 | 0 |
| **$\Theta_4$** | **2H2** | **H2:TTgg** | **0.125** | **0.375** | **0.25** | **0.25** | 0.52 | 0.48 | 0 | 0 |
| **$\Theta_4$** | **2H2** | **H2:TTgg** | **0.125** | **0.375** | **0.25** | **0.5** | 0.52 | 0.48 | 0 | 0 |
| **$\Theta_4$** | **2H2** | **H2:TTgg** | **0.125** | **0.375** | **0.25** | **0.75** | 0.52 | 0.48 | 0 | 0 |
| **$\Theta_4$** | **2H2** | **H2:TTgg** | **0.125** | **0.375** | **0.5** | **0.25** | 0.53 | 0.47 | 0 | 0 |
| **$\Theta_4$** | **2H2** | **H2:TTgg** | **0.125** | **0.375** | **0.5** | **0.5** | 0.53 | 0.47 | 0 | 0 |
| **$\Theta_4$** | **2H2** | **H2:TTgg** | **0.125** | **0.375** | **0.5** | **0.75** | 0.54 | 0.46 | 0 | 0 |
| **$\Theta_4$** | **2H2** | **H2:TTgg** | **0.25** | **0.25** | **0.25** | **0.25** | 0.53 | 0.47 | 0 | 0 |
| **$\Theta_4$** | **2H2** | **H2:TTgg** | **0.25** | **0.25** | **0.25** | **0.5** | 0.54 | 0.46 | 0 | 0 |
| **$\Theta_4$** | **2H2** | **H2:TTgg** | **0.25** | **0.25** | **0.25** | **0.75** | 0.54 | 0.46 | 0 | 0 |
| **$\Theta_4$** | **2H2** | **H2:TTgg** | **0.25** | **0.25** | **0.5** | **0.25** | 0.60 | 0.40 | 0 | 0 |
| **$\Theta_4$** | **2H2** | **H2:TTgg** | **0.25** | **0.25** | **0.5** | **0.5** | 0.59 | 0.41 | 0 | 0 |
| **$\Theta_4$** | **2H2** | **H2:TTgg** | **0.25** | **0.25** | **0.5** | **0.75** | 0.60 | 0.40 | 0 | 0 |
| **$\Theta_4$** | **2H2** | **H2:TTgg** | **0.25** | **0.75** | **0.25** | **0.25** | 0.58 | 0.42 | 0 | 0 |
| **$\Theta_4$** | **2H2** | **H2:TTgg** | **0.25** | **0.75** | **0.25** | **0.5** | 0.59 | 0.41 | 0 | 0 |
| **$\Theta_4$** | **2H2** | **H2:TTgg** | **0.25** | **0.75** | **0.25** | **0.75** | 0.58 | 0.42 | 0 | 0 |

| | | | | | | | | | | |
|---|---|---|---|---|---|---|---|---|---|---|
| **$\Theta_4$** | **2H2** | **H2:TTgg** | **0.25** | **0.75** | **0.5** | **0.25** | 0.65 | 0.35 | 0 | 0 |
| **$\Theta_4$** | **2H2** | **H2:TTgg** | **0.25** | **0.75** | **0.5** | **0.5** | 0.66 | 0.34 | 0 | 0 |
| **$\Theta_4$** | **2H2** | **H2:TTgg** | **0.25** | **0.75** | **0.5** | **0.75** | 0.65 | 0.35 | 0 | 0 |
| **$\Theta_4$** | **2H2** | **H2:TTgg** | **0.375** | **0.125** | **0.25** | **0.25** | 0.52 | 0.48 | 0 | 0 |
| **$\Theta_4$** | **2H2** | **H2:TTgg** | **0.375** | **0.125** | **0.25** | **0.5** | 0.52 | 0.47 | 0 | 0.01 |
| **$\Theta_4$** | **2H2** | **H2:TTgg** | **0.375** | **0.125** | **0.25** | **0.75** | 0.53 | 0.46 | 0 | 0.01 |
| **$\Theta_4$** | **2H2** | **H2:TTgg** | **0.375** | **0.125** | **0.5** | **0.25** | 0.57 | 0.43 | 0 | 0 |
| **$\Theta_4$** | **2H2** | **H2:TTgg** | **0.375** | **0.125** | **0.5** | **0.5** | 0.57 | 0.43 | 0 | 0 |
| **$\Theta_4$** | **2H2** | **H2:TTgg** | **0.375** | **0.125** | **0.5** | **0.75** | 0.57 | 0.42 | 0 | 0.01 |
| **$\Theta_4$** | **2H2** | **H2:TTgg** | **0.375** | **1.125** | **0.25** | **0.25** | 0.64 | 0.36 | 0 | 0 |
| **$\Theta_4$** | **2H2** | **H2:TTgg** | **0.375** | **1.125** | **0.25** | **0.5** | 0.65 | 0.35 | 0 | 0 |
| **$\Theta_4$** | **2H2** | **H2:TTgg** | **0.375** | **1.125** | **0.25** | **0.75** | 0.64 | 0.36 | 0 | 0 |
| **$\Theta_4$** | **2H2** | **H2:TTgg** | **0.375** | **1.125** | **0.5** | **0.25** | 0.82 | 0.18 | 0 | 0 |
| **$\Theta_4$** | **2H2** | **H2:TTgg** | **0.375** | **1.125** | **0.5** | **0.5** | 0.81 | 0.19 | 0 | 0 |
| **$\Theta_4$** | **2H2** | **H2:TTgg** | **0.375** | **1.125** | **0.5** | **0.75** | 0.80 | 0.20 | 0 | 0 |
| **$\Theta_4$** | **2H2** | **H2:TTgg** | **0.5** | **0.5** | **0.25** | **0.25** | 0.79 | 0.21 | 0 | 0 |
| **$\Theta_4$** | **2H2** | **H2:TTgg** | **0.5** | **0.5** | **0.25** | **0.5** | 0.79 | 0.21 | 0 | 0 |
| **$\Theta_4$** | **2H2** | **H2:TTgg** | **0.5** | **0.5** | **0.25** | **0.75** | 0.77 | 0.23 | 0 | 0 |
| **$\Theta_4$** | **2H2** | **H2:TTgg** | **0.5** | **0.5** | **0.5** | **0.25** | 0.97 | 0.03 | 0 | 0 |
| **$\Theta_4$** | **2H2** | **H2:TTgg** | **0.5** | **0.5** | **0.5** | **0.5** | 0.97 | 0.03 | 0 | 0 |
| **$\Theta_4$** | **2H2** | **H2:TTgg** | **0.5** | **0.5** | **0.5** | **0.75** | 0.97 | 0.03 | 0 | 0 |
| **$\Theta_4$** | **2H2** | **H2:TTgg** | **0.75** | **0.25** | **0.25** | **0.25** | 0.84 | 0.16 | 0 | 0 |
| **$\Theta_4$** | **2H2** | **H2:TTgg** | **0.75** | **0.25** | **0.25** | **0.5** | 0.80 | 0.20 | 0 | 0 |
| **$\Theta_4$** | **2H2** | **H2:TTgg** | **0.75** | **0.25** | **0.25** | **0.75** | 0.76 | 0.23 | 0 | 0.01 |
| **$\Theta_4$** | **2H2** | **H2:TTgg** | **0.75** | **0.25** | **0.5** | **0.25** | 0.84 | 0.16 | 0 | 0 |
| **$\Theta_4$** | **2H2** | **H2:TTgg** | **0.75** | **0.25** | **0.5** | **0.5** | 0.84 | 0.16 | 0 | 0 |
| **$\Theta_4$** | **2H2** | **H2:TTgg** | **0.75** | **0.25** | **0.5** | **0.75** | 0.83 | 0.16 | 0 | 0.01 |
| **$\Theta_4$** | **2H2** | **H2:TTgg** | **0.75** | **0.75** | **0.25** | **0.25** | 0.94 | 0.06 | 0 | 0 |
| **$\Theta_4$** | **2H2** | **H2:TTgg** | **0.75** | **0.75** | **0.25** | **0.5** | 0.93 | 0.07 | 0 | 0 |
| **$\Theta_4$** | **2H2** | **H2:TTgg** | **0.75** | **0.75** | **0.25** | **0.75** | 0.91 | 0.09 | 0 | 0 |
| **$\Theta_4$** | **2H2** | **H2:TTgg** | **0.75** | **0.75** | **0.5** | **0.25** | 1 | 0 | 0 | 0 |
| **$\Theta_4$** | **2H2** | **H2:TTgg** | **0.75** | **0.75** | **0.5** | **0.5** | 1 | 0 | 0 | 0 |
| **$\Theta_4$** | **2H2** | **H2:TTgg** | **0.75** | **0.75** | **0.5** | **0.75** | 0.97 | 0.03 | 0 | 0 |
| **$\Theta_4$** | **2H2** | **H2:TTgg** | **1.125** | **0.375** | **0.25** | **0.25** | 0.96 | 0.04 | 0 | 0 |
| **$\Theta_4$** | **2H2** | **H2:TTgg** | **1.125** | **0.375** | **0.25** | **0.5** | 0.95 | 0.05 | 0 | 0 |
| **$\Theta_4$** | **2H2** | **H2:TTgg** | **1.125** | **0.375** | **0.25** | **0.75** | 0.91 | 0.08 | 0 | 0.01 |
| **$\Theta_4$** | **2H2** | **H2:TTgg** | **1.125** | **0.375** | **0.5** | **0.25** | 0.94 | 0.06 | 0 | 0 |
| **$\Theta_4$** | **2H2** | **H2:TTgg** | **1.125** | **0.375** | **0.5** | **0.5** | 0.94 | 0.06 | 0 | 0 |
| **$\Theta_4$** | **2H2** | **H2:TTgg** | **1.125** | **0.375** | **0.5** | **0.75** | 0.92 | 0.08 | 0 | 0 |

### Table S6.

**Testing the *stepwise* method and empirical distribution criterion (eCDF) for frequency of correct recognition of models in the Poisson randomized set.** The first column indicates group *Θj*, the following two columns show the trivial name of the model and its alias in TTgg notation, and the following four columns describe the initial values of parameters T1, T3, $\gamma_1$, $\gamma_3$ taken in a Poisson-distribution based randomization process. The last four columns present the frequency of the correct model recognition: $\boldsymbol{\Theta_j^i}$; frequency of error recognition of the model in the direction of a simple model:***Θj-***; frequency of error recognition of the model to a complex model: ***Θj+***; frequency of error recognition of the model because the optimization process selects another model from the same group: *Θj*.

| Group | Model | Alias | $T_1$ | $T_3$ | $\gamma_1$ | $\gamma_3$ | $\Theta_j^i$ | $\Theta_{j-}$ | $\Theta_{j+}$ | $\Theta_j$ |
|---|---|---|---|---|---|---|---|---|---|---|
| $\Theta_0$ | P | H1:00nn | 0 | 0 | - | - | 0.96 | 0 | 0.04 | 0 |
| $\Theta_1$ | TP | H1:0Tn1 | 0 | 0.5 | - | 1 | 0.96 | 0 | 0.04 | 0 |
| $\Theta_1$ | TP | H1:0Tn1 | 0 | 1 | - | 1 | 0.96 | 0 | 0.04 | 0 |
| $\Theta_1$ | TP | H1:0Tn1 | 0 | 1.5 | - | 1 | 0.94 | 0 | 0.06 | 0 |
| $\Theta_1$ | PT | H1:T00n | 0.5 | 0 | 0 | - | 0.95 | 0 | 0.05 | 0 |
| $\Theta_1$ | PT | H1:T00n | 1 | 0 | 0 | - | 0.96 | 0 | 0.04 | 0 |
| $\Theta_1$ | PT | H1:T00n | 1.5 | 0 | 0 | - | 0.95 | 0 | 0.05 | 0 |
| $\Theta_2$ | 1HP1 | H1:0Tng | 0 | 0.5 | - | 0.25 | 0.61 | 0.32 | 0.05 | 0.02 |
| $\Theta_2$ | 1HP1 | H1:0Tng | 0 | 0.5 | - | 0.5 | 0.91 | 0.07 | 0.01 | 0.01 |
| $\Theta_2$ | 1HP1 | H1:0Tng | 0 | 1 | - | 0.25 | 0.97 | 0 | 0.03 | 0 |
| $\Theta_2$ | 1HP1 | H1:0Tng | 0 | 1 | - | 0.5 | 0.96 | 0 | 0.04 | 0 |
| $\Theta_2$ | 1HP1 | H1:0Tng | 0 | 1.5 | - | 0.25 | 0.95 | 0 | 0.05 | 0 |
| $\Theta_2$ | 1HP1 | H1:0Tng | 0 | 1.5 | - | 0.5 | 0.96 | 0 | 0.04 | 0 |
| $\Theta_2$ | 1HP2 | H1:T0g0 | 0.5 | 0 | 0.25 | 0 | 0.79 | 0.06 | 0.05 | 0.1 |
| $\Theta_2$ | 1HP2 | H1:T0g0 | 0.5 | 0 | 0.5 | 0 | 0.95 | 0.03 | 0.02 | 0 |
| $\Theta_2$ | 1HP2 | H1:T0g0 | 0.5 | 0 | 0.75 | 0 | 0.90 | 0 | 0.05 | 0.05 |
| $\Theta_2$ | 1HP2 | H1:T0g0 | 1 | 0 | 0.25 | 0 | 0.95 | 0 | 0.05 | 0 |
| $\Theta_2$ | 1HP2 | H1:T0g0 | 1 | 0 | 0.5 | 0 | 0.95 | 0 | 0.05 | 0 |
| $\Theta_2$ | 1HP2 | H1:T0g0 | 1 | 0 | 0.75 | 0 | 0.96 | 0 | 0.04 | 0 |
| $\Theta_2$ | 1HP2 | H1:T0g0 | 1.5 | 0 | 0.25 | 0 | 0.95 | 0 | 0.05 | 0 |
| $\Theta_2$ | 1HP2 | H1:T0g0 | 1.5 | 0 | 0.5 | 0 | 0.95 | 0 | 0.05 | 0 |
| $\Theta_2$ | 1HP2 | H1:T0g0 | 1.5 | 0 | 0.75 | 0 | 0.94 | 0 | 0.06 | 0 |
| $\Theta_2$ | 1HP3 | H1:T0g1 | 0.5 | 0 | 0.5 | 1 | 0.74 | 0.16 | 0.04 | 0.06 |
| $\Theta_2$ | 1HP3 | H1:T0g1 | 0.5 | 0 | 0.25 | 1 | 0.92 | 0.04 | 0.03 | 0.01 |
| $\Theta_2$ | 1HP3 | H1:T0g1 | 1 | 0 | 0.5 | 1 | 0.96 | 0 | 0.04 | 0 |
| $\Theta_2$ | 1HP3 | H1:T0g1 | 1 | 0 | 0.25 | 1 | 0.95 | 0 | 0.05 | 0 |
| $\Theta_2$ | 1HP3 | H1:T0g1 | 1.5 | 0 | 0.5 | 1 | 0.95 | 0 | 0.05 | 0 |
| $\Theta_2$ | 1HP3 | H1:T0g1 | 1.5 | 0 | 0.25 | 1 | 0.95 | 0 | 0.05 | 0 |
| $\Theta_2$ | T1 | H1:TT10 | 0.125 | 0.375 | 1 | 0 | 0.42 | 0.45 | 0.08 | 0.05 |
| $\Theta_2$ | T1 | H1:TT10 | 0.25 | 0.25 | 1 | 0 | 0.76 | 0.08 | 0 | 0.16 |
| $\Theta_2$ | T1 | H1:TT10 | 0.25 | 0.75 | 1 | 0 | 0.84 | 0.12 | 0.04 | 0 |
| $\Theta_2$ | T1 | H1:TT10 | 0.375 | 0.125 | 1 | 0 | 0.56 | 0.09 | 0 | 0.35 |
| $\Theta_2$ | T1 | H1:TT10 | 0.375 | 1.125 | 1 | 0 | 0.94 | 0.03 | 0.03 | 0 |
| $\Theta_2$ | T1 | H1:TT10 | 0.5 | 0.5 | 1 | 0 | 0.94 | 0.02 | 0.03 | 0.01 |
| $\Theta_2$ | T1 | H1:TT10 | 0.75 | 0.25 | 1 | 0 | 0.69 | 0.03 | 0 | 0.28 |
| $\Theta_2$ | T1 | H1:TT10 | 0.75 | 0.75 | 1 | 0 | 0.94 | 0.03 | 0.03 | 0 |
| $\Theta_2$ | T1 | H1:TT10 | 1.125 | 0.375 | 1 | 0 | 0.76 | 0 | 0.06 | 0.18 |
| $\Theta_2$ | T2 | H1:TT01 | 0.125 | 0.375 | 0 | 1 | 0.38 | 0.40 | 0.14 | 0.08 |

| | | | | | | | | | | |
|---|---|---|---|---|---|---|---|---|---|---|
| $\Theta_2$ | T2 | H1:TT01 | 0.25 | 0.25 | 0 | 1 | 0.84 | 0.09 | 0.01 | 0.06 |
| $\Theta_2$ | T2 | H1:TT01 | 0.25 | 0.75 | 0 | 1 | 0.89 | 0.07 | 0.04 | 0 |
| $\Theta_2$ | T2 | H1:TT01 | 0.375 | 0.125 | 0 | 1 | 0.36 | 0.40 | 0.07 | 0.17 |
| $\Theta_2$ | T2 | H1:TT01 | 0.375 | 1.125 | 0 | 1 | 0.95 | 0 | 0.05 | 0 |
| $\Theta_2$ | T2 | H1:TT01 | 0.5 | 0.5 | 0 | 1 | 0.95 | 0 | 0.05 | 0 |
| $\Theta_2$ | T2 | H1:TT01 | 0.75 | 0.25 | 0 | 1 | 0.85 | 0.1 | 0.02 | 0.03 |
| $\Theta_2$ | T2 | H1:TT01 | 0.75 | 0.75 | 0 | 1 | 0.95 | 0 | 0.05 | 0 |
| $\Theta_3$ | T2 | H1:TT01 | 1.125 | 0.375 | 0 | 0 | 0.94 | 0 | 0.06 | 0 |
| $\Theta_3$ | 1H1 | H1:TTg0 | 0.125 | 0.375 | 0.25 | 0 | 0.07 | 0.61 | 0.30 | 0.02 |
| $\Theta_3$ | 1H1 | H1:TTg0 | 0.125 | 0.375 | 0.5 | 0 | 0.08 | 0.61 | 0.30 | 0.01 |
| $\Theta_3$ | 1H1 | H1:TTg0 | 0.125 | 0.375 | 0.75 | 0 | 0.07 | 0.62 | 0.27 | 0.04 |
| $\Theta_3$ | 1H1 | H1:TTg0 | 0.25 | 0.25 | 0.25 | 0 | 0.09 | 0.60 | 0.28 | 0.03 |
| $\Theta_3$ | 1H1 | H1:TTg0 | 0.25 | 0.25 | 0.5 | 0 | 0.12 | 0.48 | 0.36 | 0.04 |
| $\Theta_3$ | 1H1 | H1:TTg0 | 0.25 | 0.25 | 0.75 | 0 | 0.09 | 0.45 | 0.38 | 0.08 |
| $\Theta_3$ | 1H1 | H1:TTg0 | 0.25 | 0.75 | 0.25 | 0 | 0.13 | 0.46 | 0.38 | 0.03 |
| $\Theta_3$ | 1H1 | H1:TTg0 | 0.25 | 0.75 | 0.5 | 0 | 0.20 | 0.46 | 0.32 | 0.02 |
| $\Theta_3$ | 1H1 | H1:TTg0 | 0.25 | 0.75 | 0.75 | 0 | 0.12 | 0.45 | 0.36 | 0.07 |
| $\Theta_3$ | 1H1 | H1:TTg0 | 0.375 | 0.125 | 0.25 | 0 | 0.07 | 0.44 | 0.45 | 0.04 |
| $\Theta_3$ | 1H1 | H1:TTg0 | 0.375 | 0.125 | 0.5 | 0 | 0.10 | 0.46 | 0.41 | 0.03 |
| $\Theta_3$ | 1H1 | H1:TTg0 | 0.375 | 0.125 | 0.75 | 0 | 0.07 | 0.45 | 0.42 | 0.06 |
| $\Theta_3$ | 1H1 | H1:TTg0 | 0.375 | 1.125 | 0.25 | 0 | 0.20 | 0.45 | 0.31 | 0.04 |
| $\Theta_3$ | 1H1 | H1:TTg0 | 0.375 | 1.125 | 0.5 | 0 | 0.43 | 0.39 | 0.16 | 0.02 |
| $\Theta_3$ | 1H1 | H1:TTg0 | 0.375 | 1.125 | 0.75 | 0 | 0.19 | 0.45 | 0.30 | 0.06 |
| $\Theta_3$ | 1H1 | H1:TTg0 | 0.5 | 0.5 | 0.25 | 0 | 0.39 | 0.40 | 0.20 | 0.01 |
| $\Theta_3$ | 1H1 | H1:TTg0 | 0.5 | 0.5 | 0.5 | 0 | 0.76 | 0.18 | 0.06 | 0 |
| $\Theta_3$ | 1H1 | H1:TTg0 | 0.5 | 0.5 | 0.75 | 0 | 0.35 | 0.42 | 0.17 | 0.06 |
| $\Theta_3$ | 1H1 | H1:TTg0 | 0.75 | 0.25 | 0.25 | 0 | 0.39 | 0.43 | 0.17 | 0.01 |
| $\Theta_3$ | 1H1 | H1:TTg0 | 0.75 | 0.25 | 0.5 | 0 | 0.39 | 0.39 | 0.22 | 0 |
| $\Theta_3$ | 1H1 | H1:TTg0 | 0.75 | 0.25 | 0.75 | 0 | 0.26 | 0.47 | 0.25 | 0.02 |
| $\Theta_3$ | 1H1 | H1:TTg0 | 0.75 | 0.75 | 0.25 | 0 | 0.71 | 0.21 | 0.07 | 0.01 |
| $\Theta_3$ | 1H1 | H1:TTg0 | 0.75 | 0.75 | 0.5 | 0 | 0.94 | 0.03 | 0.03 | 0 |
| $\Theta_3$ | 1H1 | H1:TTg0 | 0.75 | 0.75 | 0.75 | 0 | 0.65 | 0.25 | 0.08 | 0.02 |
| $\Theta_3$ | 1H1 | H1:TTg0 | 1.125 | 0.375 | 0.25 | 0 | 0.74 | 0.19 | 0.07 | 0 |
| $\Theta_3$ | 1H1 | H1:TTg0 | 1.125 | 0.375 | 0.5 | 0 | 0.59 | 0.29 | 0.12 | 0 |
| $\Theta_3$ | 1H1 | H1:TTg0 | 1.125 | 0.375 | 0.75 | 0 | 0.45 | 0.36 | 0.18 | 0.01 |
| $\Theta_3$ | 1H2 | H1:TT1g | 0.125 | 0.375 | 1 | 0.25 | 0.25 | 0.47 | 0.27 | 0.01 |
| $\Theta_3$ | 1H2 | H1:TT1g | 0.125 | 0.375 | 1 | 0.5 | 0.39 | 0.41 | 0.200 | 0 |
| $\Theta_3$ | 1H2 | H1:TT1g | 0.125 | 0.375 | 1 | 0.75 | 0.28 | 0.46 | 0.24 | 0.02 |
| $\Theta_3$ | 1H2 | H1:TT1g | 0.25 | 0.25 | 1 | 0.25 | 0.34 | 0.43 | 0.19 | 0.04 |
| $\Theta_3$ | 1H2 | H1:TT1g | 0.25 | 0.25 | 1 | 0.5 | 0.6 | 0.31 | 0.05 | 0.04 |
| $\Theta_3$ | 1H2 | H1:TT1g | 0.25 | 0.25 | 1 | 0.75 | 0.42 | 0.41 | 0.14 | 0.03 |
| $\Theta_3$ | 1H2 | H1:TT1g | 0.25 | 0.75 | 1 | 0.25 | 0.86 | 0.11 | 0.03 | 0 |
| $\Theta_3$ | 1H2 | H1:TT1g | 0.25 | 0.75 | 1 | 0.5 | 0.86 | 0.11 | 0.03 | 0 |
| $\Theta_3$ | 1H2 | H1:TT1g | 0.25 | 0.75 | 1 | 0.75 | 0.86 | 0.10 | 0.04 | 0 |
| $\Theta_3$ | 1H2 | H1:TT1g | 0.375 | 0.125 | 1 | 0.25 | 0.18 | 0.43 | 0.33 | 0.06 |
| $\Theta_3$ | 1H2 | H1:TT1g | 0.375 | 0.125 | 1 | 0.5 | 0.24 | 0.42 | 0.27 | 0.07 |
| $\Theta_3$ | 1H2 | H1:TT1g | 0.375 | 0.125 | 1 | 0.75 | 0.25 | 0.44 | 0.26 | 0.05 |
| $\Theta_3$ | 1H2 | H1:TT1g | 0.375 | 1.125 | 1 | 0.25 | 0.94 | 0 | 0.06 | 0 |
| $\Theta_3$ | 1H2 | H1:TT1g | 0.375 | 1.125 | 1 | 0.5 | 0.95 | 0 | 0.05 | 0 |
| $\Theta_3$ | 1H2 | H1:TT1g | 0.375 | 1.125 | 1 | 0.75 | 0.94 | 0 | 0.06 | 0 |

| | | | | | | | | | | |
|---|---|---|---|---|---|---|---|---|---|---|
| **$\Theta_3$** | **1H2** | **H1:TT1g** | **0.5** | **0.5** | **1** | **0.25** | 0.90 | 0.08 | 0.02 | 0 |
| **$\Theta_3$** | **1H2** | **H1:TT1g** | **0.5** | **0.5** | **1** | **0.5** | 0.94 | 0 | 0.06 | 0 |
| **$\Theta_3$** | **1H2** | **H1:TT1g** | **0.5** | **0.5** | **1** | **0.75** | 0.94 | 0 | 0.06 | 0 |
| **$\Theta_3$** | **1H2** | **H1:TT1g** | **0.75** | **0.25** | **1** | **0.25** | 0.41 | 0.39 | 0.13 | 0.07 |
| **$\Theta_3$** | **1H2** | **H1:TT1g** | **0.75** | **0.25** | **1** | **0.5** | 0.58 | 0.31 | 0.07 | 0.04 |
| **$\Theta_3$** | **1H2** | **H1:TT1g** | **0.75** | **0.25** | **1** | **0.75** | 0.71 | 0.22 | 0.02 | 0.05 |
| **$\Theta_3$** | **1H2** | **H1:TT1g** | **0.75** | **0.75** | **1** | **0.25** | 0.95 | 0 | 0.05 | 0 |
| **$\Theta_3$** | **1H2** | **H1:TT1g** | **0.75** | **0.75** | **1** | **0.5** | 0.95 | 0 | 0.05 | 0 |
| **$\Theta_3$** | **1H2** | **H1:TT1g** | **0.75** | **0.75** | **1** | **0.75** | 0.96 | 0 | 0.04 | 0 |
| **$\Theta_3$** | **1H2** | **H1:TT1g** | **1.125** | **0.375** | **1** | **0.25** | 0.62 | 0.29 | 0.07 | 0.02 |
| **$\Theta_3$** | **1H2** | **H1:TT1g** | **1.125** | **0.375** | **1** | **0.5** | 0.82 | 0.13 | 0.04 | 0.01 |
| **$\Theta_3$** | **1H2** | **H1:TT1g** | **1.125** | **0.375** | **1** | **0.75** | 0.91 | 0.06 | 0.02 | 0.01 |
| **$\Theta_3$** | **1H3** | **H1:TT0g** | **0.125** | **0.375** | **0** | **0.25** | 0.16 | 0.43 | 0.33 | 0.08 |
| **$\Theta_3$** | **1H3** | **H1:TT0g** | **0.125** | **0.375** | **0** | **0.5** | 0.25 | 0.45 | 0.24 | 0.06 |
| **$\Theta_3$** | **1H3** | **H1:TT0g** | **0.25** | **0.25** | **0** | **0.25** | 0.16 | 0.42 | 0.34 | 0.08 |
| **$\Theta_3$** | **1H3** | **H1:TT0g** | **0.25** | **0.25** | **0** | **0.5** | 0.21 | 0.43 | 0.28 | 0.08 |
| **$\Theta_3$** | **1H3** | **H1:TT0g** | **0.25** | **0.75** | **0** | **0.25** | 0.76 | 0.19 | 0.03 | 0.02 |
| **$\Theta_3$** | **1H3** | **H1:TT0g** | **0.25** | **0.75** | **0** | **0.5** | 0.89 | 0.08 | 0.03 | 0 |
| **$\Theta_3$** | **1H3** | **H1:TT0g** | **0.375** | **0.125** | **0** | **0.25** | 0.12 | 0.41 | 0.42 | 0.05 |
| **$\Theta_3$** | **1H3** | **H1:TT0g** | **0.375** | **0.125** | **0** | **0.5** | 0.12 | 0.41 | 0.43 | 0.04 |
| **$\Theta_3$** | **1H3** | **H1:TT0g** | **0.375** | **1.125** | **0** | **0.25** | 0.95 | 0 | 0.05 | 0 |
| **$\Theta_3$** | **1H3** | **H1:TT0g** | **0.375** | **1.125** | **0** | **0.5** | 0.95 | 0 | 0.05 | 0 |
| **$\Theta_3$** | **1H3** | **H1:TT0g** | **0.5** | **0.5** | **0** | **0.25** | 0.43 | 0.38 | 0.12 | 0.07 |
| **$\Theta_3$** | **1H3** | **H1:TT0g** | **0.5** | **0.5** | **0** | **0.5** | 0.77 | 0.17 | 0.03 | 0.03 |
| **$\Theta_3$** | **1H3** | **H1:TT0g** | **0.75** | **0.25** | **0** | **0.25** | 0.16 | 0.42 | 0.34 | 0.08 |
| **$\Theta_3$** | **1H3** | **H1:TT0g** | **0.75** | **0.25** | **0** | **0.5** | 0.21 | 0.43 | 0.30 | 0.06 |
| **$\Theta_3$** | **1H3** | **H1:TT0g** | **0.75** | **0.75** | **0** | **0.25** | 0.73 | 0.21 | 0.02 | 0.04 |
| **$\Theta_3$** | **1H3** | **H1:TT0g** | **0.75** | **0.75** | **0** | **0.5** | 0.95 | 0 | 0.05 | 0 |
| **$\Theta_3$** | **1H3** | **H1:TT0g** | **1.125** | **0.375** | **0** | **0.25** | 0.23 | 0.41 | 0.28 | 0.08 |
| **$\Theta_3$** | **1H3** | **H1:TT0g** | **1.125** | **0.375** | **0** | **0.5** | 0.42 | 0.38 | 0.14 | 0.06 |
| **$\Theta_3$** | **1H4** | **H1:TTg1** | **0.125** | **0.375** | **0.25** | **1** | 0.12 | 0.41 | 0.44 | 0.03 |
| **$\Theta_3$** | **1H4** | **H1:TTg1** | **0.125** | **0.375** | **0.5** | **1** | 0.12 | 0.42 | 0.44 | 0.02 |
| **$\Theta_3$** | **1H4** | **H1:TTg1** | **0.25** | **0.25** | **0.25** | **1** | 0.17 | 0.43 | 0.31 | 0.09 |
| **$\Theta_3$** | **1H4** | **H1:TTg1** | **0.25** | **0.25** | **0.5** | **1** | 0.29 | 0.43 | 0.23 | 0.05 |
| **$\Theta_3$** | **1H4** | **H1:TTg1** | **0.25** | **0.75** | **0.25** | **1** | 0.16 | 0.42 | 0.31 | 0.11 |
| **$\Theta_3$** | **1H4** | **H1:TTg1** | **0.25** | **0.75** | **0.5** | **1** | 0.24 | 0.43 | 0.29 | 0.04 |
| **$\Theta_3$** | **1H4** | **H1:TTg1** | **0.375** | **0.125** | **0.25** | **1** | 0.24 | 0.44 | 0.26 | 0.06 |
| **$\Theta_3$** | **1H4** | **H1:TTg1** | **0.375** | **0.125** | **0.5** | **1** | 0.48 | 0.37 | 0.13 | 0.02 |
| **$\Theta_3$** | **1H4** | **H1:TTg1** | **0.375** | **1.125** | **0.25** | **1** | 0.22 | 0.43 | 0.22 | 0.13 |
| **$\Theta_3$** | **1H4** | **H1:TTg1** | **0.375** | **1.125** | **0.5** | **1** | 0.44 | 0.38 | 0.14 | 0.04 |
| **$\Theta_3$** | **1H4** | **H1:TTg1** | **0.5** | **0.5** | **0.25** | **1** | 0.44 | 0.37 | 0.10 | 0.09 |
| **$\Theta_3$** | **1H4** | **H1:TTg1** | **0.5** | **0.5** | **0.5** | **1** | 0.86 | 0.09 | 0.04 | 0.01 |
| **$\Theta_3$** | **1H4** | **H1:TTg1** | **0.75** | **0.25** | **0.25** | **1** | 0.81 | 0.14 | 0.03 | 0.02 |
| **$\Theta_3$** | **1H4** | **H1:TTg1** | **0.75** | **0.25** | **0.5** | **1** | 0.96 | 0 | 0.04 | 0 |
| **$\Theta_3$** | **1H4** | **H1:TTg1** | **0.75** | **0.75** | **0.25** | **1** | 0.76 | 0.18 | 0.03 | 0.03 |
| **$\Theta_3$** | **1H4** | **H1:TTg1** | **0.75** | **0.75** | **0.5** | **1** | 0.94 | 0 | 0.06 | 0 |
| **$\Theta_3$** | **1H4** | **H1:TTg1** | **1.125** | **0.375** | **0.25** | **1** | 0.96 | 0 | 0.04 | 0 |
| **$\Theta_3$** | **1H4** | **H1:TTg1** | **1.125** | **0.375** | **0.5** | **1** | 0.95 | 0 | 0.05 | 0 |
| **$\Theta_3$** | **2PH** | **H2:T0gg** | **0.5** | **0** | **0.25** | **0.5** | 0.38 | 0.43 | 0.12 | 0.07 |
| **$\Theta_3$** | **2PH** | **H2:T0gg** | **0.5** | **0** | **0.25** | **0.75** | 0.32 | 0.42 | 0.16 | 0.1 |

| | | | | | | | | | | |
|---|---|---|---|---|---|---|---|---|---|---|
| **$\Theta_3$** | **2PH** | **H2:T0gg** | **1** | **0** | **0.25** | **0.5** | 0.92 | 0.04 | 0.04 | 0 |
| **$\Theta_3$** | **2PH** | **H2:T0gg** | **1** | **0** | **0.25** | **0.75** | 0.89 | 0.06 | 0.05 | 0 |
| **$\Theta_3$** | **2PH** | **H2:T0gg** | **1.5** | **0** | **0.25** | **0.5** | 0.94 | 0 | 0.06 | 0 |
| **$\Theta_3$** | **2PH** | **H2:T0gg** | **1.5** | **0** | **0.25** | **0.75** | 0.95 | 0 | 0.05 | 0 |
| **$\Theta_4$** | **2H1** | **H1:TTgg** | **0.125** | **0.375** | **0.25** | **0.25** | 0.19 | 0.80 | 0 | 0.01 |
| **$\Theta_4$** | **2H1** | **H1:TTgg** | **0.125** | **0.375** | **0.25** | **0.5** | 0.19 | 0.81 | 0 | 0 |
| **$\Theta_4$** | **2H1** | **H1:TTgg** | **0.125** | **0.375** | **0.25** | **0.75** | 0.20 | 0.80 | 0 | 0 |
| **$\Theta_4$** | **2H1** | **H1:TTgg** | **0.125** | **0.375** | **0.5** | **0.25** | 0.18 | 0.81 | 0 | 0.01 |
| **$\Theta_4$** | **2H1** | **H1:TTgg** | **0.125** | **0.375** | **0.5** | **0.5** | 0.20 | 0.80 | 0 | 0 |
| **$\Theta_4$** | **2H1** | **H1:TTgg** | **0.125** | **0.375** | **0.5** | **0.75** | 0.19 | 0.81 | 0 | 0 |
| **$\Theta_4$** | **2H1** | **H1:TTgg** | **0.125** | **0.375** | **0.75** | **0.25** | 0.18 | 0.81 | 0 | 0.01 |
| **$\Theta_4$** | **2H1** | **H1:TTgg** | **0.125** | **0.375** | **0.75** | **0.5** | 0.19 | 0.81 | 0 | 0 |
| **$\Theta_4$** | **2H1** | **H1:TTgg** | **0.125** | **0.375** | **0.75** | **0.75** | 0.19 | 0.80 | 0 | 0.01 |
| **$\Theta_4$** | **2H1** | **H1:TTgg** | **0.25** | **0.25** | **0.25** | **0.25** | 0.18 | 0.81 | 0 | 0.01 |
| **$\Theta_4$** | **2H1** | **H1:TTgg** | **0.25** | **0.25** | **0.25** | **0.5** | 0.20 | 0.80 | 0 | 0 |
| **$\Theta_4$** | **2H1** | **H1:TTgg** | **0.25** | **0.25** | **0.25** | **0.75** | 0.20 | 0.80 | 0 | 0 |
| **$\Theta_4$** | **2H1** | **H1:TTgg** | **0.25** | **0.25** | **0.5** | **0.25** | 0.21 | 0.78 | 0 | 0.01 |
| **$\Theta_4$** | **2H1** | **H1:TTgg** | **0.25** | **0.25** | **0.5** | **0.5** | 0.25 | 0.75 | 0 | 0 |
| **$\Theta_4$** | **2H1** | **H1:TTgg** | **0.25** | **0.25** | **0.5** | **0.75** | 0.22 | 0.78 | 0 | 0 |
| **$\Theta_4$** | **2H1** | **H1:TTgg** | **0.25** | **0.25** | **0.75** | **0.25** | 0.19 | 0.80 | 0 | 0.01 |
| **$\Theta_4$** | **2H1** | **H1:TTgg** | **0.25** | **0.25** | **0.75** | **0.5** | 0.23 | 0.77 | 0 | 0 |
| **$\Theta_4$** | **2H1** | **H1:TTgg** | **0.25** | **0.25** | **0.75** | **0.75** | 0.23 | 0.77 | 0 | 0 |
| **$\Theta_4$** | **2H1** | **H1:TTgg** | **0.25** | **0.75** | **0.25** | **0.25** | 0.25 | 0.74 | 0 | 0.01 |
| **$\Theta_4$** | **2H1** | **H1:TTgg** | **0.25** | **0.75** | **0.25** | **0.5** | 0.25 | 0.75 | 0 | 0 |
| **$\Theta_4$** | **2H1** | **H1:TTgg** | **0.25** | **0.75** | **0.25** | **0.75** | 0.25 | 0.75 | 0 | 0 |
| **$\Theta_4$** | **2H1** | **H1:TTgg** | **0.25** | **0.75** | **0.5** | **0.25** | 0.30 | 0.70 | 0 | 0 |
| **$\Theta_4$** | **2H1** | **H1:TTgg** | **0.25** | **0.75** | **0.5** | **0.5** | 0.31 | 0.69 | 0 | 0 |
| **$\Theta_4$** | **2H1** | **H1:TTgg** | **0.25** | **0.75** | **0.5** | **0.75** | 0.32 | 0.68 | 0 | 0 |
| **$\Theta_4$** | **2H1** | **H1:TTgg** | **0.25** | **0.75** | **0.75** | **0.25** | 0.25 | 0.75 | 0 | 0 |
| **$\Theta_4$** | **2H1** | **H1:TTgg** | **0.25** | **0.75** | **0.75** | **0.5** | 0.27 | 0.73 | 0 | 0 |
| **$\Theta_4$** | **2H1** | **H1:TTgg** | **0.25** | **0.75** | **0.75** | **0.75** | 0.24 | 0.76 | 0 | 0 |
| **$\Theta_4$** | **2H1** | **H1:TTgg** | **0.375** | **0.125** | **0.25** | **0.25** | 0.19 | 0.81 | 0 | 0 |
| **$\Theta_4$** | **2H1** | **H1:TTgg** | **0.375** | **0.125** | **0.25** | **0.5** | 0.19 | 0.81 | 0 | 0 |
| **$\Theta_4$** | **2H1** | **H1:TTgg** | **0.375** | **0.125** | **0.25** | **0.75** | 0.20 | 0.80 | 0 | 0 |
| **$\Theta_4$** | **2H1** | **H1:TTgg** | **0.375** | **0.125** | **0.5** | **0.25** | 0.20 | 0.79 | 0 | 0.01 |
| **$\Theta_4$** | **2H1** | **H1:TTgg** | **0.375** | **0.125** | **0.5** | **0.5** | 0.25 | 0.75 | 0 | 0 |
| **$\Theta_4$** | **2H1** | **H1:TTgg** | **0.375** | **0.125** | **0.5** | **0.75** | 0.26 | 0.74 | 0 | 0 |
| **$\Theta_4$** | **2H1** | **H1:TTgg** | **0.375** | **0.125** | **0.75** | **0.25** | 0.18 | 0.81 | 0 | 0.01 |
| **$\Theta_4$** | **2H1** | **H1:TTgg** | **0.375** | **0.125** | **0.75** | **0.5** | 0.24 | 0.76 | 0 | 0 |
| **$\Theta_4$** | **2H1** | **H1:TTgg** | **0.375** | **0.125** | **0.75** | **0.75** | 0.26 | 0.74 | 0 | 0 |
| **$\Theta_4$** | **2H1** | **H1:TTgg** | **0.375** | **1.125** | **0.25** | **0.25** | 0.33 | 0.67 | 0 | 0 |
| **$\Theta_4$** | **2H1** | **H1:TTgg** | **0.375** | **1.125** | **0.25** | **0.5** | 0.33 | 0.67 | 0 | 0 |
| **$\Theta_4$** | **2H1** | **H1:TTgg** | **0.375** | **1.125** | **0.25** | **0.75** | 0.33 | 0.67 | 0 | 0 |
| **$\Theta_4$** | **2H1** | **H1:TTgg** | **0.375** | **1.125** | **0.5** | **0.25** | 0.50 | 0.50 | 0 | 0 |
| **$\Theta_4$** | **2H1** | **H1:TTgg** | **0.375** | **1.125** | **0.5** | **0.5** | 0.50 | 0.50 | 0 | 0 |
| **$\Theta_4$** | **2H1** | **H1:TTgg** | **0.375** | **1.125** | **0.5** | **0.75** | 0.51 | 0.49 | 0 | 0 |
| **$\Theta_4$** | **2H1** | **H1:TTgg** | **0.375** | **1.125** | **0.75** | **0.25** | 0.33 | 0.67 | 0 | 0 |
| **$\Theta_4$** | **2H1** | **H1:TTgg** | **0.375** | **1.125** | **0.75** | **0.5** | 0.33 | 0.67 | 0 | 0 |
| **$\Theta_4$** | **2H1** | **H1:TTgg** | **0.375** | **1.125** | **0.75** | **0.75** | 0.32 | 0.68 | 0 | 0 |
| **$\Theta_4$** | **2H1** | **H1:TTgg** | **0.5** | **0.5** | **0.25** | **0.25** | 0.34 | 0.63 | 0 | 0.03 |

| | | | | | | | | | | |
|---|---|---|---|---|---|---|---|---|---|---|
| $\Theta_4$ | **2H1** | **H1:TTgg** | **0.5** | **0.5** | **0.25** | **0.5** | 0.40 | 0.60 | 0 | 0 |
| $\Theta_4$ | **2H1** | **H1:TTgg** | **0.5** | **0.5** | **0.25** | **0.75** | 0.35 | 0.65 | 0 | 0 |
| $\Theta_4$ | **2H1** | **H1:TTgg** | **0.5** | **0.5** | **0.5** | **0.25** | 0.56 | 0.43 | 0 | 0.01 |
| $\Theta_4$ | **2H1** | **H1:TTgg** | **0.5** | **0.5** | **0.5** | **0.5** | 0.76 | 0.24 | 0 | 0 |
| $\Theta_4$ | **2H1** | **H1:TTgg** | **0.5** | **0.5** | **0.5** | **0.75** | 0.72 | 0.28 | 0 | 0 |
| $\Theta_4$ | **2H1** | **H1:TTgg** | **0.5** | **0.5** | **0.75** | **0.25** | 0.43 | 0.57 | 0 | 0 |
| $\Theta_4$ | **2H1** | **H1:TTgg** | **0.5** | **0.5** | **0.75** | **0.5** | 0.48 | 0.52 | 0 | 0 |
| $\Theta_4$ | **2H1** | **H1:TTgg** | **0.5** | **0.5** | **0.75** | **0.75** | 0.50 | 0.50 | 0 | 0 |
| $\Theta_4$ | **2H1** | **H1:TTgg** | **0.75** | **0.25** | **0.25** | **0.25** | 0.25 | 0.72 | 0 | 0.03 |
| $\Theta_4$ | **2H1** | **H1:TTgg** | **0.75** | **0.25** | **0.25** | **0.5** | 0.37 | 0.62 | 0 | 0.01 |
| $\Theta_4$ | **2H1** | **H1:TTgg** | **0.75** | **0.25** | **0.25** | **0.75** | 0.35 | 0.65 | 0 | 0 |
| $\Theta_4$ | **2H1** | **H1:TTgg** | **0.75** | **0.25** | **0.5** | **0.25** | 0.33 | 0.62 | 0 | 0.05 |
| $\Theta_4$ | **2H1** | **H1:TTgg** | **0.75** | **0.25** | **0.5** | **0.5** | 0.63 | 0.37 | 0 | 0 |
| $\Theta_4$ | **2H1** | **H1:TTgg** | **0.75** | **0.25** | **0.5** | **0.75** | 0.68 | 0.32 | 0 | 0 |
| $\Theta_4$ | **2H1** | **H1:TTgg** | **0.75** | **0.25** | **0.75** | **0.25** | 0.37 | 0.59 | 0 | 0.04 |
| $\Theta_4$ | **2H1** | **H1:TTgg** | **0.75** | **0.25** | **0.75** | **0.5** | 0.65 | 0.35 | 0 | 0 |
| $\Theta_4$ | **2H1** | **H1:TTgg** | **0.75** | **0.25** | **0.75** | **0.75** | 0.77 | 0.23 | 0 | 0 |
| $\Theta_4$ | **2H1** | **H1:TTgg** | **0.75** | **0.75** | **0.25** | **0.25** | 0.67 | 0.32 | 0 | 0.01 |
| $\Theta_4$ | **2H1** | **H1:TTgg** | **0.75** | **0.75** | **0.25** | **0.5** | 0.72 | 0.28 | 0 | 0 |
| $\Theta_4$ | **2H1** | **H1:TTgg** | **0.75** | **0.75** | **0.25** | **0.75** | 0.68 | 0.32 | 0 | 0 |
| $\Theta_4$ | **2H1** | **H1:TTgg** | **0.75** | **0.75** | **0.5** | **0.25** | 0.94 | 0.06 | 0 | 0 |
| $\Theta_4$ | **2H1** | **H1:TTgg** | **0.75** | **0.75** | **0.5** | **0.5** | 0.99 | 0.01 | 0 | 0 |
| $\Theta_4$ | **2H1** | **H1:TTgg** | **0.75** | **0.75** | **0.5** | **0.75** | 0.98 | 0.02 | 0 | 0 |
| $\Theta_4$ | **2H1** | **H1:TTgg** | **0.75** | **0.75** | **0.75** | **0.25** | 0.72 | 0.28 | 0 | 0 |
| $\Theta_4$ | **2H1** | **H1:TTgg** | **0.75** | **0.75** | **0.75** | **0.5** | 0.73 | 0.27 | 0 | 0 |
| $\Theta_4$ | **2H1** | **H1:TTgg** | **0.75** | **0.75** | **0.75** | **0.75** | 0.75 | 0.25 | 0 | 0 |
| $\Theta_4$ | **2H1** | **H1:TTgg** | **1.125** | **0.375** | **0.25** | **0.25** | 0.39 | 0.57 | 0 | 0.04 |
| $\Theta_4$ | **2H1** | **H1:TTgg** | **1.125** | **0.375** | **0.25** | **0.5** | 0.68 | 0.32 | 0 | 0 |
| $\Theta_4$ | **2H1** | **H1:TTgg** | **1.125** | **0.375** | **0.25** | **0.75** | 0.59 | 0.41 | 0 | 0 |
| $\Theta_4$ | **2H1** | **H1:TTgg** | **1.125** | **0.375** | **0.5** | **0.25** | 0.58 | 0.38 | 0 | 0.04 |
| $\Theta_4$ | **2H1** | **H1:TTgg** | **1.125** | **0.375** | **0.5** | **0.5** | 0.93 | 0.07 | 0 | 0 |
| $\Theta_4$ | **2H1** | **H1:TTgg** | **1.125** | **0.375** | **0.5** | **0.75** | 0.89 | 0.11 | 0 | 0 |
| $\Theta_4$ | **2H1** | **H1:TTgg** | **1.125** | **0.375** | **0.75** | **0.25** | 0.65 | 0.33 | 0 | 0.02 |
| $\Theta_4$ | **2H1** | **H1:TTgg** | **1.125** | **0.375** | **0.75** | **0.5** | 0.94 | 0.06 | 0 | 0 |
| $\Theta_4$ | **2H1** | **H1:TTgg** | **1.125** | **0.375** | **0.75** | **0.75** | 0.98 | 0.02 | 0 | 0 |
| $\Theta_4$ | **2H2** | **H2:TTgg** | **0.125** | **0.375** | **0.25** | **0.25** | 0.19 | 0.81 | 0 | 0 |
| $\Theta_4$ | **2H2** | **H2:TTgg** | **0.125** | **0.375** | **0.25** | **0.5** | 0.20 | 0.80 | 0 | 0 |
| $\Theta_4$ | **2H2** | **H2:TTgg** | **0.125** | **0.375** | **0.25** | **0.75** | 0.20 | 0.80 | 0 | 0 |
| $\Theta_4$ | **2H2** | **H2:TTgg** | **0.125** | **0.375** | **0.5** | **0.25** | 0.20 | 0.80 | 0 | 0 |
| $\Theta_4$ | **2H2** | **H2:TTgg** | **0.125** | **0.375** | **0.5** | **0.5** | 0.20 | 0.80 | 0 | 0 |
| $\Theta_4$ | **2H2** | **H2:TTgg** | **0.125** | **0.375** | **0.5** | **0.75** | 0.20 | 0.80 | 0 | 0 |
| $\Theta_4$ | **2H2** | **H2:TTgg** | **0.25** | **0.25** | **0.25** | **0.25** | 0.21 | 0.79 | 0 | 0 |
| $\Theta_4$ | **2H2** | **H2:TTgg** | **0.25** | **0.25** | **0.25** | **0.5** | 0.22 | 0.78 | 0 | 0 |
| $\Theta_4$ | **2H2** | **H2:TTgg** | **0.25** | **0.25** | **0.25** | **0.75** | 0.21 | 0.79 | 0 | 0 |
| $\Theta_4$ | **2H2** | **H2:TTgg** | **0.25** | **0.25** | **0.5** | **0.25** | 0.25 | 0.75 | 0 | 0 |
| $\Theta_4$ | **2H2** | **H2:TTgg** | **0.25** | **0.25** | **0.5** | **0.5** | 0.25 | 0.75 | 0 | 0 |
| $\Theta_4$ | **2H2** | **H2:TTgg** | **0.25** | **0.25** | **0.5** | **0.75** | 0.25 | 0.75 | 0 | 0 |
| $\Theta_4$ | **2H2** | **H2:TTgg** | **0.25** | **0.75** | **0.25** | **0.25** | 0.25 | 0.75 | 0 | 0 |
| $\Theta_4$ | **2H2** | **H2:TTgg** | **0.25** | **0.75** | **0.25** | **0.5** | 0.27 | 0.73 | 0 | 0 |
| $\Theta_4$ | **2H2** | **H2:TTgg** | **0.25** | **0.75** | **0.25** | **0.75** | 0.26 | 0.74 | 0 | 0 |

| | | | | | | | | | | |
|---|---|---|---|---|---|---|---|---|---|---|
| $\Theta_4$ | 2H2 | H2:TTgg | 0.25 | 0.75 | 0.5 | 0.25 | 0.32 | 0.68 | 0 | 0 |
| $\Theta_4$ | 2H2 | H2:TTgg | 0.25 | 0.75 | 0.5 | 0.5 | 0.32 | 0.68 | 0 | 0 |
| $\Theta_4$ | 2H2 | H2:TTgg | 0.25 | 0.75 | 0.5 | 0.75 | 0.32 | 0.68 | 0 | 0 |
| $\Theta_4$ | 2H2 | H2:TTgg | 0.375 | 0.125 | 0.25 | 0.25 | 0.20 | 0.80 | 0 | 0 |
| $\Theta_4$ | 2H2 | H2:TTgg | 0.375 | 0.125 | 0.25 | 0.5 | 0.20 | 0.79 | 0 | 0.01 |
| $\Theta_4$ | 2H2 | H2:TTgg | 0.375 | 0.125 | 0.25 | 0.75 | 0.20 | 0.79 | 0 | 0.01 |
| $\Theta_4$ | 2H2 | H2:TTgg | 0.375 | 0.125 | 0.5 | 0.25 | 0.23 | 0.77 | 0 | 0 |
| $\Theta_4$ | 2H2 | H2:TTgg | 0.375 | 0.125 | 0.5 | 0.5 | 0.23 | 0.77 | 0 | 0 |
| $\Theta_4$ | 2H2 | H2:TTgg | 0.375 | 0.125 | 0.5 | 0.75 | 0.23 | 0.76 | 0 | 0.01 |
| $\Theta_4$ | 2H2 | H2:TTgg | 0.375 | 1.125 | 0.25 | 0.25 | 0.31 | 0.69 | 0 | 0 |
| $\Theta_4$ | 2H2 | H2:TTgg | 0.375 | 1.125 | 0.25 | 0.5 | 0.31 | 0.69 | 0 | 0 |
| $\Theta_4$ | 2H2 | H2:TTgg | 0.375 | 1.125 | 0.25 | 0.75 | 0.31 | 0.69 | 0 | 0 |
| $\Theta_4$ | 2H2 | H2:TTgg | 0.375 | 1.125 | 0.5 | 0.25 | 0.51 | 0.49 | 0 | 0 |
| $\Theta_4$ | 2H2 | H2:TTgg | 0.375 | 1.125 | 0.5 | 0.5 | 0.49 | 0.51 | 0 | 0 |
| $\Theta_4$ | 2H2 | H2:TTgg | 0.375 | 1.125 | 0.5 | 0.75 | 0.50 | 0.50 | 0 | 0 |
| $\Theta_4$ | 2H2 | H2:TTgg | 0.5 | 0.5 | 0.25 | 0.25 | 0.48 | 0.52 | 0 | 0 |
| $\Theta_4$ | 2H2 | H2:TTgg | 0.5 | 0.5 | 0.25 | 0.5 | 0.46 | 0.54 | 0 | 0 |
| $\Theta_4$ | 2H2 | H2:TTgg | 0.5 | 0.5 | 0.25 | 0.75 | 0.43 | 0.57 | 0 | 0 |
| $\Theta_4$ | 2H2 | H2:TTgg | 0.5 | 0.5 | 0.5 | 0.25 | 0.80 | 0.20 | 0 | 0 |
| $\Theta_4$ | 2H2 | H2:TTgg | 0.5 | 0.5 | 0.5 | 0.5 | 0.80 | 0.20 | 0 | 0 |
| $\Theta_4$ | 2H2 | H2:TTgg | 0.5 | 0.5 | 0.5 | 0.75 | 0.78 | 0.22 | 0 | 0 |
| $\Theta_4$ | 2H2 | H2:TTgg | 0.75 | 0.25 | 0.25 | 0.25 | 0.46 | 0.54 | 0 | 0 |
| $\Theta_4$ | 2H2 | H2:TTgg | 0.75 | 0.25 | 0.25 | 0.5 | 0.45 | 0.55 | 0 | 0 |
| $\Theta_4$ | 2H2 | H2:TTgg | 0.75 | 0.25 | 0.25 | 0.75 | 0.40 | 0.59 | 0 | 0.01 |
| $\Theta_4$ | 2H2 | H2:TTgg | 0.75 | 0.25 | 0.5 | 0.25 | 0.54 | 0.46 | 0 | 0 |
| $\Theta_4$ | 2H2 | H2:TTgg | 0.75 | 0.25 | 0.5 | 0.5 | 0.52 | 0.48 | 0 | 0 |
| $\Theta_4$ | 2H2 | H2:TTgg | 0.75 | 0.25 | 0.5 | 0.75 | 0.54 | 0.45 | 0 | 0.01 |
| $\Theta_4$ | 2H2 | H2:TTgg | 0.75 | 0.75 | 0.25 | 0.25 | 0.73 | 0.27 | 0 | 0 |
| $\Theta_4$ | 2H2 | H2:TTgg | 0.75 | 0.75 | 0.25 | 0.5 | 0.72 | 0.28 | 0 | 0 |
| $\Theta_4$ | 2H2 | H2:TTgg | 0.75 | 0.75 | 0.25 | 0.75 | 0.70 | 0.30 | 0 | 0 |
| $\Theta_4$ | 2H2 | H2:TTgg | 0.75 | 0.75 | 0.5 | 0.25 | 0.99 | 0.01 | 0 | 0 |
| $\Theta_4$ | 2H2 | H2:TTgg | 0.75 | 0.75 | 0.5 | 0.5 | 0.99 | 0.01 | 0 | 0 |
| $\Theta_4$ | 2H2 | H2:TTgg | 0.75 | 0.75 | 0.5 | 0.75 | 0.95 | 0.05 | 0 | 0 |
| $\Theta_4$ | 2H2 | H2:TTgg | 1.125 | 0.375 | 0.25 | 0.25 | 0.77 | 0.23 | 0 | 0 |
| $\Theta_4$ | 2H2 | H2:TTgg | 1.125 | 0.375 | 0.25 | 0.5 | 0.74 | 0.26 | 0 | 0 |
| $\Theta_4$ | 2H2 | H2:TTgg | 1.125 | 0.375 | 0.25 | 0.75 | 0.69 | 0.30 | 0 | 0.01 |
| $\Theta_4$ | 2H2 | H2:TTgg | 1.125 | 0.375 | 0.5 | 0.25 | 0.72 | 0.28 | 0 | 0 |
| $\Theta_4$ | 2H2 | H2:TTgg | 1.125 | 0.375 | 0.5 | 0.5 | 0.71 | 0.29 | 0 | 0 |
| $\Theta_4$ | 2H2 | H2:TTgg | 1.125 | 0.375 | 0.5 | 0.75 | 0.72 | 0.28 | 0 | 0 |

### Table S7.

**Testing the dependence of the number of user makers on the frequency of correct recognition of models in the Poisson randomized set.** The first column indicates group *Θj*, the following two columns show the trivial name of the model and its alias in TTgg notation, and the following four columns describe the initial values of parameters T1, T3, $\gamma_1$, $\gamma_3$ taken in a Poisson-distribution based randomization process. The next column indicated the number of markers (n) taken as an initial value and controlled in simulations. The last four columns present the correct model recognition frequency under different statistical criteria ($\chi^2$ and eCDF) and algorithms (*reverse* and *stepwise*).

| Group | Model | Alias | $T_1$ | $T_3$ | $\gamma_1$ | $\gamma_3$ | n | χ2 reverse | χ2 stepwise | eCDF reverse | eCDF stepwise |
|---|---|---|---|---|---|---|---|---|---|---|---|
| $\Theta_0$ | P | H1:00nn | 0 | 0 | - | - | 50 | 0.92 | 0.75 | 0.97 | 0.83 |
| | | | | | | | 100 | 0.90 | 0.74 | 0.97 | 0.81 |
| | | | | | | | 200 | 0.94 | 0.77 | 0.98 | 0.85 |
| | | | | | | | 300 | 0.93 | 0.76 | 0.98 | 0.96 |
| | | | | | | | 400 | 0.92 | 0.76 | 0.99 | 0.97 |
| | | | | | | | 1000 | 0.93 | 0.76 | 1.00 | 0.98 |
| $\Theta_1$ | TP | H1:0Tn1 | 0 | 0.5 | - | 1 | 50 | 0.85 | 0.79 | 0.89 | 0.87 |
| | | | | | | | 100 | 0.94 | 0.81 | 0.95 | 0.89 |
| | | | | | | | 200 | 0.93 | 0.84 | 0.96 | 0.92 |
| | | | | | | | 300 | 0.94 | 0.85 | 0.98 | 0.96 |
| | | | | | | | 400 | 0.93 | 0.84 | 0.98 | 0.92 |
| | | | | | | | 1000 | 0.92 | 0.85 | 0.99 | 0.94 |
| $\Theta_1$ | PT | H1:T00n | 0.5 | 0 | 0 | - | 50 | 0.85 | 0.79 | 0.88 | 0.83 |
| | | | | | | | 100 | 0.94 | 0.83 | 0.96 | 0.88 |
| | | | | | | | 200 | 0.91 | 0.81 | 0.96 | 0.89 |
| | | | | | | | 300 | 0.95 | 0.85 | 0.97 | 0.95 |
| | | | | | | | 400 | 0.93 | 0.81 | 0.95 | 0.97 |
| | | | | | | | 1000 | 0.92 | 0.81 | 0.99 | 0.99 |
| $\Theta_2$ | 1HP1 | H1:0Tng | 0 | 0.5 | - | 0.5 | 50 | 0.29 | 0.32 | 0.32 | 0.35 |
| | | | | | | | 100 | 0.63 | 0.62 | 0.69 | 0.68 |
| | | | | | | | 200 | 0.87 | 0.85 | 0.91 | 0.85 |
| | | | | | | | 300 | 0.91 | 0.88 | 0.95 | 0.91 |
| | | | | | | | 400 | 0.92 | 0.91 | 0.96 | 0.93 |
| | | | | | | | 1000 | 0.93 | 0.92 | 0.97 | 0.98 |
| $\Theta_2$ | 1HP2 | H1:T0g0 | 0.5 | 0 | 0.5 | 0 | 50 | 0.17 | 0.48 | 0.19 | 0.53 |
| | | | | | | | 100 | 0.48 | 0.72 | 0.53 | 0.79 |
| | | | | | | | 200 | 0.83 | 0.83 | 0.91 | 0.91 |
| | | | | | | | 300 | 0.90 | 0.85 | 0.97 | 0.95 |
| | | | | | | | 400 | 0.94 | 0.86 | 0.98 | 0.97 |
| | | | | | | | 1000 | 0.95 | 0.87 | 0.97 | 0.99 |

| Group | Model | Alias | $T_1$ | $T_3$ | $\gamma_1$ | $\gamma_3$ | n | $\chi2$ reverse | $\chi2$ stepwise | eCDF reverse | eCDF stepwise |
|---|---|---|---|---|---|---|---|---|---|---|---|
| $\Theta_2$ | 1HP3 | H1:T0g1 | 0.5 | 0 | 0.5 | 1 | 50 | 0.20 | 0.32 | 0.22 | 0.35 |
| | | | | | | | 100 | 0.49 | 0.61 | 0.54 | 0.67 |
| | | | | | | | 200 | 0.86 | 0.86 | 0.87 | 0.73 |
| | | | | | | | 300 | 0.81 | 0.83 | 0.86 | 0.74 |
| | | | | | | | 400 | 0.94 | 0.89 | 0.93 | 0.89 |
| | | | | | | | 1000 | 0.94 | 0.88 | 0.95 | 0.94 |
| $\Theta_2$ | T1 | H1:TT10 | 0.5 | 0.5 | 1 | 0 | 50 | 0.42 | 0.50 | 0.46 | 0.55 |
| | | | | | | | 100 | 0.77 | 0.73 | 0.85 | 0.80 |
| | | | | | | | 200 | 0.88 | 0.83 | 0.97 | 0.91 |
| | | | | | | | 300 | 0.92 | 0.87 | 0.96 | 0.94 |
| | | | | | | | 400 | 0.92 | 0.85 | 0.97 | 0.89 |
| | | | | | | | 1000 | 0.92 | 0.84 | 1.00 | 0.91 |
| $\Theta_2$ | T2 | H1:TT01 | 0.5 | 0.5 | 0 | 1 | 50 | 0.62 | 0.68 | 0.68 | 0.75 |
| | | | | | | | 100 | 0.89 | 0.85 | 0.98 | 0.94 |
| | | | | | | | 200 | 0.92 | 0.87 | 0.97 | 0.96 |
| | | | | | | | 300 | 0.94 | 0.87 | 0.97 | 0.95 |
| | | | | | | | 400 | 0.92 | 0.86 | 0.97 | 0.90 |
| | | | | | | | 1000 | 0.93 | 0.88 | 1.00 | 0.96 |
| $\Theta_3$ | 1H1 | H1:TTg0 | 0.5 | 0.5 | 0.5 | 0 | 50 | 0.01 | 0.02 | 0.01 | 0.02 |
| | | | | | | | 100 | 0.16 | 0.27 | 0.18 | 0.30 |
| | | | | | | | 200 | 0.66 | 0.73 | 0.67 | 0.75 |
| | | | | | | | 300 | 0.81 | 0.86 | 0.88 | 0.76 |
| | | | | | | | 400 | 0.88 | 0.89 | 0.97 | 0.95 |
| | | | | | | | 1000 | 0.91 | 0.91 | 0.96 | 0.96 |
| $\Theta_3$ | 1H2 | H1:TT1g | 0.5 | 0.5 | 1 | 0.5 | 50 | 0.32 | 0.48 | 0.35 | 0.53 |
| | | | | | | | 100 | 0.79 | 0.85 | 0.87 | 0.88 |
| | | | | | | | 200 | 0.95 | 0.96 | 0.96 | 0.93 |
| | | | | | | | 300 | 0.95 | 0.95 | 0.97 | 0.94 |
| | | | | | | | 400 | 0.97 | 0.97 | 0.98 | 0.95 |
| | | | | | | | 1000 | 0.96 | 0.96 | 0.98 | 0.96 |
| $\Theta_3$ | 1H3 | H1:TT0g | 0.5 | 0.5 | 0 | 0.5 | 50 | 0.08 | 0.15 | 0.09 | 0.17 |
| | | | | | | | 100 | 0.30 | 0.42 | 0.33 | 0.46 |
| | | | | | | | 200 | 0.68 | 0.75 | 0.75 | 0.75 |
| | | | | | | | 300 | 0.86 | 0.90 | 0.89 | 0.77 |
| | | | | | | | 400 | 0.93 | 0.94 | 0.96 | 0.88 |
| | | | | | | | 1000 | 0.95 | 0.95 | 0.99 | 0.95 |
| $\Theta_3$ | 1H4 | H1:TTg1 | 0.5 | 0.5 | 0.5 | 1 | 50 | 0.14 | 0.20 | 0.15 | 0.22 |
| | | | | | | | 100 | 0.38 | 0.48 | 0.42 | 0.53 |
| | | | | | | | 200 | 0.76 | 0.81 | 0.84 | 0.81 |
| | | | | | | | 300 | 0.89 | 0.89 | 0.93 | 0.86 |
| | | | | | | | 400 | 0.90 | 0.90 | 0.95 | 0.90 |
| | | | | | | | 1000 | 0.93 | 0.93 | 0.97 | 0.95 |
| $\Theta_3$ | 2PH | H2:T0gg | 0.5 | 0 | 0.25 | 0.5 | 50 | 0.18 | 0.33 | 0.35 | 0.28 |
| | | | | | | | 100 | 0.52 | 0.64 | 0.56 | 0.31 |
| | | | | | | | 200 | 0.56 | 0.84 | 0.66 | 0.34 |
| | | | | | | | 300 | 0.51 | 0.62 | 0.67 | 0.38 |
| | | | | | | | 400 | 0.93 | 0.89 | 0.88 | 0.45 |
| | | | | | | | 1000 | 0.93 | 0.88 | 0.94 | 0.68 |

| Group | Model | Alias | $T_1$ | $T_3$ | $\gamma_1$ | $\gamma_3$ | n | $\chi 2$ reverse | $\chi 2$ stepwise | eCDF reverse | eCDF stepwise |
|---|---|---|---|---|---|---|---|---|---|---|---|
| $\Theta_4$ | 2H1 | H1:TTgg | 0.5 | 0.5 | 0.5 | 0.5 | 50 | 0.08 | 0.09 | 0.11 | 0.07 |
| | | | | | | | 100 | 0.26 | 0.28 | 0.29 | 0.26 |
| | | | | | | | 200 | 0.70 | 0.71 | 0.73 | 0.69 |
| | | | | | | | 300 | 0.91 | 0.92 | 0.96 | 0.76 |
| | | | | | | | 400 | 0.97 | 0.97 | 0.98 | 0.88 |
| | | | | | | | 1000 | 1.00 | 1.00 | 1.00 | 0.97 |
| $\Theta_4$ | 2H2 | H2:TTgg | 0.5 | 0.5 | 0.5 | 0.5 | 50 | 0.02 | 0.05 | 0.12 | 0.09 |
| | | | | | | | 100 | 0.24 | 0.31 | 0.33 | 0.35 |
| | | | | | | | 200 | 0.74 | 0.78 | 0.81 | 0.75 |
| | | | | | | | 300 | 0.94 | 0.94 | 0.97 | 0.80 |
| | | | | | | | 400 | 0.98 | 0.99 | 0.99 | 0.95 |
| | | | | | | | 1000 | 1.00 | 1.00 | 1.00 | 0.99 |
