## Supplementary tables (non-conflicting patterns) for "A 4-lineage statistical suite to evaluate the support of large-scale retrotransposon insertion data to reconstruct evolutionary trees"

**Table S1.**

**Table S1. Simple conflict-free patterns.** The first column shows non-zero value(s) in y-notation. The second column contains the respective presence/absence patterns. The third column indicates model names. The “Tree” column presents the exact tree model and species succession. The last column shows the resulting order of species converted to A,B,C, and D with an initial order A,B,C,D corresponding to 1234. In the case of model T1, the two lines in “Tree” and “order of species” indicate the condition where the number of markers related to the first pattern is larger than the second one; the second line indicates the opposite situation.

| Non-zero value positions ($y_{ij}$) | Presence/absence Pattern | Model | Tree | Order of species |
|---|---|---|---|---|
| $y_{11}$ | -+++ | TP | H1:T00n:1234 | A,B,C,D |
| $y_{12}$ | --++ | PT | H1:0Tn1:1234 | A,B,C,D |
| $y_{13}$ | -+-+ | PT | H1:0Tn1:1324 | A,C,B,D |
| $y_{14}$ | -++- | PT | H1:0Tn1:1432 | A,D,B,C |
| $y_{22}$ | +-++ | TP | H1:T00n: 2134 | B,A,C,D |
| $y_{23}$ | +--+ | PT | H1:0Tn1:2314 | B,C,A,D |
| $y_{24}$ | +-+- | PT | H1:0Tn1:2413 | B,D,A,C |
| $y_{33}$ | ++-+ | TP | H1:T00n: 3124 | C,A,B,D |
| $y_{34}$ | ++-- | PT | H1:0Tn1:3412 | C,D,A,B |
| $y_{44}$ | +++- | TP | H1:T00n: 4123 | D,A,B,C |
| $y_{12}$ and $y_{34}$ | --++<br>++-- | T1 | H1:TT10:1324<br>H1:TT10:3142 | A,C,B,D<br>C,A,D,B |
| $y_{13}$ and $y_{24}$ | -+-+<br>+-+- | T1 | H1:TT10:1234<br>H1:TT10:2143 | A,B,C,D<br>B,A,D,C |
| $y_{14}$ and $y_{23}$ | +--+<br>-++- | T1 | H1:TT10:1243<br>H1:TT10:2134 | A,B,D,C<br>B,A,C,D |
| $y_{11}$ and $y_{12}$ | -+++ and --++ | T2 | H1:TT01:1234 | A,B,C,D |
| $y_{11}$ and $y_{13}$ | -+++ and -+-+ | T2 | H1:TT01:1324 | A,C,B,D |
| $y_{11}$ and $y_{14}$ | -+++and -++- | T2 | H1:TT01:1423 | A,D,B,C |
| $y_{22}$ and $y_{12}$ | +-++ and --++ | T2 | H1:TT01:2134 | B,A,C,D |
| $y_{22}$ and $y_{23}$ | +-++ and +--+ | T2 | H1:TT01:2314 | B,C,A,D |
| $y_{22}$ and $y_{24}$ | +-++ and +-+- | T2 | H1:TT01:2413 | B,D,A,C |
| $y_{33}$ and $y_{13}$ | ++-+ and -+-+ | T2 | H1:TT01:3124 | C,A,B,D |
| $y_{33}$ and $y_{23}$ | ++-+ and +--+ | T2 | H1:TT01:3214 | C,B,A,D |
| $y_{33}$ and $y_{34}$ | ++-+ and ++-- | T2 | H1:TT01:3412 | C,D,A,B |
| $y_{44}$ and $y_{14}$ | +++- and -++- | T2 | H1:TT01:4123 | D,A,B,C |
| $y_{44}$ and $y_{24}$ | +++- and +-+- | T2 | H1:TT01:4213 | D,B,A,C |
| $y_{44}$ and $y_{34}$ | +++- and ++-- | T2 | H1:TT01:4312 | D,C,A,B |

**Table S2. Formulas for generating data for KKSC comparisons of resolved tree topologies (without hybridization).** The first column shows the exact tree model and species succession, and can be derived from the likelihood result table. The second column presents the respective model names. The last two columns present algorithm for calculating the three values $Y_1$, $Y_2$, and $Y_3$ for KKSC comparisons.

| Tree | Model | $Y_1$, $Y_2$, and $Y_3$ (first comparison) | $Y_1$, $Y_2$, and $Y_3$ (second comparison) |
|---|---|---|---|
| H1:T00n:1234 | TP | $Y_1=y_{11}+y_{12}$; $Y_2=y_{23}+y_{33}$; $Y_3=y_{24}+y_{44}$. | |
| H1:0Tn1:1234 | PT | $Y_1=y_{12}+y_{22}$; $Y_2=y_{13}+y_{33}$; $Y_3=y_{14}+y_{44}$. | |
| H1:0Tn1:1324 | PT | $Y_1=y_{13}+y_{33}$; $Y_2=y_{12}+y_{22}$; $Y_3=y_{14}+y_{44}$. | |
| H1:0Tn1:1423 | PT | $Y_1=y_{14}+y_{44}$; $Y_2=y_{12}+y_{22}$; $Y_3=y_{13}+y_{33}$. | |
| H1:T00n:2134 | TP | $Y_1=y_{12}+y_{22}$; $Y_2=y_{13}+y_{33}$; $Y_3=y_{14}+y_{44}$. | |
| H1:0Tn1:2314 | PT | $Y_1=y_{23}+y_{33}$; $Y_2=y_{11}+y_{12}$; $Y_3=y_{24}+y_{44}$. | |
| H1:0Tn1:2413 | PT | $Y_1=y_{24}+y_{44}$; $Y_2=y_{11}+y_{12}$; $Y_3=y_{23}+y_{33}$. | |
| H1:T00n: 3124 | TP | $Y_1=y_{13}+y_{33}$; $Y_2=y_{12}+y_{22}$; $Y_3=y_{14}+y_{44}$. | |
| H1:0Tn1:3412 | PT | $Y_1=y_{34}+y_{44}$; $Y_2=y_{22}+y_{23}$; $Y_3=y_{11}+y_{13}$. | |
| H1:T00n: 4123 | TP | $Y_1=y_{14}+y_{44}$; $Y_2=y_{12}+y_{22}$; $Y_3=y_{13}+y_{33}$. | |
| H1:TT10:1324 | T1 | $Y_1=y_{34}+y_{44}$; $Y_2=y_{11}+y_{13}$; $Y_3=y_{22}+y_{23}$. | $Y_1=y_{12}+y_{22}$; $Y_2=y_{13}+y_{33}$; $Y_3=y_{14}+y_{44}$. |
| H1:TT10:3142 | T1 | $Y_1=y_{12}+y_{22}$; $Y_2=y_{13}+y_{33}$; $Y_3=y_{14}+y_{44}$. | $Y_1=y_{34}+y_{44}$; $Y_2=y_{11}+y_{13}$; $Y_3=y_{22}+y_{23}$. |
| H1:TT10:1234 | T1 | $Y_1=y_{24}+y_{44}$; $Y_2=y_{11}+y_{12}$; $Y_3=y_{23}+y_{33}$. | $Y_1=y_{13}+y_{33}$; $Y_2=y_{12}+y_{22}$; $Y_3=y_{14}+y_{44}$. |
| H1:TT10:2143 | T1 | $Y_1=y_{13}+y_{33}$; $Y_2=y_{12}+y_{22}$; $Y_3=y_{14}+y_{44}$. | $Y_1=y_{24}+y_{44}$; $Y_2=y_{11}+y_{12}$; $Y_3=y_{23}+y_{33}$. |
| H1:TT10:1243 | T1 | $Y_1=y_{23}+y_{33}$; $Y_2=y_{11}+y_{12}$; $Y_3=y_{24}+y_{44}$. | $Y_1=y_{14}+y_{44}$; $Y_2=y_{12}+y_{22}$; $Y_3=y_{13}+y_{33}$. |
| H1:TT10:2134 | T1 | $Y_1=y_{14}+y_{44}$; $Y_2=y_{12}+y_{22}$; $Y_3=y_{13}+y_{33}$. | $Y_1=y_{23}+y_{33}$; $Y_2=y_{11}+y_{12}$; $Y_3=y_{24}+y_{44}$. |
| H1:TT01:1234 | T2 | $Y_1=y_{11}+y_{13}$; $Y_2=y_{22}+y_{23}$; $Y_3=y_{34}+y_{44}$. | $Y_1=y_{12}+y_{22}$; $Y_2=y_{13}+y_{33}$; $Y_3=y_{14}+y_{44}$. |
| H1:TT01:1324 | T2 | $Y_1=y_{34}+y_{44}$; $Y_2=y_{11}+y_{13}$; $Y_3=y_{22}+y_{23}$. | $Y_1=y_{12}+y_{22}$; $Y_2=y_{13}+y_{33}$; $Y_3=y_{14}+y_{44}$. |
| H1:TT01:1423 | T2 | $Y_1=y_{11}+y_{12}$; $Y_2=y_{24}+y_{44}$; $Y_3=y_{23}+y_{33}$. | $Y_1=y_{14}+y_{44}$; $Y_2=y_{12}+y_{22}$; $Y_3=y_{13}+y_{33}$. |
| H1:TT01:2134 | T2 | $Y_1=y_{22}+y_{23}$; $Y_2=y_{11}+y_{13}$; $Y_3=Y_{34}+Y_{44}$. | $Y_1=y_{11}+y_{12}$; $Y_2=y_{23}+y_{33}$; $Y_3=y_{24}+y_{44}$. |
| H1:TT01:2314 | T2 | $Y_1=y_{12}+y_{22}$; $Y_2=y_{13}+y_{33}$; $Y_{3'}=y_{14}+y_{44}$. | $Y_1=y_{23}+y_{33}$; $Y_2=y_{11}+y_{12}$; $Y_3=y_{24}+y_{44}$. |
| H1:TT01:2413 | T2 | $Y_1=y_{12}+y_{22}$; $Y_2=y_{14}+y_{44}$; $Y_3=Y_{13}+Y_{33}$. | $Y_1=y_{24}+y_{44}$; $Y_2=y_{23}+y_{33}$; $Y_3=y_{11}+y_{12}$. |
| H1:TT01:3124 | T2 | $Y_1=y_{23}+y_{33}$; $Y_2=y_{11}+y_{12}$; $Y_3=y_{24}+y_{44}$. | $Y_1=y_{11}+y_{13}$; $Y_2=y_{22}+y_{23}$; $Y_3=y_{34}+y_{44}$. |
| H1:TT01:3214 | T2 | $Y_1=y_{13}+y_{33}$; $Y_2=y_{12}+y_{22}$; $Y_3=y_{14}+y_{44}$. | $Y_1=y_{22}+y_{23}$; $Y_2=y_{11}+y_{13}$; $Y_3=y_{34}+y_{44}$. |
| H1:TT01:3412 | T2 | $Y_1=y_{13}+y_{33}$; $Y_2=y_{12}+y_{22}$; $Y_3=y_{14}+y_{44}$. | $Y_1=y_{34}+y_{44}$; $Y_2=y_{23}+y_{33}$; $Y_3=y_{11}+y_{13}$. |
| H1:TT01:4123 | T2 | $Y_1=y_{24}+y_{44}$; $Y_2=y_{11}+y_{12}$; $Y_3=y_{23}+y_{33}$. | $Y_1=y_{11}+y_{14}$; $Y_2=y_{22}+y_{24}$; $Y_3=y_{33}+y_{34}$. |
| H1:TT01:4213 | T2 | $Y_1=y_{14}+y_{44}$; $Y_2=y_{12}+y_{22}$; $Y_3=y_{13}+y_{33}$. | $Y_1=y_{24}+y_{44}$; $Y_2=y_{11}+y_{14}$; $Y_3=y_{33}+y_{34}$. |
| H1:TT01:4312 | T2 | $Y_1=y_{14}+y_{44}$; $Y_3=y_{13}+y_{33}$; $Y_{3'}=y_{12}+y_{22}$. | $Y_1=y_{33}+y_{34}$; $Y_2=y_{11}+y_{14}$; $Y_3=y_{22}+y_{24}$. |
