## Supplementary material for "A 4-lineage statistical suite to evaluate the support of large-scale retrotransposon insertion data to reconstruct evolutionary trees": Sources of genomes used and RepeatMasker reports.

### Great ape phylogenetic project

***Homo sapiens***

Genome: http://hgdownload.soe.ucsc.edu/goldenPath/hg38/bigZips/hg38.chromFa.tar.gz

RepeatMasker report: http://hgdownload.soe.ucsc.edu/goldenPath/hg38/bigZips/hg38.fa.out.gz

***Pan troglodytes***

Genome:
ftp://ftp.ncbi.nlm.nih.gov/genomes/all/GCA/002/880/755/GCA_002880755.3_Clint_PTRv2/GCA_002880755.3_Clint_PTRv2_genomic.fna.gz

RepeatMasker report:
ftp://ftp.ncbi.nlm.nih.gov/genomes/all/GCA/002/880/755/GCA_002880755.3_Clint_PTRv2/GCA_002880755.3_Clint_PTRv2_rm.out.gz

***Pan paniscus***

Genome:
ftp://ftp.ncbi.nlm.nih.gov/genomes/all/GCA/000/258/655/GCA_000258655.2_panpan1.1/GCA_000258655.2_panpan1.1_genomic.fna.gz

RepeatMasker report:
ftp://ftp.ncbi.nlm.nih.gov/genomes/all/GCA/000/258/655/GCA_000258655.2_panpan1.1/GCA_000258655.2_panpan1.1_rm.out.gz

***Gorilla gorilla***

Genome:
ftp://ftp.ncbi.nlm.nih.gov/genomes/all/GCA/000/151/905/GCA_000151905.3_gorGor4/GCA_000151905.3_gorGor4_genomic.fna.gz

RepeatMasker report:
ftp://ftp.ncbi.nlm.nih.gov/genomes/all/GCA/000/151/905/GCA_000151905.3_gorGor4/GCA_000151905.3_gorGor4_rm.out.gz

***Pongo abelii***

Genome:
ftp://ftp.ncbi.nlm.nih.gov/genomes/all/GCF/002/880/775/GCF_002880775.1_Susie_PABv2/GCF_002880775.1_Susie_PABv2_genomic.fna.gz

RepeatMasker report:
ftp://ftp.ncbi.nlm.nih.gov/genomes/all/GCF/002/880/775/GCF_002880775.1_Susie_PABv2/GCF_002880775.1_Susie_PABv2_rm.out.gz

### Dog lineage diversification project

**Boxer**

Genome:

https://ftp.ncbi.nlm.nih.gov/genomes/all/GCA/000/002/285/GCA_000002285.2_CanFam3.1/GCA_000002285.2_CanFam3.1_genomic.fna.gz

RepeatMasker report:

https://ftp.ncbi.nlm.nih.gov/genomes/all/GCA/000/002/285/GCA_000002285.2_CanFam3.1/GCA_000002285.2_CanFam3.1_rm.out.gz

**Beagle**

Genome:

ftp://ftp.ncbi.nlm.nih.gov/genomes/all/GCA/000/331/495/GCA_000331495.1_Beagle/GCA_000331495.1_Beagle_genomic.fna.gz

RepeatMasker report:

ftp://ftp.ncbi.nlm.nih.gov/genomes/all/GCA/000/331/495/GCA_000331495.1_Beagle/GCA_000331495.1_Beagle_rm.out.gz

**Gray Wolf**

Genome:

ftp://ftp.ncbi.nlm.nih.gov/genomes/all/GCA/007/922/845/GCA_007922845.1_UniMelb_Wolf_Refassem_1/GCA_007922845.1_UniMelb_Wolf_Refassem_1_genomic.fna.gz

RepeatMasker report:

ftp://ftp.ncbi.nlm.nih.gov/genomes/all/GCA/007/922/845/GCA_007922845.1_UniMelb_Wolf_Refassem_1/GCA_007922845.1_UniMelb_Wolf_Refassem_1.rm.out.gz

**German shepherd**

Genome:

ftp://ftp.ncbi.nlm.nih.gov/genomes/all/GCA/008/641/055/GCA_008641055.1_ASM864105v1/GCA_008641055.1_ASM864105v1_genomic.fna.gz

RepeatMasker report:

ftp://ftp.ncbi.nlm.nih.gov/genomes/all/GCA/008/641/055/GCA_008641055.1_ASM864105v1/GCA_008641055.1_ASM864105v1_rm.out.gz

### Mouse strains project.

**CBA/J**

Genome:

ftp://ftp.ncbi.nlm.nih.gov/genomes/all/GCA/001/624/475/GCA_001624475.1_CBA_J_v1/GCA_001624475.1_CBA_J_v1_genomic.fna.gz

RepeatMasker report:

ftp://ftp.ncbi.nlm.nih.gov/genomes/all/GCA/001/624/475/GCA_001624475.1_CBA_J_v1/GCA_001624475.1_CBA_J_v1_rm.out.gz

**C57BL/6J**

Genome:

ftp://ftp.ncbi.nlm.nih.gov/genomes/all/GCA/003/774/525/GCA_003774525.2_ASM377452v2/GCA_003774525.2_ASM377452v2_genomic.fna.gz

RepeatMasker report:

ftp://ftp.ncbi.nlm.nih.gov/genomes/all/GCA/003/774/525/GCA_003774525.2_ASM377452v2/GCA_003774525.2_ASM377452v2_rm.out.gz

**BALB/cJ**

Genome:

ftp://ftp.ncbi.nlm.nih.gov/genomes/all/GCA/001/632/525/GCA_001632525.1_BALB_cJ_v1/GCA_001632525.1_BALB_cJ_v1_genomic.fna.gz

RepeatMasker report:

ftp://ftp.ncbi.nlm.nih.gov/genomes/all/GCA/001/632/525/GCA_001632525.1_BALB_cJ_v1/GCA_001632525.1_BALB_cJ_v1_rm.out.gz

**DBA/2J**

Genome:

ftp://ftp.ncbi.nlm.nih.gov/genomes/all/GCA/001/624/505/GCA_001624505.1_DBA_2J_v1/GCA_001624505.1_DBA_2J_v1_genomic.fna.gz

**RepeatMasker report:**

ftp://ftp.ncbi.nlm.nih.gov/genomes/all/GCA/001/624/505/GCA_001624505.1_DBA_2J_v1/GCA_001624505.1_DBA_2J_v1_rm.out.gz
